# Novel insights into pulmonary phosphate homeostasis and osteoclastogenesis emerge from the study of pulmonary alveolar microlithiasis

**DOI:** 10.1101/2021.07.11.451970

**Authors:** Yasuaki Uehara, Nikolaos M. Nikolaidis, Lori B. Pitstick, Huixing Wu, Jane J. Yu, Erik Zhang, Yoshihiro Hasegawa, Yusuke Tanaka, John G. Noel, Jason C. Gardner, Elizabeth J. Kopras, Wendy D. Haffey, Kenneth D. Greis, Jinbang Guo, Jason C. Woods, Kathryn A. Wikenheiser-Brokamp, Shuyang Zhao, Yan Xu, Jennifer E. Kyle, Charles Ansong, Steven L. Teitelbaum, Yoshikazu Inoue, Göksel Altinişik, Francis X. McCormack

## Abstract

Pulmonary alveolar microlithiasis (PAM) is an autosomal recessive lung disease caused by a deficiency in the pulmonary epithelial Npt2b sodium-phosphate co-transporter that results in accumulation of phosphate and formation of hydroxyapatite microliths in the alveolar space. The single cell transcriptomic analysis of a PAM lung explant showing a robust osteoclast gene signature in alveolar monocytes and the finding that calcium phosphate microliths contain a rich protein and lipid matrix that includes bone resorbing osteoclast enzymes suggested a role for osteoclast-like cells in the defense against microliths. While investigating the mechanisms of microlith clearance, we found that Npt2b modulates pulmonary phosphate homeostasis through effects on alternative phosphate transporter activity and alveolar osteoprotegerin, and that microliths induce osteoclast formation and activity in a receptor activator of nuclear factor-κB ligand (RANKL) and dietary phosphate dependent manner. This work reveals that Npt2b and pulmonary osteoclast-like cells play key roles in pulmonary homeostasis and suggest potential new therapeutic targets for the treatment of lung disease.

## Introduction

Pulmonary alveolar microlithiasis (PAM) is an autosomal recessive disorder (OMIM 265100) caused by mutations of the SLC34A2 gene, which results in deficiency of the Npt2b sodium-phosphate co-transporter^1^. As surfactant phospholipids are catabolized by alveolar macrophages, phosphate accumulates in the alveolar space due to loss of the epithelial pump. Calcium phosphate crystals form spontaneously in the lumen of pulmonary alveolar airspaces, resulting in pulmonary fibrosis and pulmonary hypertension. Over 1000 PAM cases have been reported, mostly from Japan, Turkey, and Italy. The disease is slowly progressive, with most patients living into middle age, but lethality is reported in all age groups including infants^2, 3^. There are currently no effective therapies for PAM. Therapeutic whole lung lavage and etidronate therapy have been attempted with mixed and generally disappointing results^4–9^, and current management options are limited to supportive care or lung transplantation.

The role of microliths in the pathogenesis of PAM have not been well studied. Barnard reported that microliths are composed of calcium and phosphate in ratios that are consistent with hydroxyapatite^10^. In our previous study, we reported that dietary phosphate restriction decreased alveolar microlith accumulation in a PAM animal model, but the mechanisms involved remained unclear^11^. In thus study, we found upregulation of multiple osteoclast genes and proteins in PAM lung, as well as in the bronchoalveolar lavage cells and the pulmonary parenchyma of Npt2b^-/-^ mice, including those for signature osteoclast proteins tartrate-resistant acid phosphatase (TRAP), ATP6v0d2, calcitonin receptor (CALCR) and cathepsin K (CTSK). Intratracheal adoptive transfer of hydroxyapatite microliths into Npt2b^+/+^ mice induced bone resorbing capacity in bronchoalveolar lavage cells isolated one week later. Dietary phosphate restriction augmented osteoclast-related gene expression in the lung, and enhanced clearance of microliths through receptor activator of nuclear factor-κB ligand (RANKL)-dependent osteoclast-like multinucleated giant cell differentiation. High dietary phosphate levels increased microlith burden in the Npt2b^-/-^ lung by elevating levels of phosphate and osteoprotegerin, the soluble RANKL decoy receptor, in the alveolar space. Npt2b^+/+^ mice were able to maintain phosphate balance in the alveolar space through a range of phosphate concentrations. These studies reveal previously unrecognized roles for Npt2b in pulmonary phosphate homeostasis and for osteoclast-like cells in response to particulate challenge in the lung, both of which have broader translational implications.

## Results

### Scanning electron microscopy and elemental, proteomic and lipidomic analysis of human and mouse PAM microliths

To better understand the pathogenesis of PAM, we first analyzed the composition of microliths isolated from the lungs of Npt2b^-/-^ mice and a PAM patient. Scanning electron microscopy (SEM) imaging revealed spherical particulate microliths with a hairy, fibrous coating, particularly for the human stones, and human and mouse microlith diameters ranged between 100 to 500 μm (Fig. 1a, b) and 5 to 20 μm (Fig. 1c, d), respectively. The sodium dodecyl sulfate (SDS) washed mouse microliths had a porous, spongy, cortical bone-like surface (Fig. 1e-g) that was similar to if somewhat less distinct than that of the synthetic hydroxyapatite spheres (Fig. 1h, i). Elemental analysis of the microliths by energy dispersive spectroscopy identified calcium and phosphorus in ratios that were consistent with hydroxyapatite, but slightly lower than that that of synthetic hydroxyapatite crystals, due to incorporation of proteins and/or lipids into the matrix of the native stones (Supplementary Table 1). Proteomic analyses of the microliths identified over 300 proteins including those commonly found in renal stones, such as alpha-2HS-glycoprotein, prelamin-A/C and annexin A2, and in the alveolar lumen, such as surfactant proteins A, B, C and D (Fig. 1j, Supplementary Table 2 and 3). Osteoclast proteins lysosomal protease, CTSK and TRAP were also detected in the mouse microliths. Lipidomic analyses revealed the presence of surfactant phospholipids, in proportions that were roughly commensurate with their abundance in pulmonary surfactant (Fig. 1k, Supplementary Table 4).

**Fig. 1.**
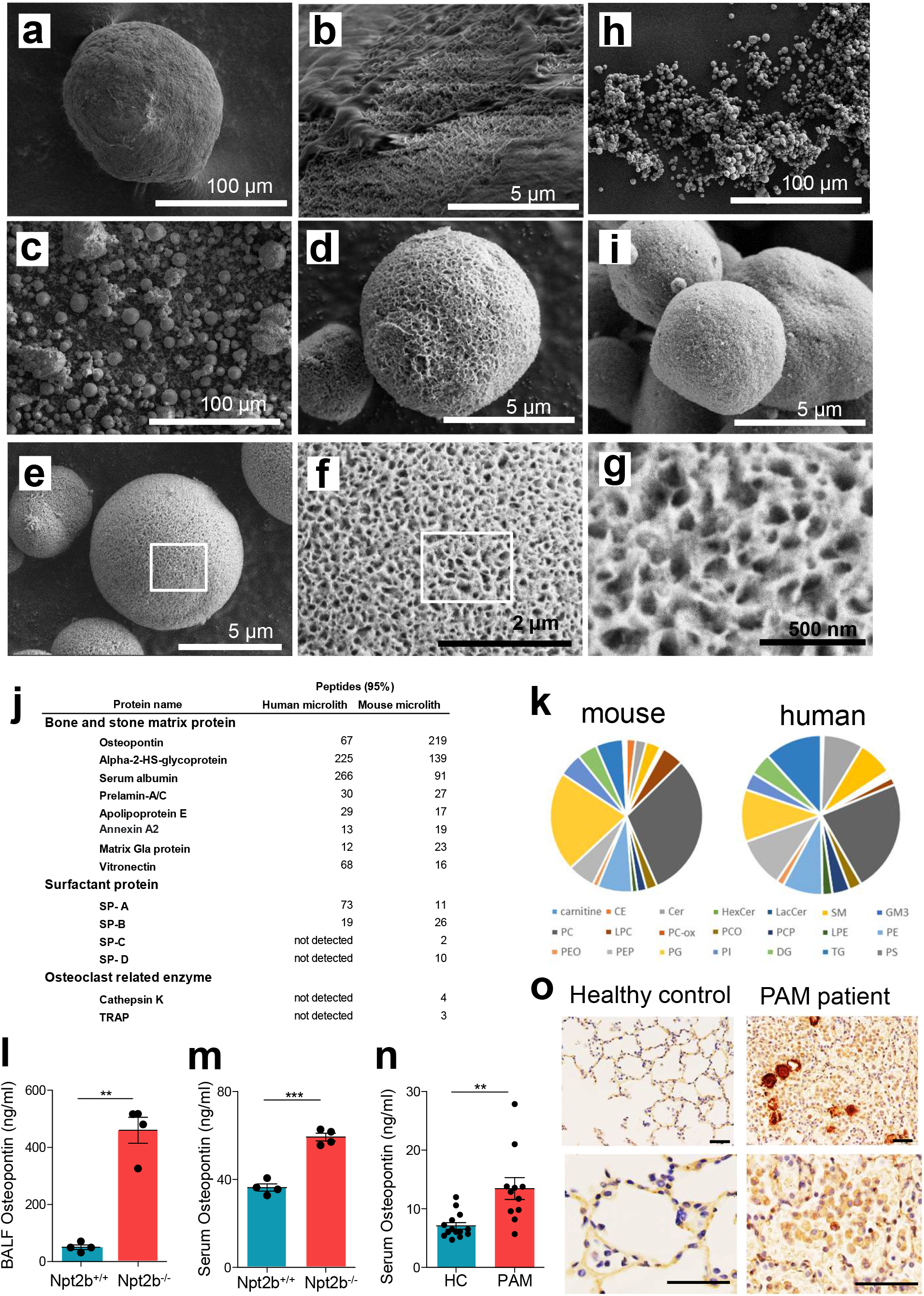
Microliths are composed of hydroxyapatite, have a porous bony surface, and contain bone matrix proteins, and alveolar components, surfactant proteins and phospholipids. **a-i**, Scanning electron microscopy (SEM) of microliths isolated from Npt2b^-/-^ mice (**a, b**), from a human infant lung explant (**c, d**) and commercially available hydroxyapatite spheres (**h, i**). Microliths isolated from Npt2b^-/-^ mice washed with 0.02% SDS at low, medium and high power (**e-g**). **j, k**, Proteins (**j**) and lipids (**k**) detected in microliths from Npt2b^-/-^ mice and PAM patient respectively. **l-n**, Osteopontin quantified by ELISA in mouse BAL fluid (l) and serum (n = 4) (**m**) and serum from PAM patients (PAM) (n=14) and healthy controls (HC) (n=11) (n). **o**, IHC for osteopontin in human control and PAM human infant. Bold scale bar, 50 μm. Data are expressed as means ± SD. **p< 0.01 and ***p<0.001. SP-A(-B, -C, -D), surfactant protein A (B, C, D, respectively); TRAP, tartrate-resistant acid phosphatase; PC, phosphatidylcholine; PE, phosphatidylethanolamine; PG, phosphatidylglycerol; PI, phosphatidylinositol; PS, phosphatidylserine; SM, sphingomyelin; PEO, 1-o-alkyl-2-acyl-PE; CE, cholesterol ester; LPC, lysophosphatidylcholine; PEP, PE plasmalogen; Cer, ceramide; PC-ox, oxidized PC; Hex-Cer, hexosylceramide; PCO, 1-o-alkyl-2-acyl-PC; Lac-Cer, lactosylceramide; PCP, PC plasmalogen; DG, diacylglycerol; LPE, lysophosphatidylethnolamine; TG, triglyceride; GM3, monosialodihexosylganglioside;.

### Osteopontin expression is increased in PAM lung

To evaluate osteopontin (OPN) expression in PAM lung and its potential as a biomarker, OPN levels in bronchoalveolar lavage (BAL) fluid (BALF) and serum were determined. OPN levels in BALF were significantly higher in Npt2b^-/-^ mice than Npt2b^+/+^ mice (Fig. 1l). Furthermore, serum OPN levels in PAM mice and the PAM patients were significantly elevated compared to healthy controls (Fig. 1m, n). Immunohistochemical staining demonstrated high OPN expression localized predominantly in alveolar epithelial type I cells, alveolar epithelial type II cells (AECII) and monocyte/macrophages present around the microliths in the PAM patient lung (Fig. 1o). In contrast, only faint OPN staining was present in alveolar macrophages (AM) and airway epithelium of control lungs.

### Osteoclast related gene signatures are upregulated in PAM lung

The finding that bone degrading enzymes produced by osteoclasts (such as TRAP and CTSK) were incorporated into the matrix of microliths led us to consider a potential role for osteoclast-like functions in microlith clearance in the PAM lung. We employed single-cell RNA-seq analysis on a lung explant from a 2 year old PAM infant and a control lung from a 2 year old infant (Fig. 2a). A heatmap illustrates upregulation of expression of key osteoclast related genes in PAM lung macrophages (Fig. 2b-e) including: a) RANKL mediated osteoclastogenesis signaling genes; GRB2 associated binding protein 2 (GAB2), nuclear factor of activated T cells 1 (NFATC1), melanocyte inducing transcription factor (MITF), TNF receptor associated factor 6 (TRAF6) and inhibitor of nuclear factor kappa B kinase subunit beta (IKBKB), b) osteoclast cell fusion related genes; dendrocyte expressed seven transmembrane protein (DCSTAMP) and osteoclast stimulatory transmembrane protein (OCSTAMP), c) osteoclast precursor specific genes; tartrate-resistant acid phosphatase (ACP5), transcription factor EC (TFEC), and integrin subunit beta 3 (ITGB3), and d) osteoclast receptors, enzymes and acid secretion genes; CALCR, ATPase H^+^ transporting V0 subunit d2 (ATP6V0D2), CTSK, and matrix metallopeptidase 9 (MMP9). On the other hand, EMR1, an AM marker, was downregulated in the PAM myeloid cells (Fig. 2e). These results are consistent with robust osteoclast differentiation in AM of the PAM lung.

**Fig. 2.**
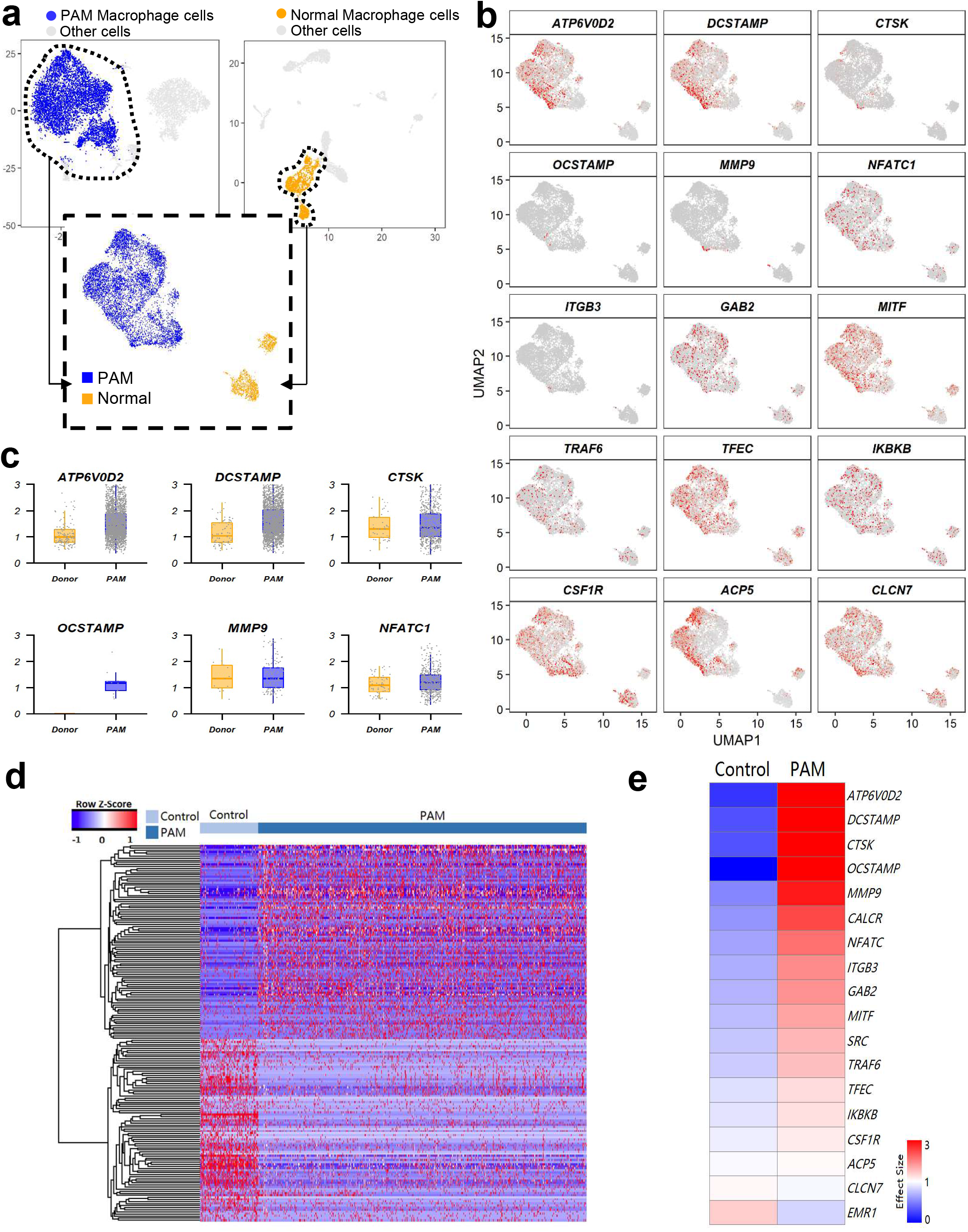
Single-cell RNA sequencing reveals upregulation of osteoclast genes in lung macrophages from a PAM patient. **a**, 2,534 macrophage cells from PAM sample and 1,831 macrophage cells from normal control sample were extracted and re-clustered. Cells are visualized by Uniform Manifold Approximation and Projection (UMAP) and colored by condition. **b**, Expression of osteoclast related genes is shown in UMAP. The expression is z-score normalized. **c**, Box-plots show the expression of selected osteoclast marker genes in PAM macrophages and normal control macrophages. Groups are color coded. The expression is normalized using default normalization method from Seurat. **d**, Heatmap represents the differentially expressed genes. Differentially expressed genes were calculated using Binomial based test (Shekhar et al., 2016). FDR < 0.1, effective size > 2, and the expression frequency > 20% were considered as significant. Hierarchical clustering of genes were performed using complete linkage and Pearson correlation-based distance measurement. Gene expression was z-score normalized. e, Heatmap represents the effective size (defined as the ratio of the % of cells in which the marker was detected between the two conditions) of osteoclast related genes in PAM and control donor lung macrophage cells.

### Co-localization of TRAP and cathepsin K expressing multinucleated giant cells and microliths

We explored osteoclast gene expression in BAL cells from Npt2b^-/-^ mice, and found upregulation of Acp5, Mmp9, Ctsk, Itgb3 and Calcr (Fig. 3a-e). In addition, the gene expression of colony stimulating factor-1 (Csf1), a cytokine important for osteoclast differentiation, was upregulated in Npt2b^-/-^ mice BAL cells (Fig. 3f). TRAP, CTSK and CALCR positive MNGCs were found in Npt2b^-/-^ mouse lungs and the PAM patient lung, but not in control lungs, and only faint TRAP, CTSK CALCR staining were found in AMs of healthy human lung and Npt2b^+/+^ mice (Fig. 3g-r). Interestingly, the airway epithelium also stained positively for CALCR (Fig. 3o, q). Many MNGCs colocalized with microliths and were often found adherent to microliths. The number of nuclei per MNGC and the size of MNGCs in human PAM lung tended to be greater than mouse MNGCs.

**Fig. 3.**
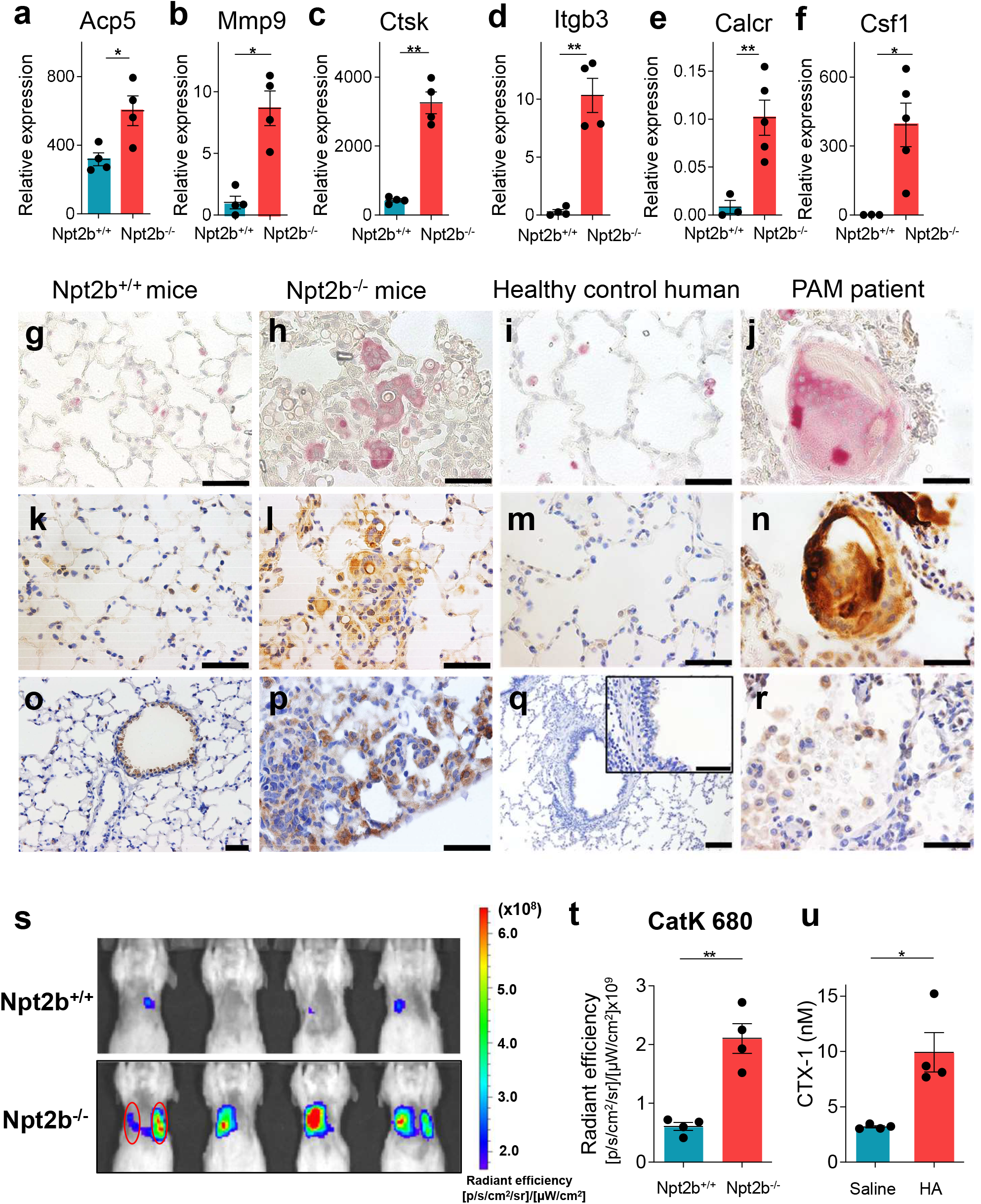
Genetic, immunohistochemical, enzymatic and functional evidence of osteoclast activity in PAM lung. **a-f**, Relative expression of osteoclast-related genes in the BAL cells from 8- to 10-week-old Npt2b^+/+^ and Npt2b^-/-^ mice were determined by RT-PCR. n = 3 to 5 mice per group. Lung sections of 8- to 10-week-old Npt2b^+/+^ (**g, k, o**) and Npt2b^-/-^ mice (**h, l, p**), and a healthy control human (**i, m, q**) and a PAM patient (**j, n, r**) were fixed and stained for TRAP (**g-j**), CTSK (**k-n**) and CALCR (**o-r**). Bold scale bar, 50 μm. **s-t**, Npt2b^+/+^ and Npt2b^-/-^ mice were injected with the cleavage activated cathepsin K substrate, CatK 680 FAST, and fluorescent images were acquired 18 hours later by IVIS (s), and the fluorescent signal was quantified (**t**). **u**. BAL cells isolated from C57BL/6J mice at day 14 post intratracheal challenge with saline (Saline) or hydroxyapatite (HA) were plated on bone bovine slices with M-CSF and RANKL. The levels of CTX-I in bone culture medium were measured by ELISA (N=4 wells per group). Data are expressed as means ± SD. *p<0.05 and **p<0.01.

### Osteoclastic activities were upregulated in the lungs of Npt2b^-/-^ mice lung and Npt2b^+/+^ recipients of adoptively transferred hydroxyapatite microliths

CTSK activity was assessed using the in vivo imaging system (IVIS) with Cat K 680 (Cat K), a cleavage activated probe. Fluorescence intensity of Cat K was increased in the lungs of Npt2b^-/-^ mice lung compared to Npt2b^+/+^ mice at 18 h post intratracheal Cat K injection (Fig. 3s, t). These results demonstrate that upregulation of an enzymatically active signature osteoclast protease (i.e CTSK) in the lungs of Npt2b^-/-^ mice.

A bone resorption assay was used to determine if microliths induce this signature osteoclast function in AM. BAL cells isolated 14d after intratracheal challenge with saline or hydroxyapatite spheres were plated on bovine bone slices, and the release of the bovine type I collagen C-telopeptide fragment, CTX-1, into the media was measured. We found the CTX-1 levels produced by the BAL cells from hydroxyapatite treated mice were twice those of saline treated mice. (Fig. 3u). This result indicates that hydroxyapatite causes differentiation of AM into mature lung osteoclasts.

### Microlith clearance is delayed in CCR2^-/-^ mice

Adoptive transfer of microliths into wild-type (WT) lungs induced Acp5, Mmp9, Ctsk and Itgb3 genes expression in BAL cells at day 7 post challenge (Fig. 4a-d). TRAP positive monocytes and macrophages engulfed microliths and formed inflammatory aggregates around microliths at day 7 post adoptive transfer, and both microliths and inflammatory changes were markedly diminished to absent by day 28 (Fig. 4e, left panel). To determine the role of tissue resident AM vs. monocyte derived macrophages in microlith clearance, we repeated the experiment in CCR2^-/-^ mice which cannot recruit monocytes because the cells fail to exit the marrow^12^. After adoptive transfer of microliths into CCR2^-/-^ mice, TRAP expression in inflammatory sites was diminished compared to CCR2^+/+^ mice, and although inflammation was largely resolved in both models by day 28, microlith clearance was delayed in CCR2^-/-^ mice compared to WT mice (Fig. 4e, right panel). These results indicate that microliths induce osteoclastic transformation of BAL cells, and monocyte recruitment is required for optimal clearance of stones.

**Fig. 4.**
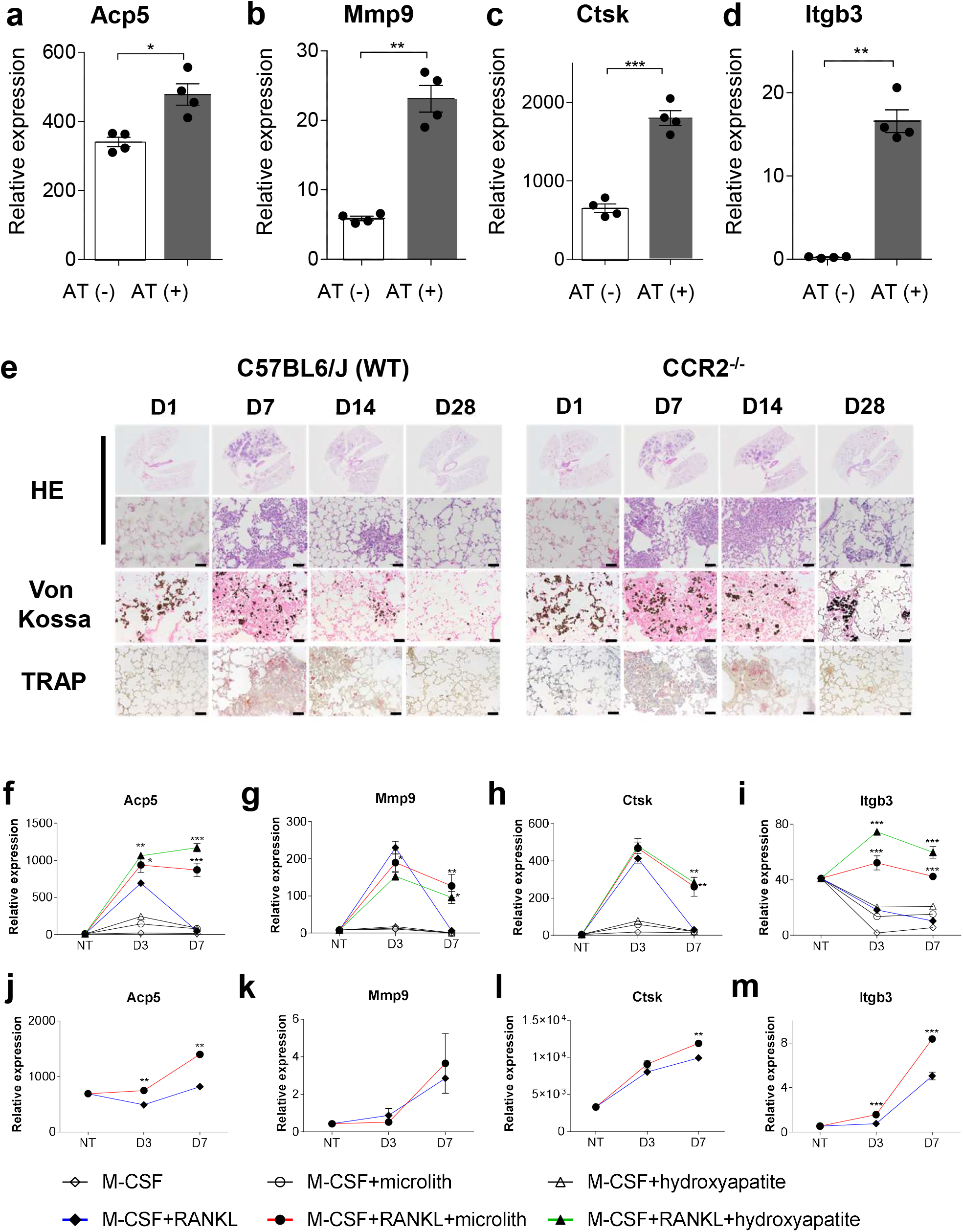
Adoptive transfer of hydroxyapatite microliths induces osteoclast expression, functional activity and stone clearance in Npt2b^+/+^ mice. **a-e**, Microliths isolated from the lungs of Npt2b^-/-^ mice were instilled into the lungs of C57BL/6J mice and CCR2^-/-^ mice. **a-d**, BAL was performed on C57BL/6J mice at day 7 post microlith instillation, and expression of osteoclast-related genes in BAL cells was determined by RT-PCR. n = 4 mice per group. Data are expressed as means ± SD. *p<0.05, **p<0.01 and ***p<0.001. **e**, CCR2^+/+^ and CCR2^-/-^ mice were sacrificed at the indicated time points after intra-tracheal microlith instillation, and the lungs were formalin fixed and stained with H&E, von Kossa, and TRAP reagent as indicated. Representative images are shown. Bold scale bar, 50 μm. **f-m**, BMDM (**f-i**) and alveolar macrophage (**j-m**) of C57BL/6J mice were cultured in presence of M-CSF ± RANKL with microliths (either isolated from the lungs of Npt2b^-/-^ mice or hydroxyapatite microliths) for 3 days or 7 days. Relative expression of osteoclast related genes in the cells was determined by real time PCR. n = 4. Data are expressed as means ± SD.*p<0.05, **p<0.01 and ***p<0.001 between M-CSF + RANKL and M-CSF + RANKL + microlith or M-CSF + RANKL + hydroxyapatite.

### Osteoclastogenesis in primary bone marrow derived monocytes cultured with RANKL and microliths

We next assessed the effect of microliths on RANKL-induced osteoclast differentiation of bone marrow derived monocytes (BMDM) and AM in culture. The combination of macrophage colony-stimulating factor (M-CSF) and RANKL are required for osteoclast survival in culture and induced osteoclast related gene expression in BMDM by day 3, but sustained expression at day 7 required exposure to either microliths or synthetic hydroxyapatite crystals in addition to these cytokines (Fig. 4f-i). Osteoclast related gene expression did not occur in the absence of RANKL. The combination of M-CSF and RANKL ± microliths also increased osteoclast related gene expression in AM, albeit to a more modest degree (Fig. 4j-m), and sustained osteoclast gene activation did not require exposure to microliths or hydroxyapatite. These results indicate that calcium phosphate crystals potentiate RANKL-induced osteoclast differentiation in monocytes and macrophages.

### Low phosphate diet reverses microlith accumulation in PAM model mice

To determine the effect of dietary phosphate intake on microlith accumulation, we fed Npt2b^+/+^ and Npt2b^-/-^ mice with low phosphate diets (LPD; 0.1% phosphate), regular diets (RD; 0.7% phosphate) and high phosphate diets (HPD; 2% phosphate) for a total of 8 weeks. The pre- and post-dietary treatment chest X-rays and micro CTs revealed that LPD reduced and HPD increased the profusion of calcific densities in all lung fields (Fig. 5a-c). Quantitative CT also demonstrated that LPD significantly decreased microlith burden (Fig. 5d, e). Next we treated Npt2b^+/+^ and Npt2b^-/-^ mice with LPD, RD or HPD for 1 week to analyze the effect of dietary phosphate intake on phosphate and calcium homeostasis. Serum parathyroid hormone (PTH) levels were higher on a regular diet at baseline in Npt2b^-/-^ mice than in Npt2b^+/+^ mice (Fig. 5f), and tended to be higher in PAM patients than healthy controls (Fig. 5g), but there were no other significant differences from cognate controls in NPT2B deficient mice or humans in serum fibroblast growth factor 23 (FGF-23) (Fig. 5h, i), 1,25-dihydroxyvitamin D (VitD3) (Fig. 5j, k), phosphate (Fig. 5l, m) or calcium (Fig. 5n, o). LPD decreased serum phosphate, PTH and FGF-23 and increased VitD3 and calcium (Fig. 5f, 5h, 5j, 5l and 5n). BAL phosphate and calcium concentrations were modulated by dietary phosphate intake (Fig. 5p and 5q) in Npt2b^-/-^ but not in Npt2b^+/+^ mice. Slc20a1 (Pit1) and Slc20a2 (Pit2) phosphate transporter gene expression was increased in isolated AECII from LPD treated mice compared to RD or HPD treated animals in mice of both genotypes. (Fig. 5r, s). Xenotropic and polytropic retrovirus receptor 1 (Xpr1) phosphate transporter, transient receptor potential cation channel subfamily V member 6 (Trpv6) and S100 calcium binding protein G (S100g) calcium transporter gene expression were also increased by LPD compared to HPD (Supplementary Fig. 1a-c). These results indicate that phosphate and calcium levels in BALF in Npt2b^-/-^ mice are sensitive to dietary phosphate content, and that the decrease of phosphate and calcium levels in alveolar space upon treatment with LPD may in part be the result of induction of alternative phosphate and calcium transporters in the lung.

**Fig. 5.**
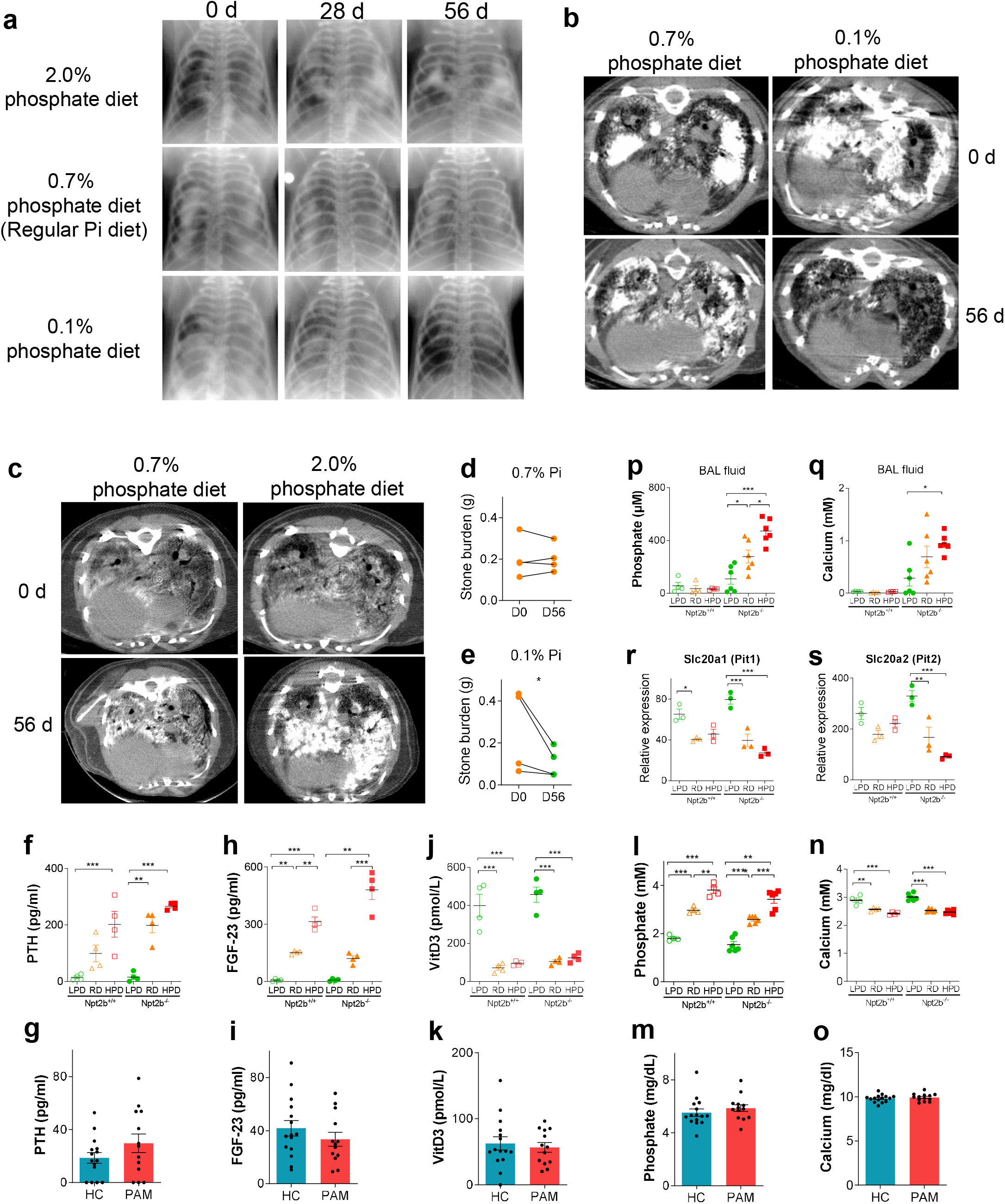
Dietary phosphate modulates phosphate homeostasis and stone accumulation in Npt2b-/- mice. **a-e**, 20 to 28 week old Npt2b^-/-^ mice (**a, b**) or 5 to 6 week old Npt2b^-/-^ mice (**c**) were fed with low (0.1%) (LPD), regular (0.7%) (RD) or high (2%) phosphate diet (HPD) for 8 weeks as indicated. **a**, Serial radiographs of individual Npt2b^-/-^ mice after 0, 28 and 56 days on the indicated phosphate diet. **b**, Serial micro CT images of individual Npt2b^-/-^ mice before and after 56d on RD (0.7% phosphate (Pi)) and LPD (0.1% Pi) diet. **c**, Micro CT images of Npt2b^-/-^ mice before and after 56 days on RD (0.7% Pi) and HPD (2.0% Pi) diet. **d, e**, Stone burden in the lung at day 0 and day 56 calculated from CT images of mice treated with RD (0.7% Pi) and LPD (0.1% Pi). **f, h, j, l, n**, and p-s, 5 to 6 week old Npt2b^+/+^ and Npt2b^-/-^ mice were fed with 0.7% Pi diet for 2 weeks as pretreatment, then regular, low or high phosphate diets (0.7% Pi (RD), 0.02% Pi (LPD) or 2% Pi (HPD), respectively) for 1 week. Serum and BAL fluid were collected after treatment; PTH (**f**), FGF-23 (**h**), VitD3 (**j**), phosphate (**l**) and calcium (**n**) levels in serum were determined. Phosphate (**p**) and calcium (**q**) levels in BAL fluid were determined. Relative expression of phosphate transporters, Slc20a1 (**r**) and Slc20a2 (**s**) in the isolated AECII cells from Npt2b^+/+^ and Npt2b^-/-^ mice fed with each indicated diet were determined by RT-PCR. N = 4 to 6 mice per group. Serum was collected from PAM patients (PAM) (n=15) and healthy controls (HC) (n=13), and PTH (**g**), FGF-23 (**i**), VitD3 (**k**), phosphate (**m**) and calcium (**o**) levels in serum were quantified by ELISA. Data are expressed as means ± SD. *p<0.05, **p<0.01 and ***p<0.001.

### RANKL-dependent osteoclast activation is required for optimal microlith clearance

We found that 2 weeks of LPD treatment of Npt2b^-/-^ mice decreased microlith burden and was associated with a decrease in TRAP positive MNGC accumulation in the lungs compared to RD treatment (Fig. 6a, b). Although HPD increased microlith burden, fewer TRAP positive cells were found in the lung. Osteoclast related gene expression profiles in whole lung homogenates revealed higher levels for the RANKL receptor, receptor activator of nuclear factor-κB (RANK), and lower levels for Acp5 and Mmp9 in the lungs of HPD treated mice compared to RD treated animals (Fig. 6c-e). Osteoprotegerin (OPG), which acts as a sink for RANKL, was significantly elevated in BALF after HPD treatment compared to LPD or RD treatment (Fig. 6f) of Npt2b^-/-^ mice, providing a plausible explanation for the decrease in RANKL dependent osteoclast gene expression and MNGC formation that were found in the lungs of these animals (Fig. 6a, b). To confirm the essential role of RANKL in osteoclast differentiation in PAM lung, we inhibited RANKL dependent osteoclastogenesis in Npt2b^-/-^ mice using a neutralizing anti-RANKL antibody. The fluorescent labeled bisphosphonate, Osteosense, was used to label microliths within the alveoli of living animals and assess the effect of the RANKL neutralization on microlith clearance. Osteosense was injected intratracheally into Npt2b^-/-^ mouse lungs and the decay in the fluorescence signal emanating from the microliths was tracked serially using IVIS for 12 days. Anti-RANKL neutralizing antibody treatment (3 times per week, i.p.) reduced the abundance of TRAP positive MNGCs (Fig. 6g, h) in histological sections and inhibited in vivo microlith clearance compared to control antibody treatment (Fig. 6i, j and Supplementary Fig. 2) or no treatment (not shown). Rather than MNGCs attached to stones, weakly TRAP positive mononuclear cells that were not aggregated or attached to stones were found in the lung sections of anti-RANKL treated Npt2b^-/-^ mice (Fig. 6g). These results indicate that RANKL-dependent osteoclast formation and actions are required for microlith clearance.

**Fig. 6.**
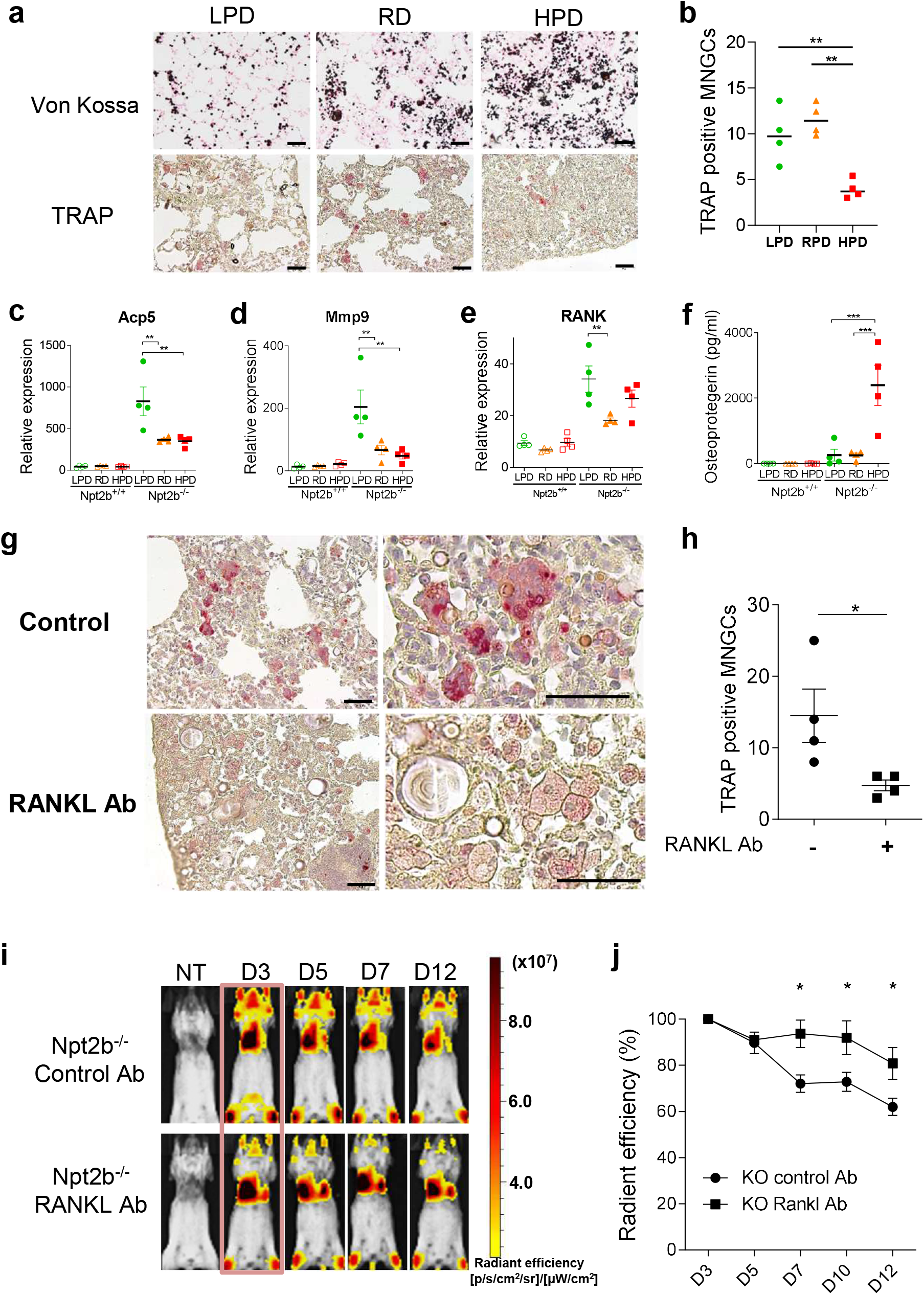
RANKL dependent osteoclast activity is required for microlith clearance. **a, b**, 5 to 6 week old Npt2b^+/+^ and Npt2b^-/-^ mice were fed with regular 0.7% Pi diet (RD) for 2 weeks as pretreatment, then fed RD, 0.02% Pi (LPD) or 2% Pi (HPD). Von Kossa and TRAP staining of the lung after 2 weeks of RD, LPD and HPD treatment (**a**) and TRAP positive MNGCs were counted in 10 random high-power fields (×100) and the average number was used for analysis (**b**). c-e, Relative expression of Acp5 (**c**), Mmp9 (**d**) and RANK (**e**) in the whole lung and osteoprotegerin levels (**f**) in BALF were determined by RT-PCR and ELISA respectively after 1 week of RD, LPD and HPD treatment. N = 4 mice per group. g, h Npt2b^-/-^ mice were treated with the RANKL neutralizing antibody or a rat IgG2a isotype control IgG intraperitoneally for 28d, then TRAP staining was performed on the lung sections (**g**) and TRAP positive MNGCs were counted in 10 random high-power fields (×100) and the average number was used for analysis (**h**). N = 4 per group. **i, j**, Npt2b^+/+^ and Npt2b^-/-^ mice were treated with the RANKL neutralizing antibody or a rat IgG2a isotype control IgG intraperitoneally for 7d, and then injected intratracheally with the OsteoSense 750EX probe to label microliths. Fluorescent reflectance imaging were acquired on the IVIS at the indicated times (before probe injection, 3, 5, 7 and 12 d after probe injection). N = 6 per group. Data are expressed as means ± SD. *p<0.05, **p<0.01 and ***p<0.001.

## Discussion

This study reveals osteoclast-like differentiation and activity in mononuclear cells of the PAM lung, and a role for these cells in limiting the accumulation of microliths in PAM. Rapid progression of hyperdense pulmonary infiltrates in a PAM patient placed on alendronate, a bisphosphonate known to inhibit osteoclast function, suggested a potential role for osteoclasts in modulating the stone burden in PAM^4, 13^. The observation that the microliths from humans and mice contained osteoclast bone degrading proteins CTSK, CALCR and TRAP, was also consistent with this notion. Gene expression analyses by single-cell RNA-seq in a human PAM lung and RT-PCR in Npt2b^-/-^ mouse lung revealed robust osteoclast signatures in pulmonary myeloid cells. Histologically, abundant TRAP and CTSK positive MNGCs were found attached to stones in mouse and human lung, and BAL cells isolated from mice that had been challenged with hydroxyapatite degraded bone when plated on slices of bovine femur. A low phosphate diet prevented and reversed microlith accumulation by mechanisms that included down modulation of alveolar OPG expression, and upregulation of osteoclast activity and calcium and phosphate transporter expression. Inhibition of osteoclastogenesis with anti-RANKL antibody attenuated osteoclast gene expression, diminished TRAP and CTSK positive MNGC formation, and slowed the clearance of stones from the lungs of Npt2b^-/-^ mice. Collectively, these data support a role for pulmonary osteoclast-like cells in pulmonary homeostasis for a rare disease, and raise the question of whether these newly described pulmonary cells may play a broader pathogenic or physiologic role in the lung.

Phosphate liberated by the catabolism of surfactant by AM is secreted into the epithelial lining fluid (ELF). The loss of functional Npt2b, due to inactivating mutations of the sodium phosphate cotransporter, Slc34a2, results in a defect in epithelial phosphate transport and an increase in ELF phosphate. Calcium also increases in the ELF of the PAM lung by an unknown mechanism, perhaps simply by passive action to maintain electroneutrality, and complexes with phosphate leading to the accumulation of microliths in the lumen of the alveolus. Analysis of microliths from PAM patients and Npt2b^-/-^ mice reveals a bone-like, porous surface, a simple elemental composition limited to calcium and phosphate at a 5:3 ratio consistent with hydroxyapatite, and a matrix containing abundant proteins and lipid species. The presence of surfactant proteins and lipids in the matrix of the microlith confirmed suspicions that microliths incorporate alveolar macromolecular components as had been suggested by their lamellar appearance on histological sections^11^.

Osteoclasts are specialized cells derived from circulating hematopoietic cells of the monocyte/macrophage lineage^14^. Osteoclast differentiation typically occurs on the bone surface and progresses in stepwise manner from mononuclear myeloid progenitor to polykaryon to activated osteoclast^15^. Under the influence of RANKL and M-CSF, osteoclastogenic genetic programs are executed by transcription factors, PU.1, MITF, TRAF6, NFkb and NFATc1 that drive the expression of proteins required for cell fusion, attachment to bone, and secretion of acid and proteases to degrade mineralized osseous tissues^16^. Membrane bound RANKL expressed on the plasma membrane of osteocytes is thought to be the major source of this primary osteoclastogenic cytokine, requiring cell to cell contact for engagement of the RANK receptor on the surface of osteoclast precursors, but soluble RANKL which is liberated by the action of proteases also plays an important role^17, 18^. OPG produced by osteoblasts, epithelial cells, endothelial cells, dendritic cells and B cells serves as a sink for RANKL, by binding and neutralizing the cytokine to modulate osteoclastogenesis^19^. DCSTAMP, OCSTAMP and DNAX activation protein of 12 kDa (DAP12) are some of the effectors known to be required for the fusion of mononuclear progenitors, to yield multinuclear giant cells with up to 15-20 nuclei^20–22^. Cytoskeletal rearrangements form actin rings that create a zone of occlusion when ITGB3 on the osteoclast plasma membrane^23^ binds to the asp-gly-xxx motif of OPN and other sialoproteins on bone. Activated osteoclasts secrete hydrochloric acid into the potential space that is formed, called Howship’s lacunae, through active transport of H^+^ and Cl^-^ by acid pumps including ATP6V0D2 and chloride voltage-gated channel 7 (CLCN7), respectively, to degrade the mineral components of bone^24, 25^. The exposed bone matrix is then degraded by CTSK and MMP9 and other secreted proteases^26, 27^. Partial dephosphorylation of OPN by TRAP leads to dissociation of the ITGB3/OPN bond and frees the osteoclast to migrate to new areas of bone^28^. A redox active iron in TRAP catalyzes the generation of reactive oxygen species, including hydrogen peroxide, hydroxyl radicals and singlet oxygen, which facilitate bone resorption and degradation^29^.

Many of these features of osteoclastogenesis outlined above were apparent in the PAM lung and Npt2b^-/-^ mouse lung. CTSK, TRAP and CALCR expressing cells were found attached to the microliths (Fig. 3g-r). The gene expression programs in the human PAM lung that were identified by RNA-seq demonstrated marked upregulation of osteoclast genes in monocyte/macrophages, including key transcription factors NFATc1, NFkB (IKBKB) and MITF, fusion proteins OCSTAMP and DCSTAMP, bone degrading proteins ATP6V0D2 and CLCN7, TRAP, CTSK and MMP9, and cell surface receptors colony stimulating factor 1 receptor (CSF1R) and CALCR. The upregulation of many of these genes was documented by RT-PCR in BAL cells from Npt2b^-/-^ mice and in Npt2b^+/+^ mice after adoptive transfer of microliths, including those characteristic of pre-fusion osteoclasts such as Acp5 and Mmp9, and also mature and activated osteoclast markers such as Ctsk and Calcr. Signature osteoclast enzymatic activities that were documented to be upregulated in the Npt2b^-/-^ lung included CTSK (Fig. 3s, t) and TRAP (Fig. 3g-j), and BAL cells from hydroxyapatite challenged Npt2b^+/+^ mice acquired the capacity to degrade bone (Fig. 3u). The delayed clearance of adoptively transferred microliths in CCR2^-/-^ mice compared to control mice suggests that recruited monocytes play an important role in microlith clearance. However, osteoclast differentiation and microlith clearance were not inhibited completely in CCR2^-/-^ mouse lung, indicating tissue resident AM contribute to osteoclast-like MNGCs differentiation and stone degradation^30^. Transient osteoclast gene expression was induced in AM and BMDM that were incubated with M-CSF and RANKL, but durable expression in BMDM required co-incubation with microliths. The demonstration that anti-RANKL antibody slowed the clearance of microliths from the lung provides evidence that osteoclast like MNGCs play a role in limiting microlith accumulation in the PAM lung.

Prior reports of true osteoclast-like cell differentiation in the lung have been limited to descriptions of MNGCs within lung tumors which most likely form as a result of DNA mutations and aberrant gene expression^31^. Inflammatory macrophages (IMs), MNGCs and foreign body giant cells (FBGCs) that express osteoclast markers are present in multiple benign lung diseases, however, including pneumoconiosis (e.g. silicosis and asbestosis) and granulomatous lung diseases (e.g. granulomatosis with polyangiitis, sarcoidosis and tuberculosis)^32–34^. Although there is no consensus on the cellular features and functions that reliably distinguish osteoclasts from IM, MNGC or FBGC, formation of a ruffled border, polarized expression of CTSK and ATP6V0D2 at the osteoclast/bone interface, the expression of the CALCR in myeloid cells, and ability to degrade both the mineral and matrix components of bone are perhaps the most unique to osteoclasts, and the latter two were documented in our models. While we found that CALCR protein expression is upregulated in the lungs of Npt2b^-/-^ mice and PAM patients (Fig. 3o-r), we also note that CALCR is expressed in pulmonary airway epithelium (Fig. 3o, q), as well as other non-myeloid compartments in kidney, uterine corpus and cardiac myocytes ^35^.

Serum phosphate levels in Npt2b^+/+^ mice and Npt2b^-/-^ mice increased with increased dietary phosphate intake, as did BAL phosphate levels in Npt2b^-/-^ mice, but BAL phosphate levels remained constant in Npt2b^+/+^ mice despite 20-fold differences in the phosphate content of their diet and 2-fold differences in serum phosphate levels. These data reveal a previously unrecognized role for Npt2b in maintaining alveolar phosphate homeostasis.

We have previously reported that microlith accumulation can be limited or reversed by feeding mice a low phosphate diet^11^, and suggested LPD mediated upregulation of Slc20a1 (Pit1) and Slc20a2 (Pit2) gene expression on AECII may have compensated for loss of Npt2b-mediated phosphate transport activity. The finding that LPD decreased phosphate and calcium levels in BALF is consistent with augmentation of calcium and phosphate export from the ELF. The Pit1 and Pit2 transporters are known to be upregulated by VitD3^36, 37^, which was elevated in the serum of LPD treated animals compared to RD treatment. TRPV6 and S100G mediated calcium transport are also known to be VitD3 dependent and gene expression of these transporters were upregulated in LPD compared to HPD. We submit that upregulation of alternative ion transporters may play a role in reducing calcium and phosphate levels in the ELF of LPD treated Npt2b^-/-^ mice compared to RD mice, although definitive proof of the role of individual transporters will require the availability of specific inhibitors of phosphate transport or knockout animals.

Dietary phosphate intake is also known to regulate bone osteoclast activity^38, 39^. When phosphate intake and serum levels drop, bone osteoclasts are activated to liberate phosphate and protect the organism from hypophosphatemia. In the Npt2b^-/-^ model, in addition to modestly lowering serum and BAL phosphate concentrations, LPD increased expression of osteoclast signature genes Acp5, Mmp9 and RANK in BAL cells (Fig. 6c-e). Surprisingly, HPD treatment resulted in a marked increase in OPG in the alveolar lining fluid (Fig. 6f). OPG functions as a soluble decoy receptor that inhibits osteoclast differentiation by binding and neutralizing RANKL^19^. Elevated phosphate concentrations are known to inhibit osteoclast differentiation both by upregulating OPG levels and by direct action on osteoclast precursor cells^40^. In HPD treated Npt2b^-/-^ mice with elevated OPG in BAL, there were fewer MNGCs and more TRAP-positive mononuclear cells in the lungs compared to RD and LPD treated Npt2b^-/-^ mice, which is similar to the findings in Npt2b^-/-^ mice treated with neutralizing RANKL antibody. Collectively, these results indicate that the microlith burden in Npt2b^-/-^ mice is modulated by dietary phosphate intake, in part through differential effects of OPG and RANKL on osteoclast differentiation and activity.

Analysis of BAL fluid and serum from the Npt2b^-/-^ mouse previously identified monocyte chemoattractant protein-1 (MCP-1) and surfactant protein D (SP-D) as potential PAM biomarkers, which were then shown to be elevated in the serum of patients with PAM. In adoptive transfer experiments, MCP-1 peaked at d7 post-microlith challenge and returned to baseline at d28, coincident with clearance of both microliths and resolution of inflammation^11^. The abundance of OPN in the proteome of the human and murine microliths led us to the finding that protein was also elevated in BAL fluid and serum of the Npt2b^-/-^ mouse, and in the serum of PAM patients, positioning OPN as another potential PAM biomarker. OPN is a multifunctional protein which is highly expressed in bone, macrophages, and epithelial cells of kidney, urinary duct, vessel and lung that is secreted in response to cellular stress, including exposure to high phosphate environments^41^. In PAM lung, OPN may protect alveolar epithelial cells against microlith induced inflammation.

It is tempting to speculate that the fibrosis that occurs in the PAM lung may be a consequence of collateral tissue injury from osteoclast products such as hydrochloric acid, TRAP, CTSK, MMP9. We intend to examine this question using animals that are deficient in genes that encode these phosphatases, proteases and ion pumps, and well as those that regulate osteoclastogenesis (e.g. RANKL, NFATC1). This concept is worth exploring since, if proven correct, there are many FDA approved therapies that target osteoclast viability and function.

In summary, studying the mechanisms of microlith clearance in the rare lung disease PAM led us to identify an osteoclast like pulmonary myeloid cell that plays an important role in limiting the stone burden in the lung. Augmenting the numbers or functioning of these cells represents a promising strategy for PAM therapeutic trials. Future studies from the laboratory will focus on whether pulmonary osteoclast like cells play a role in the response to other particulate challenges and in development of pulmonary fibrosis.

## Methods

### Mice

C57BL/6J WT mice were obtained from Jackson Laboratories (Bar Harbor, ME). The epithelium-targeted Npt2b^-/-^ mouse model was developed by breeding mice homozygous for floxed Slc34a2 with mice expressing Cre recombinase under the influence of the sonic hedgehog (Shh) promoter, as previously reported^11^. CCR2^-/-^ mice, B6.129S4-Ccr2^tm1Ifc^/J were purchased from Jackson Laboratories (Bar Harbor, ME, USA). For terminal experiments, mice were euthanized by intraperitoneal injection of Euthasol (Virbac, Fort Worth, TX). All animals were maintained in a specific pathogen– free facility and were handled according to a University of Cincinnati Institutional Animal Care and Use Committee–approved protocol and National Institutes of Health guidelines.

### Preparation of crystals

Synthetic hydroxyapatite particles were purchased from Fluidinova (Moreira da Maia, Portugal). Mouse microliths were isolated from the lungs of Npt2b^-/-^ mice after euthanasia by intratracheal instillation of 3 ml of dispase (50 caseinolytic units/ml, Corning, NY), submersion of the organ in 1 ml of dispase for 45 min at room temperature, and transfer to a culture dish containing water. The parenchymal lung tissue was gently teased from the bronchi, then homogenized. Human PAM microliths were isolated from the lung explant of a PAM infant undergoing lung transplant, by gently teasing the lung parenchyma apart with forceps in a culture dish containing water. Microliths were collected by centrifugation, and washed 5 times with cell culture grade water. For some experiments, mouse microliths were isolated without dispase treatment.

### Scanning electron microscopy and energy-dispersive X-ray spectroscopy analyses

Microliths were collected without dispase treatment from Npt2b^-/-^ mice and a PAM patient explant as describe above. Microliths were incubated with distilled water or 0.02% SDS for 5 min, then washed 5 times with water to remove residual SDS and dried. The samples were placed on conductive carbon tape. Imaging was performed at 2kV on a scanning electron microscope (FESEM, FEI Scios DualBeam, Germany). The elemental analyses were obtained at 15 kV beam voltage using an EDAX Octane Elite Super detector.

### Protein extraction and protein identification

Microliths were incubated in buffer containing 50 mM TEAB, 5 mM Tris, and 75 mM EGTA with sonication and end over end shaking in the cold room. The supernatant was then buffer exchanged with 50 mM TEAB for 3 cycles using 3 kDa filters (Amicon). The final volume of the sample was brought to 100 µl with 50 mM TEAB. The samples were reduced with 20 µl of 50 mM tris-(2 carboxyethyl) phosphine (TCEP), alkylated with 10 µl of 200 mM methyl mehanethiosulfonate (MMTS), digested with trypsin and dried in a lyophyllizer. Samples were analyzed by nanoLC-MS/MS on a Sciex 5600 quadrupole-TOF system and identified using mus musculus or the homo sapien databases with the Protein Pilot (ver 5.0, rev 4769) program (Sciex) as described previously^42^.

### Lipid extraction and lipid identification

Total lipid extracts (TLEs) were collected using a modified Folch extraction^43, 44^. Briefly, chloroform, methanol and water were added to the microliths to a final ratio of 8:4:3 respectively. The samples were vortexed for 30 sec, chilled on ice for 5 minutes, vortexed again and centrifuged at 10,000 x g for 10min at 4℃. The lower organic lipid containing layer was removed, dried in vacuo and reconstituted in 50 µl of methanol. Due to the differing yields of TLE from Npt2b^+/+^ and Npt2b^-/-^ mice, the Npt2b^+/+^ BALF pellets were reconstituted in 100 µl of methanol and the Npt2b^-/-^ BALF in 1500 µl of methanol. Samples were analyzed using mass spectrometry, using LC-MS/MS parameters and identification algorithms outlined by Kyle et al.^45^. A Waters Aquity UPLC H class system interfaced with a Velos-ETD Orbitrap mass spectrometer for LC-ESI-MS/MS analyses. Reconstituted TLEs were injected onto a reversed phase Waters CSH column (3.0 mm x 150 mm x 1.7 µm particle size), and lipids were separated over a 34 min gradient (mobile phase A: ACN/H2O (40:60) containing 10 mM ammonium acetate; mobile phase B: ACN/IPA (10:90) containing 10 mM ammonium acetate) at a flow rate of 250 µl/min. Samples were analyzed in both positive and negative ionization modes using HCD (higher-energy collision dissociation) and CID (collision-induced dissociation). The LC-MS/MS raw data files were analyzed using LIQUID and all identifications were manually validated^45^ which included examining the fragmentation spectra for diagnostic ions and fragment ions corresponding to the acyl chains, the precursor mass isotopic profile and mass ppm error, the extracted ion chromatograph, and retention time. To facilitate quantification of the lipids, a reference database for BALF lipids identified from 4 of the raw data files (2 WT and 2 KO) using LIQUID was created and features from each analysis were aligned to the reference database based on their identification, m/z and retention time using MZmine 2^46^. A separate reference dataset was created from both of the microlith samples analyzed. Aligned features were manually verified and peak apex intensity values were exported for subsequent statistical analysis.

### Single-cell RNA-seq analysis

Cell Ranger R kit version 2.1.1 (10X Genomics) was used to process raw sequencing data and paired-end sequence alignment to the human genome (hg19). We used Sincera^47, 48^ and Seurat 2^49, 50^ for downstream analysis. For pro-filtering, we required that cells express more than 500 genes (transcript count > 0), and that less than 10% of transcript counts mapped to mitochondrial genes. Genes with transcripts detected in less than 2 cells were removed. Total of 14210 cells and 21,075 genes passed the pro-filtering and were used for further analysis. Transcript Counts in each cell were normalized by dividing by the total number of transcripts in each cell multiplied by 10,000 and log-transformed (normalized count +1). For clustering, principal-component analysis was performed for dimension reduction. Reduced dimensions were used for cell cluster identification using the Jaccard-Louvain clustering algorithm^51^. For cell type mapping, we first identified major cell types (i.e. -epithelial, endothelial, immune, and mesenchymal) in PAM and control lungs separately, and then integrated the data using “Seurat Alignment” to remove batch effect. Major cell types from PAM and control lungs were merged. We then extracted the immune cell cluster for sub-clustering. For this study, we choose to focus on macrophage subtypes. All the downstream analysis including signature markers comparison, functional enrichment and pathway analysis were done in macrophages of PAM and donor samples. Differential expression of genes across the PAM lung sample and control lung sample were tested using a nonparametric binomial test^51^. Cell specific signature genes were defined based on following criteria: (1) <0.1 false discovery rate (FDR) of the binomial test, (2) minimum two-fold effective size, (3) detection in at least 20% of cells in the group.

### Histology and immunohistochemistry

Mouse and human lung tissues were fixed with 10% buffered formalin phosphate, embedded in paraffin, and stained with hematoxylin and eosin (H&E). The von Kossa technique was used to stain microliths by incubation of histological sections with 3% silver nitrate, exposure to UV light for 30 min, and counterstaining with Nuclear Fast Red (Newcomer Supply; Middleton, WI) for 5 min. For TRAP staining, tissue sections were incubated with pre-warmed (37 °C) TRAP staining mix (University of Rochester Medical Center TRAP protocol) for 1h and counterstaining with Harris hematoxylin for 5 sec. Immunohistochemical analysis was performed on formalin-fixed, paraffin embedded material. Five micrometer sections were deparaffinized and rehydrated with dH_2_O. The sections were subjected to antigen retrieval with citrate buffer, blocked with normal serum, and probed with specific primary antibodies anti-OPN (ab8448, polyclonal, 1:7500) and anti-Cathepsin K (ab19027, polyclonal, 1:750) obtained from Abcam (Cambridge, UK), anti-CALCR LS (LS-A769, polyclonal, 9ug/ml) obtained from LS Bio (Seattle,WA) followed by horseradish peroxidase linked anti-rabbit secondary antibody (Cell Signaling Technology Inc., Beverly, MA). Slides were developed with DAB substrate, counter stained with hematoxylin, dehydrated, and mounted.

### Adoptive transfer of microliths

Microliths recovered from Npt2b^-/-^ mice lungs were washed with water 5 times then dried. Dried microliths were resuspended in normal saline at a concentration of 100 mg/ml, and 100 µL of the suspension was intratracheally instilled into each mouse, followed by histologic and radiographic analyses at time points indicated.

### BAL cell and fluid collection

Mice were sacrificed and subjected to BAL by intratracheal intubation followed by 5 cycles of instillation and aspiration of one ml of 0.9% saline. The lavage was pooled on ice and BAL cells were pelleted by centrifugation at 500 x g for 10 minutes at 4°C. Gene expression in BAL cells was assessed by RT-PCR, and the supernatant from the first lavage was used for ELISA and colorimetric assays.

### Cell culture

Primary mouse BMDM or AM were prepared from 8 to 10-week-old C57BL/6J mice. BMDM were isolated with a monocyte isolation kit (Miltenyi Biotec, Bergisch Gladbach, Germany) following the manufacturer’s instructions. AM were collected from BALF as described above. BMDM and AM were cultured in Dulbeccòs Modified Eagle’s Medium (DMEM) + 10% heat-inactivated fetal bovine serum (FBS) supplemented with 20 ng/ml recombinant murine M-CSF (PeproTech, Rocky Hill, NJ) and 20 ng/ml recombinant murine RANKL (PeproTech). Medium was changed every 2 to 3 days. At day 3 or day 7 of culture, total RNA was purified from cells for RT-PCR analysis.

### Serum and alveolar lining fluid collection and measurements

Phosphate and calcium levels in serum and BALF were measured using the Phosphate Colorimetric Assay Kit (BioVision, Milpitas, CA) and Calcium Colorimetric Assay Kit (BioVision), respectively. ELISAs were used to measure SP-D (Rat/Mouse SP-D kit, YAMASA, Tokyo, Japan), and MCP-1 (CCL2 /JE/MCP-1 ELISA kit, R&D Systems Inc., Minneapolis, MN) in mouse serum. Osteopontin in mouse BALF was measured using the Mouse/Rat Osteopontin Duoset ELISA kit, and human serum osteopontin was measured using the Human Osteopontin DuoSet ELISA kit (R&D Systems Inc.). Phosphate homeostasis hormones were measured in mouse and human serum using the FGF-23 ELISA Kit (Kainos Laboratories Inc., Tokyo, Japan), the mouse or human PTH 1-84 ELISA Kit (Immutopics, San Clemente, CA), the 25-Hydroxy Vitamin DS EIA kit (Immunodiagnostic Systems, Boldon, UK) and 1,25-Dihydroxy Vitamin D EIA kit (Immunodiagnostic Systems). Human sample collection and laboratory analysis were approved by the Institutional Review Board at The University of Cincinnati College of Medicine.

### Preparation of RNA and RT-PCR

RNA was isolated from murine BAL pellet or cultured cells using RNAzol RT (Molecular Research Center, Cincinnati, OH); RNA from tissues of Npt2b^-/-^ and Npt2b^+/+^ mice was isolated using the RNeasy Micro Kit (Qiagen, Hilden, Germany). cDNA was generated using the SuperScript III First-Strand Synthesis System (Life Technologies, Carlsbad, CA). After first-strand complementary DNA synthesis using SuperScript III Reverse Transcriptase (Invitrogen, Carlsbad, CA), quantitative RT-PCR was performed using a SYBR Green Master Mix (Applied Biosystems, Foster City, CA) and primer pairs for sodium phosphate cotransporters, as well as β-actin as an internal control. The nucleotide sequences of the primer pairs were 5′-GCCACAGTTATGTTTGTACGTG-3′ and 5′-ACAGATTGCATACTCTAAGATCTCC-3′ for Acp5, 5′-GTGGGAGGTATAGTGGGACA-3′ and 5′-GACATAGACGGCATCCAGTATC-3′ for Mmp9, 5′-ATCTCTCTGTACCCTCTGCAT-3′ and 5′-GACTCTGAAGATGCTTACCCA-3′ for Ctsk, 5′-ACAGTCATCCTCGTTCTTGTAG -3′ and 5′-GAACGCTCCATGAAGAAAACAC-3′ for Itgb3, 5′-GGTTTGCCTCATCTTGGTCA-3′ and 5′-TCTACTACAACGACAACTGCTG-3′ for Calcr, 5′-GGAAGATGGTAGGAGAGGGTA-3′, and 5′-AGGATGAGGACAGACAGGT-3′ for Csf1, 5′-AGTGCTGTCTTCTGATATTCTGT -3′ and 5′-CAGGAGAGGCATTATGAGCAT-3′ for Tnfrsf11a, 5′-GTGTCCCTTCTCTTCCAGTTC-3′ and 5′-TGTTGCCGCTTTTGTAGAG-3′ for Slc20a1, 5′-CCTGCTCTTCCACTTCCTG-3′ and 5′-TCTTGTGTAACTCCGCCTTG-3′ for Slc20a2 and, 5′-ACCTTCTACAATGAGCTGCG-3′ and 5′-CTGGATGGCTACGTACATGG-3′ for β-actin.

### Phosphate diets

Regular-phosphate content chow (RD), low-phosphate chow (LPD) and high-phosphate (HPD) chow containing, respectively, 0.7%/0.02%/2.0% phosphate, 16.3%/16% /16.3% protein, 66.3%/68.0%/63.3% carbohydrate, 5.0%/5.0%/5.0% fat, 1.2%/1.2%/1.2% calcium and vitamin D3 (2900 IU/Kg) were specially prepared by Harlan Laboratories (Madison, WI) and purchased from Harlan Sprague Dawley (Indianapolis, IN, USA). For some experiments, 0.1 % phosphate containing chow were used. 5- to 6-week-old Npt2b^+/+^ and Npt2b^-/-^ mice were fed with RD for 2 weeks as pretreatment, followed by RD, LPD or HPD for the indicated time intervals. For RT-PCR studies, mice were assigned to a diet for 1 week, lungs were explanted and homogenized for RT-PCR. For histological analysis, mice were treated with RD, LPD or HPD for 2 weeks, lungs were fixed for H&E and von Kossa staining. To determine the effect of dietary phosphate restriction and excess on microlith burden in mice with mature PAM lesions, 20- to 28-week-old Npt2b^+/+^ and Npt2b^-/-^ mice were fed with 0.1% phosphate LPD, 0.7% phosphate RD or 2.0% phosphate HPD for 8 weeks, follow by analyses including chest x-ray, chest CT and bone density measurement.

### MicroCT, X-ray imaging and bone density

After animals were anesthetized with isoflurane, two-dimensional (2D) chest x-ray images (In-Vivo Multispectral Imaging System FX, Bruker) were obtained, and microCT scans (Inveon, Siemens) with respiratory gating were performed. For microlith burden assessment, a commercial graphic processing software package (Amira, FEI Company, Hillsboro OR) was used to segment the lung from other soft tissues in CT images. Mean lung density and percentage of high-density lung volumes (> 1.5 g/cc and > 1.25 g/cc, calculated from the Hounsfield units in CT images) were calculated from the entire lung in MATLAB. Total mass of the lung was calculated by multiplying average density by the CT-derived lung volume. Stone burden was calculated as the product of lung volume and the increase in lung density over the average density of wild-type lungs. Bone density was calculated from three minute x-ray acquisition of mouse femur by the Bruker MS FX pro imaging system bone density software module.

The bone density software estimates bone mineralization by modeling cylindrical symmetry to the analytical X-ray image of a long-bone segment (femurs) of mice with the appropriate calibration, based upon CaPO_4_ immersed in water. The software performs a cylindrical fitting routine to extract and calculate the important parameters of the selected (ROI) bone segment. Profiles are generated in rows over the bone segment contained within the ROI. The bone density software performs a chi square weighted average of all the profiles that converge successfully to produce twelve relevant results including a CaPO_4_ coefficient (cm^2^/g) and bone density (g/cm^3^).

### Isolation of AECII

Mice were sacrificed by intraperitoneal injection of Euthasol, and lungs were perfused with 10 ml of sterile normal saline via the pulmonary artery. The airway was cannulated via tracheostomy with a 20-gauge metallic angiocatheter, and 3 ml of dispase (50 caseinolytic units/ml, Corning) was instilled, followed by 0.5 ml of 1% low-melt agarose (warmed to 45°C). Lungs were rapidly cooled on ice for 2 min, submerged in 1 ml of dispase for 45 min at room temperature, and transferred to a culture dish containing deoxyribonuclease I (100 U/ml) (Worthington Biochemicals, Malvern, PA). The parenchymal lung tissue was gently teased from the bronchi, and homogenized. Cell suspensions were filtered, collected by centrifugation, and panned over prewashed 100-mm tissue culture plates coated with CD45 and CD16/32 antibodies (BD Biosciences, San Jose, CA). After incubation for 60 min at 37°C in a 5% CO_2_ atmosphere to promote adherence of contaminating macrophages and fibroblasts, the AECII were gently decanted from the plate, collected by centrifugation, and counted. For the Npt2b^-/-^ animals, differential centrifugation was used to separate microliths from the cells. Cell viability determined with trypan blue staining was routinely >90%, and cell purity determined by SP-C staining ranged from 75 to 90%.

### Fluorescent reflectance imaging (FRI)

Two commercially available fluorescent imaging probes (PerkinElmer Inc., Hopkinton, MA) were used to evaluate osteoclast functions in the lung. The fluorescent bisphosphonate imaging agent, OsteoSense 750EX (Osteosense) was used to tag microliths in the lung. The Cathepsin K cleavage activated fluorescent probe; Cat K 680 FAST probe (Cat K) were used to detect cathepsin K activity in the lung. Npt2b^+/+^ and Npt2b^-/-^ mice were shaved around the chest prior to imaging, and 1 nmol Cat K or 1 nmol Osteosense was administered intratracheally. The fluorescence signal emanating from the chest was monitored using fluorescent IVIS (IVIS Spectrum, PerkinElmer) at 18 h post-injection for Cat K, and at days 1, 3, 5, 7, 10 and 12 post-injection for Osteosense. Quantification of fluorescence intensity was performed by evaluating the total radiant efficiency ([photons/sec]/[μW/cm^2^]) of the signal within a region of interest (ROI). The ROI was defined by an area with a radius of 0.94 cm^2^ that encompassed the lungs of the mice [Fig. 3s] and the total radiant efficiency of Osteosense was normalized to that present at day 3, to limit the mouse-to-mouse variation in delivery of the fluorescent probe.

### Bone resorption assay

Hydroxyapatite (Fluidinova) microspheres were resuspended in normal saline at a concentration of 10 mg/ml, and 100 µL of the suspension or 100 ul of normal saline were intratracheally instilled into C57BL/6J mice. BAL cells were collected d14 after adoptive transfer of saline or hydroxyapatite. Freshly collected BAL cells were plated on bovine bone slices (Immunodiagnostic Systems) in 96-well plates (5.0 x 10^5^ cells/well) and incubated with DMEM containing 10% FCS in the presence of 100 ng/ml recombinant murine RANKL (PeproTech) with 20 ng/ml recombinant murine M-CSF (PeproTech). After 24 hours, medium was changed to remove unattached cells, and replaced with fresh medium. After 48 hours incubation, culture medium was collected and the levels of CTX-I in culture medium were measured using the the CrossLaps^®^ for culture (CTX-I) ELISA (Immunodiagnostic Systems).

### Anti-RANKL antibody treatment

Microliths in the Npt2b^-/-^ mice lung were labeled by intratracheal administration of Osteosense as described above. 250 μg/mouse of anti-mouse RANKL mAb (clone IK22-5, Bio X Cell, West Lebanon, NH) or a Rat IgG2a isotype control IgG (clone 2A3, Bio X Cell) were administrated intraperitoneally into Npt2b^-/-^ mice three times per week for 20 days or 28 days. For the IVIS study, mice were pre-treated with anti-RANKL for 7 days before Osteosense injection. The treatment effect on microlith dissociation and osteoclastogenesis were monitored by quantifying the residual fluorescence signal using IVIS or histological analysis of the lung.

### Statistical analysis

Statistical analyses were performed using GraphPad Prism 5. Significance between two groups was determined using Welch’s t-test. In experiments in which more than two groups were compared, a one-way analysis of variance (ANOVA) was used followed by Bonferroni’s multiple comparisons test. Differences were considered significant at p < 0.05.

## Author contributions

Yasuaki Uehara designed and performed experiments, analyzed data, and wrote the manuscript. Nikolaos M. Nikolaidis designed experiments and analyzed data. Lori B. Pitstick performed histological and pulmonary physiological experiments. Huixing Wu and Yoshihiro Hasegawa and Yusuke Tanaka performed ELISA assays. Jane J. Yu and Erik Zhang performed IVIS assay. John G. Noel and Jason C. Gardner isolated mouse BMMs. Elizabeth J. Kopras wrote the manuscript. Wendy D. Haffey and Kenneth D. Greis performed mass spectrometry and analyzed proteomic data. Jinbang Guo and Jason C. Woods performed quantitative CT analysis, Kathryn A. Wikenheiser-Brokamp prepared tissues for single cell RNA-seq and assisted with pathological analyses. Shuyang Zhao and Yan Xu analyzed single cell RNA-seq data. Jennifer E. Kyle and Charles Ansong performed mass spectrometry and analyzed lipidomics data. Steven L. Teitelbaum assisted with data analysis. Yoshikazu Inoue and Göksel Altinişik provided PAM patients samples. F.X.M developed the concept, designed experiments, analyzed data, and wrote the manuscript. All co-authors contributed to writing the manuscript.

## Acknowledgements

Mass Spectrometry data were collected in the UC Proteomics laboratory on an instrument funded in part by a National Institutes of Health (NIH) shared instrumentation grant (S10 RR027015) to KDG. We thank the Heydlauff family for donation of their infant son’s lung for this research.

