## SupplementaryInformation for "Novel insights into pulmonary phosphate homeostasis and osteoclastogenesis emerge from the study of pulmonary alveolar microlithiasis"

**Supplementary Fig. 1**

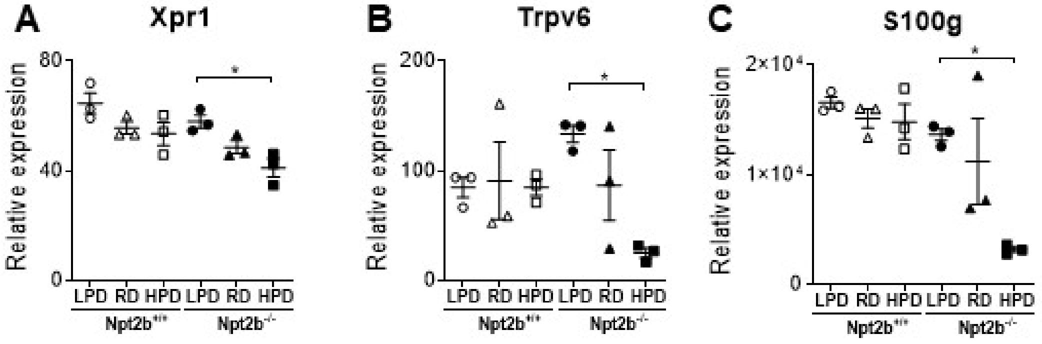


Supplementary Fig. 1 | Dietary phosphate modulates Xpr1 phosphate transporter and $VitD_3$ dependent calcium transporters in $Npt2b^{-/-}$ mice. 5 to 6 week old $Npt2b^{+/+}$ and $Npt2b^{-/-}$ mice were fed with 0.7% Pi diet for 2 weeks as pretreatment, then they were fed with regular, low or high phosphate diets (0.7% Pi (RD), 0.02% Pi (LPD) or 2% Pi (HPD), respectively) for 1 week. Relative expression of phosphate transporter Xpr1 (A), and calcium transporters, Trpv6 (B) and S100g (C) in the isolated AECII cells from $Npt2b^{+/+}$ and $Npt2b^{-/-}$ mice fed with each indicated diet were determined by RT-PCR. N = 4 mice per group. Data are expressed as means ± SD. $^*p<0.05$.

**Supplementary Fig. 2**

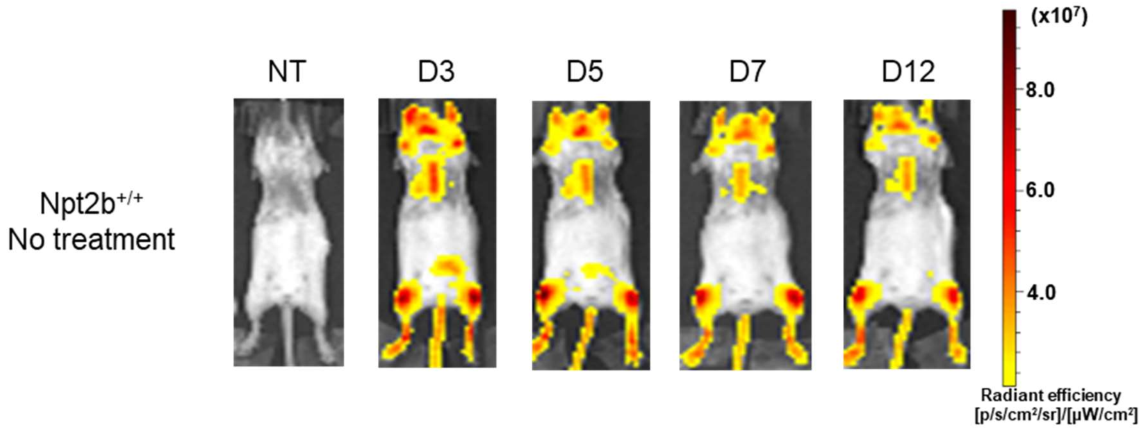


Supplementary Fig. 2 | Administration of OsteoSense 750EX in Npt2b$^{+/+}$ mouse lung. Npt2b$^{+/+}$ mice were injected intratracheally with OsteoSense 750EX probe. Fluorescent reflectance imaging were acquired on the IVIS at the indicated times (before probe injection, 3, 5, 7 and 12 d after probe injection). N = 6.

**Supplementary Table 1. The molar ratios and weight percentages of calcium and phosphorus in microliths.**

| Microliths | Weight percentage Ca | Weight percentage P | Ca/P Molar ratio |
|---|---|---|---|
| Hydroxyapatite | 29.12 ± 6.48 | 12.62 ± 2.84 | 2.31 ± 0.06 |
| Mouse microliths | 18.98 ± 1.19 | 9.47 ± 0.71 | 2.01 ± 0.03 |
| Human microliths | 18.50 ± 3.98 | 9.75 ± 1.40 | 1.88 ± 0.18 |

**Supplementary Table 2. Identified proteins from human microlith.**

Please refer to the EXCEL file attached separately.

**Supplementary Table 3. Identified proteins from mouse microlith.**

Please refer to the EXCEL file attached separately.

**Supplementary Table 4. Lipid species detected in the BALF and microlith isolated from human and Npt2b$^{-/-}$ mice.**

| | | Number of lipids identified | | |
|---|---|---|---|---|
| Subclass | | Mouse BALF | Mouse microlith | Human microlith |
| Carnitine | carnitine | 2 | 0 | 0 |
| CE | cholesteryl ester | 4 | 5 | 1 |
| Cer | ceramide | 12 | 6 | 16 |
| HexCer | hexosylceramide | 2 | 1 | 1 |
| LacCer | lactosylceramide | 1 | 1 | 1 |
| SM | sphingomyelin | 10 | 8 | 14 |
| LPC | lysophosphatidylcholine | 0 | 12 | 3 |
| PC | phosphatidylcholine | 88 | 80 | 45 |
| PCox | oxidized PC | 1 | 1 | 0 |
| PCO | 1-O-alkyl-2-acyl-PC | 5 | 6 | 5 |
| PCP | PC plasmalogen | 1 | 5 | 7 |
| LPE | lysophosphatidylethanolamine | 0 | 3 | 4 |
| PE | phosphatidylethanolamine | 28 | 19 | 16 |
| PEO | 1-O-alkyl-2-acyl-PE | 3 | 3 | 3 |
| PEP | PE plasmalogen | 14 | 15 | 20 |
| LPG | lysophosphatidylglycerol | 4 | 0 | 0 |
| PG | phosphatidylglycerol | 28 | 55 | 21 |
| PI | phosphatidylinositol | 1 | 13 | 7 |
| PS | phosphatidylserine | 0 | 1 | 0 |
| MG | Monoacylglycerol | 3 | 0 | 0 |
| DG | Diacylglycerols | 13 | 11 | 9 |
| TG | triacylglycerol | 20 | 15 | 23 |

### SUPPLEMENTAL METHODS

#### Isolation of AECII.

5 to 6 week old $Npt2b^{+/+}$ and $Npt2b^{-/-}$ mice were fed with 0.7% Pi diet for 2 weeks as pretreatment, then they were fed with regular, low or high phosphate diets (0.7% Pi (RD), 0.02% Pi (LPD) or 2% Pi (HPD), respectively) for 1 week. After treatment, mice were sacrificed by intraperitoneal injection of Euthasol, and lungs were perfused with 10 ml of sterile normal saline via the pulmonary artery. The airway was cannulated via tracheostomy with a 20-gauge metallic angiocatheter, and 3 ml of dispase (50 caseinolytic units/ml, Corning) was instilled, followed by 0.5 ml of 1% low-melt agarose (warmed to 45°C). Lungs were rapidly cooled on ice for 2 min, submerged in 1 ml of dispase for 45 min at room temperature, and transferred to a culture dish containing deoxyribonuclease I (100 U/ml) (Worthington Biochemicals, Malvern, PA). The parenchymal lung tissue was gently teased from the bronchi, and homogenized. Cell suspensions were filtered, collected by centrifugation, and panned over prewashed 100-mm tissue culture plates coated with CD45 and CD16/32 antibodies (BD Biosciences, San Jose, CA). After incubation for 60 min at 37°C in a 5% $CO_2$ atmosphere to promote adherence of contaminating macrophages and fibroblasts, the AECII were gently decanted from the plate, collected by centrifugation, and counted. For the $Npt2b^{-/-}$ animals, differential centrifugation was used to separate microliths from the cells. Cell viability determined with trypan blue staining was routinely >90%, and cell purity determined by SP-C staining ranged from 75 to 90%.

#### Preparation of RNA and RT-PCR.

RNA was isolated from murine AECII using RNAzol RT (Molecular Research Center, Cincinnati, OH). cDNA was generated using the SuperScript III First-Strand

Synthesis System (Life Technologies, Carlsbad, CA). After first-strand complementary DNA synthesis using SuperScript III Reverse Transcriptase (Invitrogen, Carlsbad, CA), quantitative RT-PCR was performed using a SYBR Green Master Mix (Applied Biosystems, Foster City, CA) and primer pairs for sodium phosphate cotransporters, as well as β-actin as an internal control. The nucleotide sequences of the primer pairs were 5′-CTGTCACCATCTTCAAGTTCATTG-3′ and 5′-GGCGAGTGATTCATGTAGAGG-3′ for Xpr1, 5'-CATGCTTAACCTCCTCATTGC-3' and 5'-TTCCGCTCTAACATCACAGTG-3' for Trpv6, 5'-TCAGAGTTCCCCAGCCT-3' and 5'-TCCTGACTTGTTCATTGTGAGAG-3' for S100g and, 5′-ACCTTCTACAATGAGCTGCG-3′ and 5′-CTGGATGGCTACGTACATGG-3′ for β-actin.

**Fluorescent reflectance imaging (FRI).**

The fluorescent bisphosphonate imaging agent, OsteoSense 750EX (Osteosense) was used to tag microliths in the lung. Npt2b$^{+/+}$ mice were shaved around the chest prior to imaging, and 1 nmol Osteosense was administered intratracheally. The fluorescence signal emanating from the chest was monitored using fluorescent live imaging system (IVIS) (IVIS Spectrum, PerkinElmer) at days 1, 3, 5, 7, 10 and 12 post-injection for Osteosense. Quantification of fluorescence intensity was performed by evaluating the total radiant efficiency ([photons/sec]/[μW/cm$^2$]) of the signal within a region of interest (ROI). The ROI was defined by an area with a radius of 0.94 cm$^2$ that encompassed the lungs of the mice [Fig. 3M].
