## Supplemental Table2 for "Novel insights into pulmonary phosphate homeostasis and osteoclastogenesis emerge from the study of pulmonary alveolar microlithiasis"

| Accession | Name | Species | Peptides(95%) |
| --- | --- | --- | --- |
| sp\|P02768\|ALBU_HUMAN | Serum albumin OS=Homo sapiens GN=ALB PE=1 SV=2 | HUMAN | 266 |
| sp\|P02765\|FETUA_HUMAN | Alpha-2-HS-glycoprotein OS=Homo sapiens GN=AHSG PE=1 SV=1 | HUMAN | 225 |
| sp\|P01024\|CO3_HUMAN | Complement C3 OS=Homo sapiens GN=C3 PE=1 SV=2 | HUMAN | 94 |
| sp\|P00734\|THRB_HUMAN | Prothrombin OS=Homo sapiens GN=F2 PE=1 SV=2 | HUMAN | 92 |
| sp\|P04004\|VTNC_HUMAN | Vitronectin OS=Homo sapiens GN=VTN PE=1 SV=1 | HUMAN | 68 |
| sp\|P10451\|OSTP_HUMAN | Osteopontin OS=Homo sapiens GN=SPP1 PE=1 SV=1 | HUMAN | 67 |
| sp\|P07900\|HS90A_HUMAN | Heat shock protein HSP 90-alpha OS=Homo sapiens GN=HSP90AA1 PE=1 SV=5 | HUMAN | 55 |
| sp\|P08238\|HS90B_HUMAN | Heat shock protein HSP 90-beta OS=Homo sapiens GN=HSP90AB1 PE=1 SV=4 | HUMAN | 45 |
| sp\|P01042\|KNG1_HUMAN | Kininogen-1 OS=Homo sapiens GN=KNG1 PE=1 SV=2 | HUMAN | 44 |
| sp\|P02647\|APOA1_HUMAN | Apolipoprotein A-I OS=Homo sapiens GN=APOA1 PE=1 SV=1 | HUMAN | 43 |
| sp\|P11021\|GRP78_HUMAN | 78 kDa glucose-regulated protein OS=Homo sapiens GN=HSPA5 PE=1 SV=2 | HUMAN | 38 |
| sp\|Q8IWL1\|SFPA2_HUMAN | Pulmonary surfactant-associated protein A2 OS=Homo sapiens GN=SFTPA2 PE=1 SV=1 | HUMAN | 37 |
| sp\|Q8IWL2\|SFTA1_HUMAN | Pulmonary surfactant-associated protein A1 OS=Homo sapiens GN=SFTPA1 PE=1 SV=2 | HUMAN | 36 |
| sp\|Q06481\|APLP2_HUMAN | Amyloid-like protein 2 OS=Homo sapiens GN=APLP2 PE=1 SV=2 | HUMAN | 35 |
| sp\|P26038\|MOES_HUMAN | Moesin OS=Homo sapiens GN=MSN PE=1 SV=3 | HUMAN | 31 |
| sp\|P01008\|ANT3_HUMAN | Antithrombin-III OS=Homo sapiens GN=SERPINC1 PE=1 SV=1 | HUMAN | 31 |
| sp\|P02545\|LMNA_HUMAN | Prelamin-A/C OS=Homo sapiens GN=LMNA PE=1 SV=1 | HUMAN | 30 |
| sp\|P02649\|APOE_HUMAN | Apolipoprotein E OS=Homo sapiens GN=APOE PE=1 SV=1 | HUMAN | 29 |
| sp\|P00747\|PLMN_HUMAN | Plasminogen OS=Homo sapiens GN=PLG PE=1 SV=2 | HUMAN | 26 |
| sp\|P08670\|VIME_HUMAN | Vimentin OS=Homo sapiens GN=VIM PE=1 SV=4 | HUMAN | 25 |
| sp\|P11142\|HSP7C_HUMAN | Heat shock cognate 71 kDa protein OS=Homo sapiens GN=HSPA8 PE=1 SV=1 | HUMAN | 25 |
| sp\|P27797\|CALR_HUMAN | Calreticulin OS=Homo sapiens GN=CALR PE=1 SV=1 | HUMAN | 24 |
| sp\|P68371\|TBB4B_HUMAN | Tubulin beta-4B chain OS=Homo sapiens GN=TUBB4B PE=1 SV=1 | HUMAN | 22 |
| sp\|P07437\|TBB5_HUMAN | Tubulin beta chain OS=Homo sapiens GN=TUBB PE=1 SV=2 | HUMAN | 21 |
| sp\|P07237\|PDIA1_HUMAN | Protein disulfide-isomerase OS=Homo sapiens GN=P4HB PE=1 SV=3 | HUMAN | 20 |
| sp\|P01903\|DRA_HUMAN | HLA class II histocompatibility antigen, DR alpha chain OS=Homo sapiens GN=HLA-DRA PE=1 SV=1 | HUMAN | 20 |
| sp\|P0C0L5\|CO4B_HUMAN | Complement C4-B OS=Homo sapiens GN=C4B PE=1 SV=2 | HUMAN | 19 |
| sp\|P0C0L4\|CO4A_HUMAN | Complement C4-A OS=Homo sapiens GN=C4A PE=1 SV=2 | HUMAN | 19 |
| sp\|P14625\|ENPL_HUMAN | Endoplasmin OS=Homo sapiens GN=HSP90B1 PE=1 SV=1 | HUMAN | 19 |
| sp\|P67936\|TPM4_HUMAN | Tropomyosin alpha-4 chain OS=Homo sapiens GN=TPM4 PE=1 SV=3 | HUMAN | 19 |
| sp\|Q9BQE3\|TBA1C_HUMAN | Tubulin alpha-1C chain OS=Homo sapiens GN=TUBA1C PE=1 SV=1 | HUMAN | 19 |
| sp\|P07988\|PSPB_HUMAN | Pulmonary surfactant-associated protein B OS=Homo sapiens GN=SFTPB PE=1 SV=3 | HUMAN | 19 |
| sp\|P68363\|TBA1B_HUMAN | Tubulin alpha-1B chain OS=Homo sapiens GN=TUBA1B PE=1 SV=1 | HUMAN | 19 |
| sp\|Q71U36\|TBA1A_HUMAN | Tubulin alpha-1A chain OS=Homo sapiens GN=TUBA1A PE=1 SV=1 | HUMAN | 18 |
| sp\|P68366\|TBA4A_HUMAN | Tubulin alpha-4A chain OS=Homo sapiens GN=TUBA4A PE=1 SV=1 | HUMAN | 15 |
| sp\|P02766\|TTHY_HUMAN | Transthyretin OS=Homo sapiens GN=TTR PE=1 SV=1 | HUMAN | 14 |
| sp\|P06753\|TPM3_HUMAN | Tropomyosin alpha-3 chain OS=Homo sapiens GN=TPM3 PE=1 SV=2 | HUMAN | 14 |
| sp\|P06727\|APOA4_HUMAN | Apolipoprotein A-IV OS=Homo sapiens GN=APOA4 PE=1 SV=3 | HUMAN | 13 |
| sp\|P07355\|ANXA2_HUMAN | Annexin A2 OS=Homo sapiens GN=ANXA2 PE=1 SV=2 | HUMAN | 13 |
| sp\|Q02818\|NUCB1_HUMAN | Nucleobindin-1 OS=Homo sapiens GN=NUCB1 PE=1 SV=4 | HUMAN | 13 |
| sp\|P63261\|ACTG_HUMAN | Actin, cytoplasmic 2 OS=Homo sapiens GN=ACTG1 PE=1 SV=1 | HUMAN | 13 |
| sp\|P60709\|ACTB_HUMAN | Actin, cytoplasmic 1 OS=Homo sapiens GN=ACTB PE=1 SV=1 | HUMAN | 13 |
| sp\|P62258\|1433E_HUMAN | 14-3-3 protein epsilon OS=Homo sapiens GN=YWHAE PE=1 SV=1 | HUMAN | 13 |
| sp\|P14314\|GLU2B_HUMAN | Glucosidase 2 subunit beta OS=Homo sapiens GN=PRKCSH PE=1 SV=2 | HUMAN | 13 |
| sp\|P02786\|TFR1_HUMAN | Transferrin receptor protein 1 OS=Homo sapiens GN=TFRC PE=1 SV=2 | HUMAN | 12 |
| sp\|Q5Y7A7\|2B1D_HUMAN | HLA class II histocompatibility antigen, DRB1-13 beta chain OS=Homo sapiens GN=HLA-DRB1 PE=1 SV=1 | HUMAN | 12 |
| sp\|O00560\|SDCB1_HUMAN | Syntenin-1 OS=Homo sapiens GN=SDCBP PE=1 SV=1 | HUMAN | 12 |
| sp\|P52566\|GDIR2_HUMAN | Rho GDP-dissociation inhibitor 2 OS=Homo sapiens GN=ARHGDIB PE=1 SV=3 | HUMAN | 12 |
| sp\|P13667\|PDIA4_HUMAN | Protein disulfide-isomerase A4 OS=Homo sapiens GN=PDIA4 PE=1 SV=2 | HUMAN | 12 |
| sp\|P63104\|1433Z_HUMAN | 14-3-3 protein zeta/delta OS=Homo sapiens GN=YWHAZ PE=1 SV=1 | HUMAN | 12 |
| sp\|P79483\|DRB3_HUMAN | HLA class II histocompatibility antigen, DR beta 3 chain OS=Homo sapiens GN=HLA-DRB3 PE=1 SV=1 | HUMAN | 12 |
| sp\|P02654\|APOC1_HUMAN | Apolipoprotein C-I OS=Homo sapiens GN=APOC1 PE=1 SV=1 | HUMAN | 12 |
| sp\|P08493\|MGP_HUMAN | Matrix Gla protein OS=Homo sapiens GN=MGP PE=1 SV=2 | HUMAN | 12 |
| sp\|P35579\|MYH9_HUMAN | Myosin-9 OS=Homo sapiens GN=MYH9 PE=1 SV=4 | HUMAN | 11 |
| sp\|P33241\|LSP1_HUMAN | Lymphocyte-specific protein 1 OS=Homo sapiens GN=LSP1 PE=1 SV=1 | HUMAN | 11 |
| sp\|P09429\|HMGB1_HUMAN | High mobility group protein B1 OS=Homo sapiens GN=HMGB1 PE=1 SV=3 | HUMAN | 11 |
| sp\|P05387\|RLA2_HUMAN | 60S acidic ribosomal protein P2 OS=Homo sapiens GN=RPLP2 PE=1 SV=1 | HUMAN | 11 |
| sp\|P06702\|S10A9_HUMAN | Protein S100-A9 OS=Homo sapiens GN=S100A9 PE=1 SV=1 | HUMAN | 11 |
| sp\|P02748\|CO9_HUMAN | Complement component C9 OS=Homo sapiens GN=C9 PE=1 SV=2 | HUMAN | 10 |
| sp\|P62158\|CALM_HUMAN | Calmodulin OS=Homo sapiens GN=CALM1 PE=1 SV=2 | HUMAN | 10 |
| sp\|P25774\|CATS_HUMAN | Cathepsin S OS=Homo sapiens GN=CTSS PE=1 SV=3 | HUMAN | 9 |
| sp\|P09668\|CATH_HUMAN | Pro-cathepsin H OS=Homo sapiens GN=CTSH PE=1 SV=4 | HUMAN | 9 |
| sp\|P19338\|NUCL_HUMAN | Nucleolin OS=Homo sapiens GN=NCL PE=1 SV=3 | HUMAN | 8 |
| sp\|Q6NZI2\|PTRF_HUMAN | Polymerase I and transcript release factor OS=Homo sapiens GN=PTRF PE=1 SV=1 | HUMAN | 8 |
| sp\|P61769\|B2MG_HUMAN | Beta-2-microglobulin OS=Homo sapiens GN=B2M PE=1 SV=1 | HUMAN | 8 |
| sp\|Q99880\|H2B1L_HUMAN | Histone H2B type 1-L OS=Homo sapiens GN=HIST1H2BL PE=1 SV=3 | HUMAN | 8 |
| sp\|Q99879\|H2B1M_HUMAN | Histone H2B type 1-M OS=Homo sapiens GN=HIST1H2BM PE=1 SV=3 | HUMAN | 8 |
| sp\|Q99877\|H2B1N_HUMAN | Histone H2B type 1-N OS=Homo sapiens GN=HIST1H2BN PE=1 SV=3 | HUMAN | 8 |

| | | | |
|---|---|---|---|
| sp\|Q93079\|H2B1H_HUMAN | Histone H2B type 1-H OS=Homo sapiens GN=HIST1H2BH PE=1 SV=3 | HUMAN | 8 |
| sp\|Q5QNW6\|H2B2F_HUMAN | Histone H2B type 2-F OS=Homo sapiens GN=HIST2H2BF PE=1 SV=3 | HUMAN | 8 |
| sp\|P62807\|H2B1C_HUMAN | Histone H2B type 1-C/E/F/G/I OS=Homo sapiens GN=HIST1H2BC PE=1 SV=4 | HUMAN | 8 |
| sp\|P58876\|H2B1D_HUMAN | Histone H2B type 1-D OS=Homo sapiens GN=HIST1H2BD PE=1 SV=2 | HUMAN | 8 |
| sp\|P57053\|H2BFS_HUMAN | Histone H2B type F-S OS=Homo sapiens GN=H2BFS PE=1 SV=2 | HUMAN | 8 |
| sp\|O60814\|H2B1K_HUMAN | Histone H2B type 1-K OS=Homo sapiens GN=HIST1H2BK PE=1 SV=3 | HUMAN | 8 |
| sp\|P15311\|EZRI_HUMAN | Ezrin OS=Homo sapiens GN=EZR PE=1 SV=4 | HUMAN | 8 |
| sp\|P01857\|IGHG1_HUMAN | Ig gamma-1 chain C region OS=Homo sapiens GN=IGHG1 PE=1 SV=1 | HUMAN | 7 |
| sp\|P23284\|PPIB_HUMAN | Peptidyl-prolyl cis-trans isomerase B OS=Homo sapiens GN=PPIB PE=1 SV=2 | HUMAN | 7 |
| sp\|P51858\|HDGF_HUMAN | Hepatoma-derived growth factor OS=Homo sapiens GN=HDGF PE=1 SV=1 | HUMAN | 7 |
| sp\|P0DMV9\|HS71B_HUMAN | Heat shock 70 kDa protein 1B OS=Homo sapiens GN=HSPA1B PE=1 SV=1 | HUMAN | 7 |
| sp\|P0DMV8\|HS71A_HUMAN | Heat shock 70 kDa protein 1A OS=Homo sapiens GN=HSPA1A PE=1 SV=1 | HUMAN | 7 |
| sp\|Q93077\|H2A1C_HUMAN | Histone H2A type 1-C OS=Homo sapiens GN=HIST1H2AC PE=1 SV=3 | HUMAN | 7 |
| sp\|Q7L7L0\|H2A3_HUMAN | Histone H2A type 3 OS=Homo sapiens GN=HIST3H2A PE=1 SV=3 | HUMAN | 7 |
| sp\|P04908\|H2A1B_HUMAN | Histone H2A type 1-B/E OS=Homo sapiens GN=HIST1H2AB PE=1 SV=2 | HUMAN | 7 |
| sp\|Q9GIY3\|2B1E_HUMAN | HLA class II histocompatibility antigen, DRB1-14 beta chain OS=Homo sapiens GN=HLA-DRB1 PE=1 SV=1 | HUMAN | 7 |
| sp\|P09493\|TPM1_HUMAN | Tropomyosin alpha-1 chain OS=Homo sapiens GN=TPM1 PE=1 SV=2 | HUMAN | 7 |
| sp\|P62805\|H4_HUMAN | Histone H4 OS=Homo sapiens GN=HIST1H4A PE=1 SV=2 | HUMAN | 6 |
| sp\|O96009\|NAPSA_HUMAN | Napsin-A OS=Homo sapiens GN=NAPSA PE=1 SV=1 | HUMAN | 6 |
| sp\|P07225\|PROS_HUMAN | Vitamin K-dependent protein S OS=Homo sapiens GN=PROS1 PE=1 SV=1 | HUMAN | 6 |
| sp\|P01009\|A1AT_HUMAN | Alpha-1-antitrypsin OS=Homo sapiens GN=SERPINA1 PE=1 SV=3 | HUMAN | 6 |
| sp\|P27824\|CALX_HUMAN | Calnexin OS=Homo sapiens GN=CANX PE=1 SV=2 | HUMAN | 6 |
| sp\|P60660\|MYL6_HUMAN | Myosin light polypeptide 6 OS=Homo sapiens GN=MYL6 PE=1 SV=2 | HUMAN | 6 |
| sp\|P50502\|F10A1_HUMAN | Hsc70-interacting protein OS=Homo sapiens GN=ST13 PE=1 SV=2 | HUMAN | 6 |
| sp\|P61916\|NPC2_HUMAN | Epididymal secretory protein E1 OS=Homo sapiens GN=NPC2 PE=1 SV=1 | HUMAN | 6 |
| sp\|P62987\|RL40_HUMAN | Ubiquitin-60S ribosomal protein L40 OS=Homo sapiens GN=UBA52 PE=1 SV=2 | HUMAN | 6 |
| sp\|P62979\|RS27A_HUMAN | Ubiquitin-40S ribosomal protein S27a OS=Homo sapiens GN=RPS27A PE=1 SV=2 | HUMAN | 6 |
| sp\|P0CG48\|UBC_HUMAN | Polyubiquitin-C OS=Homo sapiens GN=UBC PE=1 SV=3 | HUMAN | 6 |
| sp\|P0CG47\|UBB_HUMAN | Polyubiquitin-B OS=Homo sapiens GN=UBB PE=1 SV=1 | HUMAN | 6 |
| sp\|P04003\|C4BPA_HUMAN | C4b-binding protein alpha chain OS=Homo sapiens GN=C4BPA PE=1 SV=2 | HUMAN | 6 |
| sp\|Q9BTM1\|H2AJ_HUMAN | Histone H2A.J OS=Homo sapiens GN=H2AFJ PE=1 SV=1 | HUMAN | 6 |
| sp\|Q99878\|H2A1J_HUMAN | Histone H2A type 1-J OS=Homo sapiens GN=HIST1H2AJ PE=1 SV=3 | HUMAN | 6 |
| sp\|Q96KK5\|H2A1H_HUMAN | Histone H2A type 1-H OS=Homo sapiens GN=HIST1H2AH PE=1 SV=3 | HUMAN | 6 |
| sp\|Q6FI13\|H2A2A_HUMAN | Histone H2A type 2-A OS=Homo sapiens GN=HIST2H2AA3 PE=1 SV=3 | HUMAN | 6 |
| sp\|Q16777\|H2A2C_HUMAN | Histone H2A type 2-C OS=Homo sapiens GN=HIST2H2AC PE=1 SV=4 | HUMAN | 6 |
| sp\|P20671\|H2A1D_HUMAN | Histone H2A type 1-D OS=Homo sapiens GN=HIST1H2AD PE=1 SV=2 | HUMAN | 6 |
| sp\|P0C0S8\|H2A1_HUMAN | Histone H2A type 1 OS=Homo sapiens GN=HIST1H2AG PE=1 SV=2 | HUMAN | 6 |
| sp\|P16104\|H2AX_HUMAN | Histone H2AX OS=Homo sapiens GN=H2AFX PE=1 SV=2 | HUMAN | 6 |
| sp\|P01861\|IGHG4_HUMAN | Ig gamma-4 chain C region OS=Homo sapiens GN=IGHG4 PE=1 SV=1 | HUMAN | 6 |
| sp\|P30457\|1A66_HUMAN | HLA class I histocompatibility antigen, A-66 alpha chain OS=Homo sapiens GN=HLA-A PE=1 SV=1 | HUMAN | 5 |
| sp\|P30453\|1A34_HUMAN | HLA class I histocompatibility antigen, A-34 alpha chain OS=Homo sapiens GN=HLA-A PE=1 SV=1 | HUMAN | 5 |
| sp\|P30450\|1A26_HUMAN | HLA class I histocompatibility antigen, A-26 alpha chain OS=Homo sapiens GN=HLA-A PE=1 SV=2 | HUMAN | 5 |
| sp\|P18462\|1A25_HUMAN | HLA class I histocompatibility antigen, A-25 alpha chain OS=Homo sapiens GN=HLA-A PE=1 SV=1 | HUMAN | 5 |
| sp\|Q13838\|DX39B_HUMAN | Spliceosome RNA helicase DDX39B OS=Homo sapiens GN=DDX39B PE=1 SV=1 | HUMAN | 5 |
| sp\|P08294\|SODE_HUMAN | Extracellular superoxide dismutase [Cu-Zn] OS=Homo sapiens GN=SOD3 PE=1 SV=2 | HUMAN | 5 |
| sp\|P06748\|NPM_HUMAN | Nucleophosmin OS=Homo sapiens GN=NPM1 PE=1 SV=2 | HUMAN | 5 |
| sp\|P02787\|TRFE_HUMAN | Serotransferrin OS=Homo sapiens GN=TF PE=1 SV=3 | HUMAN | 5 |
| sp\|Q8NFI4\|F10A5_HUMAN | Putative protein FAM10A5 OS=Homo sapiens GN=ST13P5 PE=5 SV=1 | HUMAN | 5 |
| sp\|P04406\|G3P_HUMAN | Glyceraldehyde-3-phosphate dehydrogenase OS=Homo sapiens GN=GAPDH PE=1 SV=3 | HUMAN | 5 |
| sp\|P50995\|ANX11_HUMAN | Annexin A11 OS=Homo sapiens GN=ANXA11 PE=1 SV=1 | HUMAN | 5 |
| sp\|P04083\|ANXA1_HUMAN | Annexin A1 OS=Homo sapiens GN=ANXA1 PE=1 SV=2 | HUMAN | 5 |
| sp\|P01834\|IGKC_HUMAN | Ig kappa chain C region OS=Homo sapiens GN=IGKC PE=1 SV=1 | HUMAN | 5 |
| sp\|P07339\|CATD_HUMAN | Cathepsin D OS=Homo sapiens GN=CTSD PE=1 SV=1 | HUMAN | 5 |
| sp\|P01920\|DQB1_HUMAN | HLA class II histocompatibility antigen, DQ beta 1 chain OS=Homo sapiens GN=HLA-DQB1 PE=1 SV=2 | HUMAN | 5 |
| sp\|P19105\|ML12A_HUMAN | Myosin regulatory light chain 12A OS=Homo sapiens GN=MYL12A PE=1 SV=2 | HUMAN | 5 |
| sp\|O14950\|ML12B_HUMAN | Myosin regulatory light chain 12B OS=Homo sapiens GN=MYL12B PE=1 SV=2 | HUMAN | 5 |
| sp\|P05109\|S10A8_HUMAN | Protein S100-A8 OS=Homo sapiens GN=S100A8 PE=1 SV=1 | HUMAN | 5 |
| sp\|Q92688\|AN32B_HUMAN | Acidic leucine-rich nuclear phosphoprotein 32 family member B OS=Homo sapiens GN=ANP32B PE=1 SV= | HUMAN | 5 |
| sp\|P02774\|VTDB_HUMAN | Vitamin D-binding protein OS=Homo sapiens GN=GC PE=1 SV=1 | HUMAN | 5 |
| sp\|P07602\|SAP_HUMAN | Prosaposin OS=Homo sapiens GN=PSAP PE=1 SV=2 | HUMAN | 5 |
| sp\|P31947\|1433S_HUMAN | 14-3-3 protein sigma OS=Homo sapiens GN=SFN PE=1 SV=1 | HUMAN | 5 |
| sp\|P61981\|1433G_HUMAN | 14-3-3 protein gamma OS=Homo sapiens GN=YWHAG PE=1 SV=2 | HUMAN | 5 |
| sp\|P02810\|PRPC_HUMAN | Salivary acidic proline-rich phosphoprotein 1/2 OS=Homo sapiens GN=PRH1 PE=1 SV=2 | HUMAN | 5 |
| sp\|P16190\|1A33_HUMAN | HLA class I histocompatibility antigen, A-33 alpha chain OS=Homo sapiens GN=HLA-A PE=1 SV=3 | HUMAN | 4 |
| sp\|P10316\|1A69_HUMAN | HLA class I histocompatibility antigen, A-69 alpha chain OS=Homo sapiens GN=HLA-A PE=1 SV=2 | HUMAN | 4 |
| sp\|P30459\|1A74_HUMAN | HLA class I histocompatibility antigen, A-74 alpha chain OS=Homo sapiens GN=HLA-A PE=1 SV=1 | HUMAN | 4 |
| sp\|P30456\|1A43_HUMAN | HLA class I histocompatibility antigen, A-43 alpha chain OS=Homo sapiens GN=HLA-A PE=1 SV=1 | HUMAN | 4 |
| sp\|P16189\|1A31_HUMAN | HLA class I histocompatibility antigen, A-31 alpha chain OS=Homo sapiens GN=HLA-A PE=1 SV=2 | HUMAN | 4 |
| sp\|P10314\|1A32_HUMAN | HLA class I histocompatibility antigen, A-32 alpha chain OS=Homo sapiens GN=HLA-A PE=1 SV=2 | HUMAN | 4 |
| sp\|P01891\|1A68_HUMAN | HLA class I histocompatibility antigen, A-68 alpha chain OS=Homo sapiens GN=HLA-A PE=1 SV=4 | HUMAN | 4 |
| sp\|O00148\|DX39A_HUMAN | ATP-dependent RNA helicase DDX39A OS=Homo sapiens GN=DDX39A PE=1 SV=2 | HUMAN | 4 |

| | | | |
|---|---|---|---|
| sp\|Q8WUM4\|PDC6I_HUMAN | Programmed cell death 6-interacting protein OS=Homo sapiens GN=PDCD6IP PE=1 SV=1 | HUMAN | 4 |
| sp\|P53634\|CATC_HUMAN | Dipeptidyl peptidase 1 OS=Homo sapiens GN=CTSC PE=1 SV=2 | HUMAN | 4 |
| sp\|O75487\|GPC4_HUMAN | Glypican-4 OS=Homo sapiens GN=GPC4 PE=1 SV=4 | HUMAN | 4 |
| sp\|O60462\|NRP2_HUMAN | Neuropilin-2 OS=Homo sapiens GN=NRP2 PE=1 SV=2 | HUMAN | 4 |
| sp\|P02751\|FINC_HUMAN | Fibronectin OS=Homo sapiens GN=FN1 PE=1 SV=4 | HUMAN | 4 |
| sp\|P16035\|TIMP2_HUMAN | Metalloproteinase inhibitor 2 OS=Homo sapiens GN=TIMP2 PE=1 SV=2 | HUMAN | 4 |
| sp\|P61626\|LYSC_HUMAN | Lysozyme C OS=Homo sapiens GN=LYZ PE=1 SV=1 | HUMAN | 4 |
| sp\|Q86VP6\|CAND1_HUMAN | Cullin-associated NEDD8-dissociated protein 1 OS=Homo sapiens GN=CAND1 PE=1 SV=2 | HUMAN | 4 |
| sp\|P00740\|FA9_HUMAN | Coagulation factor IX OS=Homo sapiens GN=F9 PE=1 SV=2 | HUMAN | 4 |
| sp\|Q16543\|CDC37_HUMAN | Hsp90 co-chaperone Cdc37 OS=Homo sapiens GN=CDC37 PE=1 SV=1 | HUMAN | 4 |
| sp\|P02656\|APOC3_HUMAN | Apolipoprotein C-III OS=Homo sapiens GN=APOC3 PE=1 SV=1 | HUMAN | 4 |
| sp\|P39687\|AN32A_HUMAN | Acidic leucine-rich nuclear phosphoprotein 32 family member A OS=Homo sapiens GN=ANP32A PE=1 SV= | HUMAN | 4 |
| sp\|P60033\|CD81_HUMAN | CD81 antigen OS=Homo sapiens GN=CD81 PE=1 SV=1 | HUMAN | 4 |
| sp\|Q04826\|1B40_HUMAN | HLA class I histocompatibility antigen, B-40 alpha chain OS=Homo sapiens GN=HLA-B PE=1 SV=1 | HUMAN | 4 |
| sp\|Q95365\|1B38_HUMAN | HLA class I histocompatibility antigen, B-38 alpha chain OS=Homo sapiens GN=HLA-B PE=1 SV=1 | HUMAN | 4 |
| sp\|Q29940\|1B59_HUMAN | HLA class I histocompatibility antigen, B-59 alpha chain OS=Homo sapiens GN=HLA-B PE=1 SV=1 | HUMAN | 4 |
| sp\|Q29836\|1B67_HUMAN | HLA class I histocompatibility antigen, B-67 alpha chain OS=Homo sapiens GN=HLA-B PE=1 SV=1 | HUMAN | 4 |
| sp\|Q29718\|1B82_HUMAN | HLA class I histocompatibility antigen, B-82 alpha chain OS=Homo sapiens GN=HLA-B PE=1 SV=1 | HUMAN | 4 |
| sp\|P30495\|1B56_HUMAN | HLA class I histocompatibility antigen, B-56 alpha chain OS=Homo sapiens GN=HLA-B PE=1 SV=1 | HUMAN | 4 |
| sp\|P30493\|1B55_HUMAN | HLA class I histocompatibility antigen, B-55 alpha chain OS=Homo sapiens GN=HLA-B PE=1 SV=1 | HUMAN | 4 |
| sp\|P30488\|1B50_HUMAN | HLA class I histocompatibility antigen, B-50 alpha chain OS=Homo sapiens GN=HLA-B PE=1 SV=1 | HUMAN | 4 |
| sp\|P30487\|1B49_HUMAN | HLA class I histocompatibility antigen, B-49 alpha chain OS=Homo sapiens GN=HLA-B PE=1 SV=2 | HUMAN | 4 |
| sp\|P30485\|1B47_HUMAN | HLA class I histocompatibility antigen, B-47 alpha chain OS=Homo sapiens GN=HLA-B PE=1 SV=1 | HUMAN | 4 |
| sp\|P30484\|1B46_HUMAN | HLA class I histocompatibility antigen, B-46 alpha chain OS=Homo sapiens GN=HLA-B PE=1 SV=1 | HUMAN | 4 |
| sp\|P30483\|1B45_HUMAN | HLA class I histocompatibility antigen, B-45 alpha chain OS=Homo sapiens GN=HLA-B PE=1 SV=1 | HUMAN | 4 |
| sp\|P30475\|1B39_HUMAN | HLA class I histocompatibility antigen, B-39 alpha chain OS=Homo sapiens GN=HLA-B PE=1 SV=1 | HUMAN | 4 |
| sp\|P30464\|1B15_HUMAN | HLA class I histocompatibility antigen, B-15 alpha chain OS=Homo sapiens GN=HLA-B PE=1 SV=2 | HUMAN | 4 |
| sp\|P18463\|1B37_HUMAN | HLA class I histocompatibility antigen, B-37 alpha chain OS=Homo sapiens GN=HLA-B PE=1 SV=1 | HUMAN | 4 |
| sp\|P03989\|1B27_HUMAN | HLA class I histocompatibility antigen, B-27 alpha chain OS=Homo sapiens GN=HLA-B PE=1 SV=2 | HUMAN | 4 |
| sp\|P30461\|1B13_HUMAN | HLA class I histocompatibility antigen, B-13 alpha chain OS=Homo sapiens GN=HLA-B PE=1 SV=1 | HUMAN | 4 |
| sp\|P30685\|1B35_HUMAN | HLA class I histocompatibility antigen, B-35 alpha chain OS=Homo sapiens GN=HLA-B PE=1 SV=1 | HUMAN | 4 |
| sp\|P30491\|1B53_HUMAN | HLA class I histocompatibility antigen, B-53 alpha chain OS=Homo sapiens GN=HLA-B PE=1 SV=1 | HUMAN | 4 |
| sp\|P30481\|1B44_HUMAN | HLA class I histocompatibility antigen, B-44 alpha chain OS=Homo sapiens GN=HLA-B PE=1 SV=1 | HUMAN | 4 |
| sp\|O75340\|PDCD6_HUMAN | Programmed cell death protein 6 OS=Homo sapiens GN=PDCD6 PE=1 SV=1 | HUMAN | 4 |
| sp\|Q04917\|1433F_HUMAN | 14-3-3 protein eta OS=Homo sapiens GN=YWHAH PE=1 SV=4 | HUMAN | 4 |
| sp\|P31946\|1433B_HUMAN | 14-3-3 protein beta/alpha OS=Homo sapiens GN=YWHAB PE=1 SV=3 | HUMAN | 4 |
| sp\|P27348\|1433T_HUMAN | 14-3-3 protein theta OS=Homo sapiens GN=YWHAQ PE=1 SV=1 | HUMAN | 4 |
| sp\|P01906\|DQA2_HUMAN | HLA class II histocompatibility antigen, DQ alpha 2 chain OS=Homo sapiens GN=HLA-DQA2 PE=1 SV=2 | HUMAN | 4 |
| sp\|P01859\|IGHG2_HUMAN | Ig gamma-2 chain C region OS=Homo sapiens GN=IGHG2 PE=1 SV=2 | HUMAN | 4 |
| sp\|O75351\|VPS4B_HUMAN | Vacuolar protein sorting-associated protein 4B OS=Homo sapiens GN=VPS4B PE=1 SV=2 | HUMAN | 3 |
| sp\|P59666\|DEF3_HUMAN | Neutrophil defensin 3 OS=Homo sapiens GN=DEFA3 PE=1 SV=1 | HUMAN | 3 |
| sp\|P59665\|DEF1_HUMAN | Neutrophil defensin 1 OS=Homo sapiens GN=DEFA1 PE=1 SV=1 | HUMAN | 3 |
| sp\|Q9NQP4\|PFD4_HUMAN | Prefoldin subunit 4 OS=Homo sapiens GN=PFDN4 PE=1 SV=1 | HUMAN | 3 |
| sp\|P09467\|F16P1_HUMAN | Fructose-1,6-bisphosphatase 1 OS=Homo sapiens GN=FBP1 PE=1 SV=5 | HUMAN | 3 |
| sp\|Q15185\|TEBP_HUMAN | Prostaglandin E synthase 3 OS=Homo sapiens GN=PTGES3 PE=1 SV=1 | HUMAN | 3 |
| sp\|P68871\|HBB_HUMAN | Hemoglobin subunit beta OS=Homo sapiens GN=HBB PE=1 SV=2 | HUMAN | 3 |
| sp\|P29692\|EF1D_HUMAN | Elongation factor 1-delta OS=Homo sapiens GN=EEF1D PE=1 SV=5 | HUMAN | 3 |
| sp\|P0CG06\|LAC3_HUMAN | Ig lambda-3 chain C regions OS=Homo sapiens GN=IGLC3 PE=1 SV=1 | HUMAN | 3 |
| sp\|P0CG05\|LAC2_HUMAN | Ig lambda-2 chain C regions OS=Homo sapiens GN=IGLC2 PE=1 SV=1 | HUMAN | 3 |
| sp\|P0CF74\|LAC6_HUMAN | Ig lambda-6 chain C region OS=Homo sapiens GN=IGLC6 PE=4 SV=1 | HUMAN | 3 |
| sp\|Q9UJU6\|DBNL_HUMAN | Drebrin-like protein OS=Homo sapiens GN=DBNL PE=1 SV=1 | HUMAN | 3 |
| sp\|P10909\|CLUS_HUMAN | Clusterin OS=Homo sapiens GN=CLU PE=1 SV=1 | HUMAN | 3 |
| sp\|O43396\|TXNL1_HUMAN | Thioredoxin-like protein 1 OS=Homo sapiens GN=TXNL1 PE=1 SV=3 | HUMAN | 3 |
| sp\|P62993\|GRB2_HUMAN | Growth factor receptor-bound protein 2 OS=Homo sapiens GN=GRB2 PE=1 SV=1 | HUMAN | 3 |
| sp\|P21926\|CD9_HUMAN | CD9 antigen OS=Homo sapiens GN=CD9 PE=1 SV=4 | HUMAN | 3 |
| sp\|P63000\|RAC1_HUMAN | Ras-related C3 botulinum toxin substrate 1 OS=Homo sapiens GN=RAC1 PE=1 SV=1 | HUMAN | 3 |
| sp\|P15153\|RAC2_HUMAN | Ras-related C3 botulinum toxin substrate 2 OS=Homo sapiens GN=RAC2 PE=1 SV=1 | HUMAN | 3 |
| sp\|P60763\|RAC3_HUMAN | Ras-related C3 botulinum toxin substrate 3 OS=Homo sapiens GN=RAC3 PE=1 SV=1 | HUMAN | 3 |
| sp\|Q9NQC3\|RTN4_HUMAN | Reticulon-4 OS=Homo sapiens GN=RTN4 PE=1 SV=2 | HUMAN | 3 |
| sp\|P14317\|HCLS1_HUMAN | Hematopoietic lineage cell-specific protein OS=Homo sapiens GN=HCLS1 PE=1 SV=3 | HUMAN | 3 |
| sp\|Q9Y5S9\|RBM8A_HUMAN | RNA-binding protein 8A OS=Homo sapiens GN=RBM8A PE=1 SV=1 | HUMAN | 3 |
| sp\|P60983\|GMFB_HUMAN | Glia maturation factor beta OS=Homo sapiens GN=GMFB PE=1 SV=2 | HUMAN | 3 |
| sp\|Q14520\|HABP2_HUMAN | Hyaluronan-binding protein 2 OS=Homo sapiens GN=HABP2 PE=1 SV=1 | HUMAN | 3 |
| sp\|P63208\|SKP1_HUMAN | S-phase kinase-associated protein 1 OS=Homo sapiens GN=SKP1 PE=1 SV=2 | HUMAN | 3 |
| sp\|P30480\|1B42_HUMAN | HLA class I histocompatibility antigen, B-42 alpha chain OS=Homo sapiens GN=HLA-B PE=1 SV=1 | HUMAN | 3 |
| sp\|P30479\|1B41_HUMAN | HLA class I histocompatibility antigen, B-41 alpha chain OS=Homo sapiens GN=HLA-B PE=1 SV=1 | HUMAN | 3 |
| sp\|P30460\|1B08_HUMAN | HLA class I histocompatibility antigen, B-8 alpha chain OS=Homo sapiens GN=HLA-B PE=1 SV=1 | HUMAN | 3 |
| sp\|P01889\|1B07_HUMAN | HLA class I histocompatibility antigen, B-7 alpha chain OS=Homo sapiens GN=HLA-B PE=1 SV=3 | HUMAN | 3 |
| sp\|P30466\|1B18_HUMAN | HLA class I histocompatibility antigen, B-18 alpha chain OS=Homo sapiens GN=HLA-B PE=1 SV=1 | HUMAN | 3 |
| sp\|P30462\|1B14_HUMAN | HLA class I histocompatibility antigen, B-14 alpha chain OS=Homo sapiens GN=HLA-B PE=1 SV=1 | HUMAN | 3 |
| sp\|P30492\|1B54_HUMAN | HLA class I histocompatibility antigen, B-54 alpha chain OS=Homo sapiens GN=HLA-B PE=1 SV=1 | HUMAN | 3 |

| | | | |
|---|---|---|---|
| sp\|P18465\|1B57_HUMAN | HLA class I histocompatibility antigen, B-57 alpha chain OS=Homo sapiens GN=HLA-B PE=1 SV=1 | HUMAN | 3 |
| sp\|P30498\|1B78_HUMAN | HLA class I histocompatibility antigen, B-78 alpha chain OS=Homo sapiens GN=HLA-B PE=1 SV=1 | HUMAN | 3 |
| sp\|P30490\|1B52_HUMAN | HLA class I histocompatibility antigen, B-52 alpha chain OS=Homo sapiens GN=HLA-B PE=1 SV=1 | HUMAN | 3 |
| sp\|P18464\|1B51_HUMAN | HLA class I histocompatibility antigen, B-51 alpha chain OS=Homo sapiens GN=HLA-B PE=1 SV=1 | HUMAN | 3 |
| sp\|P10319\|1B58_HUMAN | HLA class I histocompatibility antigen, B-58 alpha chain OS=Homo sapiens GN=HLA-B PE=1 SV=1 | HUMAN | 3 |
| sp\|Q31610\|1B81_HUMAN | HLA class I histocompatibility antigen, B-81 alpha chain OS=Homo sapiens GN=HLA-B PE=1 SV=1 | HUMAN | 3 |
| sp\|P30486\|1B48_HUMAN | HLA class I histocompatibility antigen, B-48 alpha chain OS=Homo sapiens GN=HLA-B PE=1 SV=1 | HUMAN | 3 |
| sp\|P09525\|ANXA4_HUMAN | Annexin A4 OS=Homo sapiens GN=ANXA4 PE=1 SV=4 | HUMAN | 3 |
| sp\|P23528\|COF1_HUMAN | Cofilin-1 OS=Homo sapiens GN=CFL1 PE=1 SV=3 | HUMAN | 3 |
| sp\|P26583\|HMGB2_HUMAN | High mobility group protein B2 OS=Homo sapiens GN=HMGB2 PE=1 SV=2 | HUMAN | 3 |
| sp\|Q15084\|PDIA6_HUMAN | Protein disulfide-isomerase A6 OS=Homo sapiens GN=PDIA6 PE=1 SV=1 | HUMAN | 3 |
| sp\|P02042\|HBD_HUMAN | Hemoglobin subunit delta OS=Homo sapiens GN=HBD PE=1 SV=2 | HUMAN | 2 |
| sp\|P0CG04\|LAC1_HUMAN | Ig lambda-1 chain C regions OS=Homo sapiens GN=IGLC1 PE=1 SV=1 | HUMAN | 2 |
| sp\|B9A064\|IGLL5_HUMAN | Immunoglobulin lambda-like polypeptide 5 OS=Homo sapiens GN=IGLL5 PE=2 SV=2 | HUMAN | 2 |
| sp\|A0M8Q6\|LAC7_HUMAN | Ig lambda-7 chain C region OS=Homo sapiens GN=IGLC7 PE=4 SV=2 | HUMAN | 2 |
| sp\|O14773\|TPP1_HUMAN | Tripeptidyl-peptidase 1 OS=Homo sapiens GN=TPP1 PE=1 SV=2 | HUMAN | 2 |
| sp\|P16070\|CD44_HUMAN | CD44 antigen OS=Homo sapiens GN=CD44 PE=1 SV=3 | HUMAN | 2 |
| sp\|Q13185\|CBX3_HUMAN | Chromobox protein homolog 3 OS=Homo sapiens GN=CBX3 PE=1 SV=4 | HUMAN | 2 |
| sp\|P08476\|INHBA_HUMAN | Inhibin beta A chain OS=Homo sapiens GN=INHBA PE=1 SV=2 | HUMAN | 2 |
| sp\|P31949\|S10AB_HUMAN | Protein S100-A11 OS=Homo sapiens GN=S100A11 PE=1 SV=2 | HUMAN | 2 |
| sp\|Q8NBP7\|PCSK9_HUMAN | Proprotein convertase subtilisin/kexin type 9 OS=Homo sapiens GN=PCSK9 PE=1 SV=3 | HUMAN | 2 |
| sp\|P50281\|MMP14_HUMAN | Matrix metalloproteinase-14 OS=Homo sapiens GN=MMP14 PE=1 SV=3 | HUMAN | 2 |
| sp\|P17655\|CAN2_HUMAN | Calpain-2 catalytic subunit OS=Homo sapiens GN=CAPN2 PE=1 SV=6 | HUMAN | 2 |
| sp\|P05067\|A4_HUMAN | Amyloid beta A4 protein OS=Homo sapiens GN=APP PE=1 SV=3 | HUMAN | 2 |
| sp\|Q9Y2B0\|CNPY2_HUMAN | Protein canopy homolog 2 OS=Homo sapiens GN=CNPY2 PE=1 SV=1 | HUMAN | 2 |
| sp\|P26447\|S10A4_HUMAN | Protein S100-A4 OS=Homo sapiens GN=S100A4 PE=1 SV=1 | HUMAN | 2 |
| sp\|P80748\|LV321_HUMAN | Immunoglobulin lambda variable 3-21 OS=Homo sapiens GN=IGLV3-21 PE=1 SV=2 | HUMAN | 2 |
| sp\|P53999\|TCP4_HUMAN | Activated RNA polymerase II transcriptional coactivator p15 OS=Homo sapiens GN=SUB1 PE=1 SV=3 | HUMAN | 2 |
| sp\|P49407\|ARRB1_HUMAN | Beta-arrestin-1 OS=Homo sapiens GN=ARRB1 PE=1 SV=2 | HUMAN | 2 |
| sp\|P36955\|PEDF_HUMAN | Pigment epithelium-derived factor OS=Homo sapiens GN=SERPINF1 PE=1 SV=4 | HUMAN | 2 |
| sp\|P31431\|SDC4_HUMAN | Syndecan-4 OS=Homo sapiens GN=SDC4 PE=1 SV=2 | HUMAN | 2 |
| sp\|P17931\|LEG3_HUMAN | Galectin-3 OS=Homo sapiens GN=LGALS3 PE=1 SV=5 | HUMAN | 2 |
| sp\|P08758\|ANXA5_HUMAN | Annexin A5 OS=Homo sapiens GN=ANXA5 PE=1 SV=2 | HUMAN | 2 |
| sp\|P04632\|CPNS1_HUMAN | Calpain small subunit 1 OS=Homo sapiens GN=CAPNS1 PE=1 SV=1 | HUMAN | 2 |
| sp\|P02747\|C1QC_HUMAN | Complement C1q subcomponent subunit C OS=Homo sapiens GN=C1QC PE=1 SV=3 | HUMAN | 2 |
| sp\|P00751\|CFAB_HUMAN | Complement factor B OS=Homo sapiens GN=CFB PE=1 SV=2 | HUMAN | 2 |
| sp\|Q8TDL5\|BPIB1_HUMAN | BPI fold-containing family B member 1 OS=Homo sapiens GN=BPIFB1 PE=1 SV=1 | HUMAN | 2 |
| sp\|Q13765\|NACA_HUMAN | Nascent polypeptide-associated complex subunit alpha OS=Homo sapiens GN=NACA PE=1 SV=1 | HUMAN | 2 |
| sp\|E9PAV3\|NACAM_HUMAN | Nascent polypeptide-associated complex subunit alpha, muscle-specific form OS=Homo sapiens GN=NACA | HUMAN | 2 |
| sp\|P84103\|SRSF3_HUMAN | Serine/arginine-rich splicing factor 3 OS=Homo sapiens GN=SRSF3 PE=1 SV=1 | HUMAN | 2 |
| sp\|P06858\|LIPL_HUMAN | Lipoprotein lipase OS=Homo sapiens GN=LPL PE=1 SV=1 | HUMAN | 2 |
| sp\|Q13510\|ASAH1_HUMAN | Acid ceramidase OS=Homo sapiens GN=ASAH1 PE=1 SV=5 | HUMAN | 2 |
| sp\|Q14697\|GANAB_HUMAN | Neutral alpha-glucosidase AB OS=Homo sapiens GN=GANAB PE=1 SV=3 | HUMAN | 2 |
| sp\|P62937\|PPIA_HUMAN | Peptidyl-prolyl cis-trans isomerase A OS=Homo sapiens GN=PPIA PE=1 SV=2 | HUMAN | 2 |
| sp\|Q01105\|SET_HUMAN | Protein SET OS=Homo sapiens GN=SET PE=1 SV=3 | HUMAN | 2 |
| sp\|P0DME0\|SETLP_HUMAN | Protein SETSIP OS=Homo sapiens GN=SETSIP PE=1 SV=1 | HUMAN | 2 |
| sp\|P26641\|EF1G_HUMAN | Elongation factor 1-gamma OS=Homo sapiens GN=EEF1G PE=1 SV=3 | HUMAN | 2 |
| sp\|P29401\|TKT_HUMAN | Transketolase OS=Homo sapiens GN=TKT PE=1 SV=3 | HUMAN | 2 |
| sp\|Q06830\|PRDX1_HUMAN | Peroxiredoxin-1 OS=Homo sapiens GN=PRDX1 PE=1 SV=1 | HUMAN | 2 |
| sp\|Q9Y5P4\|C43BP_HUMAN | Collagen type IV alpha-3-binding protein OS=Homo sapiens GN=COL4A3BP PE=1 SV=1 | HUMAN | 2 |
| sp\|Q96HN2\|SAHH3_HUMAN | Adenosylhomocysteinase 3 OS=Homo sapiens GN=AHCYL2 PE=1 SV=1 | HUMAN | 2 |
| sp\|O43865\|SAHH2_HUMAN | Adenosylhomocysteinase 2 OS=Homo sapiens GN=AHCYL1 PE=1 SV=2 | HUMAN | 2 |
| sp\|Q9NT62\|ATG3_HUMAN | Ubiquitin-like-conjugating enzyme ATG3 OS=Homo sapiens GN=ATG3 PE=1 SV=1 | HUMAN | 2 |
| sp\|P16403\|H12_HUMAN | Histone H1.2 OS=Homo sapiens GN=HIST1H1C PE=1 SV=2 | HUMAN | 2 |
| sp\|Q9BVG4\|PBDC1_HUMAN | Protein PBDC1 OS=Homo sapiens GN=PBDC1 PE=1 SV=1 | HUMAN | 2 |
| sp\|P17096\|HMGA1_HUMAN | High mobility group protein HMG-I/HMG-Y OS=Homo sapiens GN=HMGA1 PE=1 SV=3 | HUMAN | 2 |
| sp\|P02652\|APOA2_HUMAN | Apolipoprotein A-II OS=Homo sapiens GN=APOA2 PE=1 SV=1 | HUMAN | 2 |
| sp\|P24534\|EF1B_HUMAN | Elongation factor 1-beta OS=Homo sapiens GN=EEF1B2 PE=1 SV=3 | HUMAN | 2 |
| sp\|P98160\|PGBM_HUMAN | Basement membrane-specific heparan sulfate proteoglycan core protein OS=Homo sapiens GN=HSPG2 PE | HUMAN | 2 |
| sp\|P21333\|FLNA_HUMAN | Filamin-A OS=Homo sapiens GN=FLNA PE=1 SV=4 | HUMAN | 2 |
| sp\|P20962\|PTMS_HUMAN | Parathymosin OS=Homo sapiens GN=PTMS PE=1 SV=2 | HUMAN | 2 |
| sp\|P04070\|PROC_HUMAN | Vitamin K-dependent protein C OS=Homo sapiens GN=PROC PE=1 SV=1 | HUMAN | 2 |
| sp\|Q00839\|HNRPU_HUMAN | Heterogeneous nuclear ribonucleoprotein U OS=Homo sapiens GN=HNRNPU PE=1 SV=6 | HUMAN | 2 |
| sp\|P0DMR1\|HNRC4_HUMAN | Heterogeneous nuclear ribonucleoprotein C-like 4 OS=Homo sapiens GN=HNRNPCL4 PE=3 SV=1 | HUMAN | 2 |
| sp\|P07910\|HNRPC_HUMAN | Heterogeneous nuclear ribonucleoproteins C1/C2 OS=Homo sapiens GN=HNRNPC PE=1 SV=4 | HUMAN | 2 |
| sp\|O60812\|HNRC1_HUMAN | Heterogeneous nuclear ribonucleoprotein C-like 1 OS=Homo sapiens GN=HNRNPCL1 PE=1 SV=1 | HUMAN | 2 |
| sp\|B7ZW38\|HNRC3_HUMAN | Heterogeneous nuclear ribonucleoprotein C-like 3 OS=Homo sapiens GN=HNRNPCL3 PE=2 SV=1 | HUMAN | 2 |
| sp\|B2RXH8\|HNRC2_HUMAN | Heterogeneous nuclear ribonucleoprotein C-like 2 OS=Homo sapiens GN=HNRNPCL2 PE=1 SV=1 | HUMAN | 2 |
| sp\|Q14116\|IL18_HUMAN | Interleukin-18 OS=Homo sapiens GN=IL18 PE=1 SV=1 | HUMAN | 2 |
| sp\|P54920\|SNAA_HUMAN | Alpha-soluble NSF attachment protein OS=Homo sapiens GN=NAPA PE=1 SV=3 | HUMAN | 2 |
| sp\|P05388\|RLA0_HUMAN | 60S acidic ribosomal protein P0 OS=Homo sapiens GN=RPLP0 PE=1 SV=1 | HUMAN | 2 |

| | | | |
|---|---|---|---|
| sp\|Q8NHW5\|RLA0L_HUMAN | 60S acidic ribosomal protein P0-like OS=Homo sapiens GN=RPLP0P6 PE=5 SV=1 | HUMAN | 2 |
| sp\|P01876\|IGHA1_HUMAN | Ig alpha-1 chain C region OS=Homo sapiens GN=IGHA1 PE=1 SV=2 | HUMAN | 2 |
| sp\|P04440\|DPB1_HUMAN | HLA class II histocompatibility antigen, DP beta 1 chain OS=Homo sapiens GN=HLA-DPB1 PE=1 SV=1 | HUMAN | 2 |
| sp\|O00757\|F16P2_HUMAN | Fructose-1,6-bisphosphatase isozyme 2 OS=Homo sapiens GN=FBP2 PE=1 SV=2 | HUMAN | 2 |
| sp\|P21757\|MSRE_HUMAN | Macrophage scavenger receptor types I and II OS=Homo sapiens GN=MSR1 PE=1 SV=1 | HUMAN | 2 |
