## Supplemental Table3 for "Novel insights into pulmonary phosphate homeostasis and osteoclastogenesis emerge from the study of pulmonary alveolar microlithiasis"

| Accession | Name | Species | Peptides(95%) |
|---|---|---|---|
| sp\|P10923\|OSTP_MOUSE | Osteopontin OS=Mus musculus GN=Spp1 PE=1 SV=1 | MOUSE | 219 |
| sp\|P29699\|FETUA_MOUSE | Alpha-2-HS-glycoprotein OS=Mus musculus GN=Ahsg PE=1 SV=1 | MOUSE | 139 |
| sp\|P07724\|ALBU_MOUSE | Serum albumin OS=Mus musculus GN=Alb PE=1 SV=3 | MOUSE | 91 |
| sp\|P11499\|HS90B_MOUSE | Heat shock protein HSP 90-beta OS=Mus musculus GN=Hsp90ab1 PE=1 SV=3 | MOUSE | 56 |
| sp\|Q8VDD5\|MYH9_MOUSE | Myosin-9 OS=Mus musculus GN=Myh9 PE=1 SV=4 | MOUSE | 54 |
| sp\|P01027\|CO3_MOUSE | Complement C3 OS=Mus musculus GN=C3 PE=1 SV=3 | MOUSE | 45 |
| sp\|P11087\|CO1A1_MOUSE | Collagen alpha-1(I) chain OS=Mus musculus GN=Col1a1 PE=1 SV=4 | MOUSE | 43 |
| sp\|Q00896\|A1AT3_MOUSE | Alpha-1-antitrypsin 1-3 OS=Mus musculus GN=Serpina1c PE=1 SV=2 | MOUSE | 42 |
| sp\|P07758\|A1AT1_MOUSE | Alpha-1-antitrypsin 1-1 OS=Mus musculus GN=Serpina1a PE=1 SV=4 | MOUSE | 41 |
| sp\|P26041\|MOES_MOUSE | Moesin OS=Mus musculus GN=Msn PE=1 SV=3 | MOUSE | 39 |
| sp\|P07901\|HS90A_MOUSE | Heat shock protein HSP 90-alpha OS=Mus musculus GN=Hsp90aa1 PE=1 SV=4 | MOUSE | 39 |
| sp\|P22599\|A1AT2_MOUSE | Alpha-1-antitrypsin 1-2 OS=Mus musculus GN=Serpina1b PE=1 SV=2 | MOUSE | 36 |
| sp\|P62806\|H4_MOUSE | Histone H4 OS=Mus musculus GN=Hist1h4a PE=1 SV=2 | MOUSE | 35 |
| sp\|Q921I1\|TRFE_MOUSE | Serotransferrin OS=Mus musculus GN=Tf PE=1 SV=1 | MOUSE | 34 |
| sp\|Q8CGP2\|H2B1P_MOUSE | Histone H2B type 1-P OS=Mus musculus GN=Hist1h2bp PE=1 SV=3 | MOUSE | 34 |
| sp\|Q8CGP1\|H2B1K_MOUSE | Histone H2B type 1-K OS=Mus musculus GN=Hist1h2bk PE=1 SV=3 | MOUSE | 34 |
| sp\|Q6ZWY9\|H2B1C_MOUSE | Histone H2B type 1-C/E/G OS=Mus musculus GN=Hist1h2bc PE=1 SV=3 | MOUSE | 34 |
| sp\|Q64525\|H2B2B_MOUSE | Histone H2B type 2-B OS=Mus musculus GN=Hist2h2bb PE=1 SV=3 | MOUSE | 34 |
| sp\|Q64478\|H2B1H_MOUSE | Histone H2B type 1-H OS=Mus musculus GN=Hist1h2bh PE=1 SV=3 | MOUSE | 34 |
| sp\|Q64475\|H2B1B_MOUSE | Histone H2B type 1-B OS=Mus musculus GN=Hist1h2bb PE=1 SV=3 | MOUSE | 34 |
| sp\|P10854\|H2B1M_MOUSE | Histone H2B type 1-M OS=Mus musculus GN=Hist1h2bm PE=1 SV=2 | MOUSE | 34 |
| sp\|P10853\|H2B1F_MOUSE | Histone H2B type 1-F/J/L OS=Mus musculus GN=Hist1h2bf PE=1 SV=2 | MOUSE | 34 |
| sp\|Q61207\|SAP_MOUSE | Prosaposin OS=Mus musculus GN=Psap PE=1 SV=2 | MOUSE | 33 |
| sp\|P60710\|ACTB_MOUSE | Actin, cytoplasmic 1 OS=Mus musculus GN=Actb PE=1 SV=1 | MOUSE | 31 |
| sp\|Q00897\|A1AT4_MOUSE | Alpha-1-antitrypsin 1-4 OS=Mus musculus GN=Serpina1d PE=1 SV=1 | MOUSE | 31 |
| sp\|P20029\|GRP78_MOUSE | 78 kDa glucose-regulated protein OS=Mus musculus GN=Hspa5 PE=1 SV=3 | MOUSE | 30 |
| sp\|P09103\|PDIA1_MOUSE | Protein disulfide-isomerase OS=Mus musculus GN=P4hb PE=1 SV=2 | MOUSE | 28 |
| sp\|P07759\|SPA3K_MOUSE | Serine protease inhibitor A3K OS=Mus musculus GN=Serpina3k PE=1 SV=2 | MOUSE | 28 |
| sp\|P19221\|THRB_MOUSE | Prothrombin OS=Mus musculus GN=F2 PE=1 SV=1 | MOUSE | 27 |
| sp\|P48678\|LMNA_MOUSE | Prelamin-A/C OS=Mus musculus GN=Lmna PE=1 SV=2 | MOUSE | 27 |
| sp\|P11589\|MUP2_MOUSE | Major urinary protein 2 OS=Mus musculus GN=Mup2 PE=1 SV=1 | MOUSE | 27 |
| sp\|P50405\|PSPB_MOUSE | Pulmonary surfactant-associated protein B OS=Mus musculus GN=Sftpb PE=1 SV=1 | MOUSE | 26 |
| sp\|P11588\|MUP1_MOUSE | Major urinary protein 1 OS=Mus musculus GN=Mup1 PE=1 SV=1 | MOUSE | 26 |
| sp\|P27773\|PDIA3_MOUSE | Protein disulfide-isomerase A3 OS=Mus musculus GN=Pdia3 PE=1 SV=2 | MOUSE | 23 |
| sp\|P08113\|ENPL_MOUSE | Endoplasmin OS=Mus musculus GN=Hsp90b1 PE=1 SV=2 | MOUSE | 23 |
| sp\|P63017\|HSP7C_MOUSE | Heat shock cognate 71 kDa protein OS=Mus musculus GN=Hspa8 PE=1 SV=1 | MOUSE | 23 |
| sp\|Q01149\|CO1A2_MOUSE | Collagen alpha-2(I) chain OS=Mus musculus GN=Col1a2 PE=1 SV=2 | MOUSE | 23 |
| sp\|P19788\|MGP_MOUSE | Matrix Gla protein OS=Mus musculus GN=Mgp PE=3 SV=1 | MOUSE | 23 |
| sp\|P99024\|TBB5_MOUSE | Tubulin beta-5 chain OS=Mus musculus GN=Tubb5 PE=1 SV=1 | MOUSE | 22 |
| sp\|P11276\|FINC_MOUSE | Fibronectin OS=Mus musculus GN=Fn1 PE=1 SV=4 | MOUSE | 21 |
| sp\|P35564\|CALX_MOUSE | Calnexin OS=Mus musculus GN=Canx PE=1 SV=1 | MOUSE | 19 |
| sp\|P07356\|ANXA2_MOUSE | Annexin A2 OS=Mus musculus GN=Anxa2 PE=1 SV=2 | MOUSE | 19 |
| sp\|P14211\|CALR_MOUSE | Calreticulin OS=Mus musculus GN=Calr PE=1 SV=1 | MOUSE | 19 |
| sp\|P68372\|TBB4B_MOUSE | Tubulin beta-4B chain OS=Mus musculus GN=Tubb4b PE=1 SV=1 | MOUSE | 19 |
| sp\|P09405\|NUCL_MOUSE | Nucleolin OS=Mus musculus GN=Ncl PE=1 SV=2 | MOUSE | 18 |
| sp\|P20152\|VIME_MOUSE | Vimentin OS=Mus musculus GN=Vim PE=1 SV=3 | MOUSE | 18 |
| sp\|P05213\|TBA1B_MOUSE | Tubulin alpha-1B chain OS=Mus musculus GN=Tuba1b PE=1 SV=2 | MOUSE | 17 |
| sp\|P08226\|APOE_MOUSE | Apolipoprotein E OS=Mus musculus GN=Apoe PE=1 SV=2 | MOUSE | 17 |
| sp\|Q8R1M2\|H2AJ_MOUSE | Histone H2A.J OS=Mus musculus GN=H2afj PE=1 SV=1 | MOUSE | 17 |
| sp\|Q8CGP7\|H2A1K_MOUSE | Histone H2A type 1-K OS=Mus musculus GN=Hist1h2ak PE=1 SV=3 | MOUSE | 17 |
| sp\|Q8CGP6\|H2A1H_MOUSE | Histone H2A type 1-H OS=Mus musculus GN=Hist1h2ah PE=1 SV=3 | MOUSE | 17 |
| sp\|Q8CGP5\|H2A1F_MOUSE | Histone H2A type 1-F OS=Mus musculus GN=Hist1h2af PE=1 SV=3 | MOUSE | 17 |
| sp\|Q8BFU2\|H2A3_MOUSE | Histone H2A type 3 OS=Mus musculus GN=Hist3h2a PE=1 SV=3 | MOUSE | 17 |
| sp\|P22752\|H2A1_MOUSE | Histone H2A type 1 OS=Mus musculus GN=Hist1h2ab PE=1 SV=3 | MOUSE | 17 |
| sp\|Q6GSS7\|H2A2A_MOUSE | Histone H2A type 2-A OS=Mus musculus GN=Hist2h2aa1 PE=1 SV=3 | MOUSE | 17 |
| sp\|Q64523\|H2A2C_MOUSE | Histone H2A type 2-C OS=Mus musculus GN=Hist2h2ac PE=1 SV=3 | MOUSE | 17 |
| sp\|P27661\|H2AX_MOUSE | Histone H2AX OS=Mus musculus GN=H2afx PE=1 SV=2 | MOUSE | 17 |
| sp\|Q00898\|A1AT5_MOUSE | Alpha-1-antitrypsin 1-5 OS=Mus musculus GN=Serpina1e PE=1 SV=1 | MOUSE | 17 |
| sp\|Q61233\|PLSL_MOUSE | Plastin-2 OS=Mus musculus GN=Lcp1 PE=1 SV=4 | MOUSE | 16 |
| sp\|P68373\|TBA1C_MOUSE | Tubulin alpha-1C chain OS=Mus musculus GN=Tuba1c PE=1 SV=1 | MOUSE | 16 |
| sp\|P06683\|CO9_MOUSE | Complement component C9 OS=Mus musculus GN=C9 PE=1 SV=2 | MOUSE | 16 |
| sp\|P29788\|VTNC_MOUSE | Vitronectin OS=Mus musculus GN=Vtn PE=1 SV=2 | MOUSE | 16 |
| sp\|P18242\|CATD_MOUSE | Cathepsin D OS=Mus musculus GN=Ctsd PE=1 SV=1 | MOUSE | 16 |
| sp\|P26350\|PTMA_MOUSE | Prothymosin alpha OS=Mus musculus GN=Ptma PE=1 SV=2 | MOUSE | 16 |

| | | | |
|---|---|---|---|
| sp\|P97429\|ANXA4_MOUSE | Annexin A4 OS=Mus musculus GN=Anxa4 PE=1 SV=4 | MOUSE | 15 |
| sp\|Q91X72\|HEMO_MOUSE | Hemopexin OS=Mus musculus GN=Hpx PE=1 SV=2 | MOUSE | 15 |
| sp\|P07309\|TTHY_MOUSE | Transthyretin OS=Mus musculus GN=Ttr PE=1 SV=1 | MOUSE | 15 |
| sp\|P10605\|CATB_MOUSE | Cathepsin B OS=Mus musculus GN=Ctsb PE=1 SV=2 | MOUSE | 15 |
| sp\|P48036\|ANXA5_MOUSE | Annexin A5 OS=Mus musculus GN=Anxa5 PE=1 SV=1 | MOUSE | 14 |
| sp\|P20060\|HEXB_MOUSE | Beta-hexosaminidase subunit beta OS=Mus musculus GN=Hexb PE=1 SV=2 | MOUSE | 14 |
| sp\|P35700\|PRDX1_MOUSE | Peroxiredoxin-1 OS=Mus musculus GN=Prdx1 PE=1 SV=1 | MOUSE | 14 |
| sp\|P26040\|EZRI_MOUSE | Ezrin OS=Mus musculus GN=Ezr PE=1 SV=3 | MOUSE | 14 |
| sp\|P68368\|TBA4A_MOUSE | Tubulin alpha-4A chain OS=Mus musculus GN=Tuba4a PE=1 SV=1 | MOUSE | 14 |
| sp\|O35639\|ANXA3_MOUSE | Annexin A3 OS=Mus musculus GN=Anxa3 PE=1 SV=4 | MOUSE | 13 |
| sp\|P10107\|ANXA1_MOUSE | Annexin A1 OS=Mus musculus GN=Anxa1 PE=1 SV=2 | MOUSE | 13 |
| sp\|P04186\|CFAB_MOUSE | Complement factor B OS=Mus musculus GN=Cfb PE=1 SV=2 | MOUSE | 13 |
| sp\|P32261\|ANT3_MOUSE | Antithrombin-III OS=Mus musculus GN=Serpinc1 PE=1 SV=1 | MOUSE | 13 |
| sp\|Q00623\|APOA1_MOUSE | Apolipoprotein A-I OS=Mus musculus GN=Apoa1 PE=1 SV=2 | MOUSE | 13 |
| sp\|P08003\|PDIA4_MOUSE | Protein disulfide-isomerase A4 OS=Mus musculus GN=Pdia4 PE=1 SV=3 | MOUSE | 13 |
| sp\|P02088\|HBB1_MOUSE | Hemoglobin subunit beta-1 OS=Mus musculus GN=Hbb-b1 PE=1 SV=2 | MOUSE | 13 |
| sp\|Q61362\|CH3L1_MOUSE | Chitinase-3-like protein 1 OS=Mus musculus GN=Chi3l1 PE=1 SV=3 | MOUSE | 13 |
| sp\|P08074\|CBR2_MOUSE | Carbonyl reductase [NADPH] 2 OS=Mus musculus GN=Cbr2 PE=1 SV=1 | MOUSE | 13 |
| sp\|P30681\|HMGB2_MOUSE | High mobility group protein B2 OS=Mus musculus GN=Hmgb2 PE=1 SV=3 | MOUSE | 13 |
| sp\|P08905\|LYZ2_MOUSE | Lysozyme C-2 OS=Mus musculus GN=Lyz2 PE=1 SV=2 | MOUSE | 13 |
| sp\|P14824\|ANXA6_MOUSE | Annexin A6 OS=Mus musculus GN=Anxa6 PE=1 SV=3 | MOUSE | 12 |
| sp\|P58252\|EF2_MOUSE | Elongation factor 2 OS=Mus musculus GN=Eef2 PE=1 SV=2 | MOUSE | 12 |
| sp\|P99027\|RLA2_MOUSE | 60S acidic ribosomal protein P2 OS=Mus musculus GN=Rplp2 PE=1 SV=3 | MOUSE | 12 |
| sp\|P13020\|GELS_MOUSE | Gelsolin OS=Mus musculus GN=Gsn PE=1 SV=3 | MOUSE | 11 |
| sp\|Q6PDM2\|SRSF1_MOUSE | Serine/arginine-rich splicing factor 1 OS=Mus musculus GN=Srsf1 PE=1 SV=3 | MOUSE | 11 |
| sp\|P35242\|SFTPA_MOUSE | Pulmonary surfactant-associated protein A OS=Mus musculus GN=Sftpa1 PE=1 SV=1 | MOUSE | 11 |
| sp\|P84228\|H32_MOUSE | Histone H3.2 OS=Mus musculus GN=Hist1h3b PE=1 SV=2 | MOUSE | 11 |
| sp\|P68433\|H31_MOUSE | Histone H3.1 OS=Mus musculus GN=Hist1h3a PE=1 SV=2 | MOUSE | 11 |
| sp\|P11835\|ITB2_MOUSE | Integrin beta-2 OS=Mus musculus GN=Itgb2 PE=1 SV=2 | MOUSE | 10 |
| sp\|Q61937\|NPM_MOUSE | Nucleophosmin OS=Mus musculus GN=Npm1 PE=1 SV=1 | MOUSE | 10 |
| sp\|P50404\|SFTPD_MOUSE | Pulmonary surfactant-associated protein D OS=Mus musculus GN=Sftpd PE=1 SV=1 | MOUSE | 10 |
| sp\|Q99P91\|GPNMB_MOUSE | Transmembrane glycoprotein NMB OS=Mus musculus GN=Gpnmb PE=1 SV=2 | MOUSE | 10 |
| sp\|P31725\|S10A9_MOUSE | Protein S100-A9 OS=Mus musculus GN=S100a9 PE=1 SV=3 | MOUSE | 10 |
| sp\|Q922R8\|PDIA6_MOUSE | Protein disulfide-isomerase A6 OS=Mus musculus GN=Pdia6 PE=1 SV=3 | MOUSE | 9 |
| sp\|P06728\|APOA4_MOUSE | Apolipoprotein A-IV OS=Mus musculus GN=Apoa4 PE=1 SV=3 | MOUSE | 9 |
| sp\|Q8BG05\|ROA3_MOUSE | Heterogeneous nuclear ribonucleoprotein A3 OS=Mus musculus GN=Hnrnpa3 PE=1 SV=1 | MOUSE | 9 |
| sp\|Q60930\|VDAC2_MOUSE | Voltage-dependent anion-selective channel protein 2 OS=Mus musculus GN=Vdac2 PE=1 SV= | MOUSE | 9 |
| sp\|P16858\|G3P_MOUSE | Glyceraldehyde-3-phosphate dehydrogenase OS=Mus musculus GN=Gapdh PE=1 SV=2 | MOUSE | 9 |
| sp\|Q3THW5\|H2AV_MOUSE | Histone H2A.V OS=Mus musculus GN=H2afv PE=1 SV=3 | MOUSE | 9 |
| sp\|P0C0S6\|H2AZ_MOUSE | Histone H2A.Z OS=Mus musculus GN=H2afz PE=1 SV=2 | MOUSE | 9 |
| sp\|Q06318\|UTER_MOUSE | Uteroglobin OS=Mus musculus GN=Scgb1a1 PE=1 SV=1 | MOUSE | 9 |
| sp\|Q61592\|GAS6_MOUSE | Growth arrest-specific protein 6 OS=Mus musculus GN=Gas6 PE=2 SV=2 | MOUSE | 8 |
| sp\|Q9CQC6\|BZW1_MOUSE | Basic leucine zipper and W2 domain-containing protein 1 OS=Mus musculus GN=Bzw1 PE=1 | MOUSE | 8 |
| sp\|Q08761\|PROS_MOUSE | Vitamin K-dependent protein S OS=Mus musculus GN=Pros1 PE=2 SV=1 | MOUSE | 8 |
| sp\|Q9Z1N5\|DX39B_MOUSE | Spliceosome RNA helicase Ddx39b OS=Mus musculus GN=Ddx39b PE=1 SV=1 | MOUSE | 8 |
| sp\|P20918\|PLMN_MOUSE | Plasminogen OS=Mus musculus GN=Plg PE=1 SV=3 | MOUSE | 8 |
| sp\|Q6ZWX6\|IF2A_MOUSE | Eukaryotic translation initiation factor 2 subunit 1 OS=Mus musculus GN=Eif2s1 PE=1 SV=3 | MOUSE | 8 |
| sp\|Q9WU78\|PDC6I_MOUSE | Programmed cell death 6-interacting protein OS=Mus musculus GN=Pdcd6ip PE=1 SV=3 | MOUSE | 8 |
| sp\|P63101\|1433Z_MOUSE | 14-3-3 protein zeta/delta OS=Mus musculus GN=Ywhaz PE=1 SV=1 | MOUSE | 8 |
| sp\|P18760\|COF1_MOUSE | Cofilin-1 OS=Mus musculus GN=Cfl1 PE=1 SV=3 | MOUSE | 8 |
| sp\|P10126\|EF1A1_MOUSE | Elongation factor 1-alpha 1 OS=Mus musculus GN=Eef1a1 PE=1 SV=3 | MOUSE | 8 |
| sp\|O70370\|CATS_MOUSE | Cathepsin S OS=Mus musculus GN=Ctss PE=1 SV=2 | MOUSE | 8 |
| sp\|O08992\|SDCB1_MOUSE | Syntenin-1 OS=Mus musculus GN=Sdcbp PE=1 SV=1 | MOUSE | 8 |
| sp\|Q6IRU2\|TPM4_MOUSE | Tropomyosin alpha-4 chain OS=Mus musculus GN=Tpm4 PE=1 SV=3 | MOUSE | 8 |
| sp\|Q9CQI6\|COTL1_MOUSE | Coactosin-like protein OS=Mus musculus GN=Cotl1 PE=1 SV=3 | MOUSE | 8 |
| sp\|P62204\|CALM_MOUSE | Calmodulin OS=Mus musculus GN=Calm1 PE=1 SV=2 | MOUSE | 8 |
| sp\|P84244\|H33_MOUSE | Histone H3.3 OS=Mus musculus GN=H3f3a PE=1 SV=2 | MOUSE | 8 |
| sp\|P02301\|H3C_MOUSE | Histone H3.3C OS=Mus musculus GN=H3f3c PE=3 SV=3 | MOUSE | 8 |
| sp\|P02089\|HBB2_MOUSE | Hemoglobin subunit beta-2 OS=Mus musculus GN=Hbb-b2 PE=1 SV=2 | MOUSE | 8 |
| sp\|P14733\|LMNB1_MOUSE | Lamin-B1 OS=Mus musculus GN=Lmnb1 PE=1 SV=3 | MOUSE | 7 |
| sp\|Q62422\|OSTF1_MOUSE | Osteoclast-stimulating factor 1 OS=Mus musculus GN=Ostf1 PE=1 SV=2 | MOUSE | 7 |
| sp\|P60843\|IF4A1_MOUSE | Eukaryotic initiation factor 4A-I OS=Mus musculus GN=Eif4a1 PE=1 SV=1 | MOUSE | 7 |
| sp\|P51881\|ADT2_MOUSE | ADP/ATP translocase 2 OS=Mus musculus GN=Slc25a5 PE=1 SV=3 | MOUSE | 7 |
| sp\|P21107\|TPM3_MOUSE | Tropomyosin alpha-3 chain OS=Mus musculus GN=Tpm3 PE=1 SV=3 | MOUSE | 7 |
| sp\|P24369\|PPIB_MOUSE | Peptidyl-prolyl cis-trans isomerase B OS=Mus musculus GN=Ppib PE=1 SV=2 | MOUSE | 7 |
| sp\|O35381\|AN32A_MOUSE | Acidic leucine-rich nuclear phosphoprotein 32 family member A OS=Mus musculus GN=Anp32 | MOUSE | 7 |

| | | | |
|---|---|---|---|
| sp\|P62984\|RL40_MOUSE | Ubiquitin-60S ribosomal protein L40 OS=Mus musculus GN=Uba52 PE=1 SV=2 | MOUSE | 7 |
| sp\|P62983\|RS27A_MOUSE | Ubiquitin-40S ribosomal protein S27a OS=Mus musculus GN=Rps27a PE=1 SV=2 | MOUSE | 7 |
| sp\|P0CG49\|UBB_MOUSE | Polyubiquitin-B OS=Mus musculus GN=Ubb PE=2 SV=1 | MOUSE | 7 |
| sp\|P0CG50\|UBC_MOUSE | Polyubiquitin-C OS=Mus musculus GN=Ubc PE=1 SV=2 | MOUSE | 7 |
| sp\|P01942\|HBA_MOUSE | Hemoglobin subunit alpha OS=Mus musculus GN=Hba PE=1 SV=2 | MOUSE | 7 |
| sp\|P43277\|H13_MOUSE | Histone H1.3 OS=Mus musculus GN=Hist1h1d PE=1 SV=2 | MOUSE | 7 |
| sp\|P16110\|LEG3_MOUSE | Galectin-3 OS=Mus musculus GN=Lgals3 PE=1 SV=3 | MOUSE | 7 |
| sp\|P23953\|EST1C_MOUSE | Carboxylesterase 1C OS=Mus musculus GN=Ces1c PE=1 SV=4 | MOUSE | 7 |
| sp\|Q6ZQ38\|CAND1_MOUSE | Cullin-associated NEDD8-dissociated protein 1 OS=Mus musculus GN=Cand1 PE=1 SV=2 | MOUSE | 7 |
| sp\|Q3THE2\|ML12B_MOUSE | Myosin regulatory light chain 12B OS=Mus musculus GN=Myl12b PE=1 SV=2 | MOUSE | 7 |
| sp\|O88947\|FA10_MOUSE | Coagulation factor X OS=Mus musculus GN=F10 PE=1 SV=1 | MOUSE | 7 |
| sp\|Q6URW6\|MYH14_MOUSE | Myosin-14 OS=Mus musculus GN=Myh14 PE=1 SV=1 | MOUSE | 7 |
| sp\|Q9CQV8\|1433B_MOUSE | 14-3-3 protein beta/alpha OS=Mus musculus GN=Ywhab PE=1 SV=3 | MOUSE | 7 |
| sp\|P68254\|1433T_MOUSE | 14-3-3 protein theta OS=Mus musculus GN=Ywhaq PE=1 SV=1 | MOUSE | 7 |
| sp\|P62259\|1433E_MOUSE | 14-3-3 protein epsilon OS=Mus musculus GN=Ywhae PE=1 SV=1 | MOUSE | 7 |
| sp\|P43274\|H14_MOUSE | Histone H1.4 OS=Mus musculus GN=Hist1h1e PE=1 SV=2 | MOUSE | 7 |
| sp\|P14869\|RLA0_MOUSE | 60S acidic ribosomal protein P0 OS=Mus musculus GN=Rplp0 PE=1 SV=3 | MOUSE | 6 |
| sp\|Q69ZN7\|MYOF_MOUSE | Myoferlin OS=Mus musculus GN=Myof PE=1 SV=2 | MOUSE | 6 |
| sp\|Q60605\|MYL6_MOUSE | Myosin light polypeptide 6 OS=Mus musculus GN=Myl6 PE=1 SV=3 | MOUSE | 6 |
| sp\|Q8VEK3\|HNRPU_MOUSE | Heterogeneous nuclear ribonucleoprotein U OS=Mus musculus GN=Hnrnpu PE=1 SV=1 | MOUSE | 6 |
| sp\|Q60932\|VDAC1_MOUSE | Voltage-dependent anion-selective channel protein 1 OS=Mus musculus GN=Vdac1 PE=1 SV= | MOUSE | 6 |
| sp\|Q03265\|ATPA_MOUSE | ATP synthase subunit alpha, mitochondrial OS=Mus musculus GN=Atp5a1 PE=1 SV=1 | MOUSE | 6 |
| sp\|Q9WUU7\|CATZ_MOUSE | Cathepsin Z OS=Mus musculus GN=Ctsz PE=1 SV=1 | MOUSE | 6 |
| sp\|P49290\|PERE_MOUSE | Eosinophil peroxidase OS=Mus musculus GN=Epx PE=1 SV=2 | MOUSE | 6 |
| sp\|Q61029\|LAP2B_MOUSE | Lamina-associated polypeptide 2, isoforms beta/delta/epsilon/gamma OS=Mus musculus GN=T | MOUSE | 6 |
| sp\|Q61033\|LAP2A_MOUSE | Lamina-associated polypeptide 2, isoforms alpha/zeta OS=Mus musculus GN=Tmpo PE=1 SV | MOUSE | 6 |
| sp\|Q9EQU5\|SET_MOUSE | Protein SET OS=Mus musculus GN=Set PE=1 SV=1 | MOUSE | 6 |
| sp\|Q99L47\|F10A1_MOUSE | Hsc70-interacting protein OS=Mus musculus GN=St13 PE=1 SV=1 | MOUSE | 6 |
| sp\|P15864\|H12_MOUSE | Histone H1.2 OS=Mus musculus GN=Hist1h1c PE=1 SV=2 | MOUSE | 6 |
| sp\|P21614\|VTDB_MOUSE | Vitamin D-binding protein OS=Mus musculus GN=Gc PE=1 SV=2 | MOUSE | 6 |
| sp\|P29416\|HEXA_MOUSE | Beta-hexosaminidase subunit alpha OS=Mus musculus GN=Hexa PE=1 SV=2 | MOUSE | 6 |
| sp\|P61979\|HNRPK_MOUSE | Heterogeneous nuclear ribonucleoprotein K OS=Mus musculus GN=Hnrnpk PE=1 SV=1 | MOUSE | 6 |
| sp\|P29391\|FRIL1_MOUSE | Ferritin light chain 1 OS=Mus musculus GN=Ftl1 PE=1 SV=2 | MOUSE | 6 |
| sp\|P63158\|HMGB1_MOUSE | High mobility group protein B1 OS=Mus musculus GN=Hmgb1 PE=1 SV=2 | MOUSE | 6 |
| sp\|Q99PT1\|GDIR1_MOUSE | Rho GDP-dissociation inhibitor 1 OS=Mus musculus GN=Arhgdia PE=1 SV=3 | MOUSE | 6 |
| sp\|Q91WP6\|SPA3N_MOUSE | Serine protease inhibitor A3N OS=Mus musculus GN=Serpina3n PE=1 SV=1 | MOUSE | 6 |
| sp\|P28798\|GRN_MOUSE | Granulins OS=Mus musculus GN=Grn PE=1 SV=2 | MOUSE | 6 |
| sp\|Q9QZQ8\|H2AY_MOUSE | Core histone macro-H2A.1 OS=Mus musculus GN=H2afy PE=1 SV=3 | MOUSE | 6 |
| sp\|P01887\|B2MG_MOUSE | Beta-2-microglobulin OS=Mus musculus GN=B2m PE=1 SV=2 | MOUSE | 6 |
| sp\|P68510\|1433F_MOUSE | 14-3-3 protein eta OS=Mus musculus GN=Ywhah PE=1 SV=2 | MOUSE | 6 |
| sp\|P27005\|S10A8_MOUSE | Protein S100-A8 OS=Mus musculus GN=S100a8 PE=1 SV=3 | MOUSE | 6 |
| sp\|Q91YQ5\|RPN1_MOUSE | Dolichyl-diphosphooligosaccharide--protein glycosyltransferase subunit 1 OS=Mus musculus G | MOUSE | 5 |
| sp\|Q61598\|GDIB_MOUSE | Rab GDP dissociation inhibitor beta OS=Mus musculus GN=Gdi2 PE=1 SV=1 | MOUSE | 5 |
| sp\|P70663\|SPRL1_MOUSE | SPARC-like protein 1 OS=Mus musculus GN=Sparcl1 PE=1 SV=3 | MOUSE | 5 |
| sp\|P62137\|PP1A_MOUSE | Serine/threonine-protein phosphatase PP1-alpha catalytic subunit OS=Mus musculus GN=Ppp | MOUSE | 5 |
| sp\|O08692\|NGP_MOUSE | Neutrophilic granule protein OS=Mus musculus GN=Ngp PE=1 SV=1 | MOUSE | 5 |
| sp\|Q9QXH4\|ITAX_MOUSE | Integrin alpha-X OS=Mus musculus GN=Itgax PE=1 SV=1 | MOUSE | 5 |
| sp\|Q3TEA8\|HP1B3_MOUSE | Heterochromatin protein 1-binding protein 3 OS=Mus musculus GN=Hp1bp3 PE=1 SV=1 | MOUSE | 5 |
| sp\|P56480\|ATPB_MOUSE | ATP synthase subunit beta, mitochondrial OS=Mus musculus GN=Atp5b PE=1 SV=2 | MOUSE | 5 |
| sp\|P19973\|LSP1_MOUSE | Lymphocyte-specific protein 1 OS=Mus musculus GN=Lsp1 PE=1 SV=2 | MOUSE | 5 |
| sp\|Q9Z0J0\|NPC2_MOUSE | Epididymal secretory protein E1 OS=Mus musculus GN=Npc2 PE=1 SV=1 | MOUSE | 5 |
| sp\|P12023\|A4_MOUSE | Amyloid beta A4 protein OS=Mus musculus GN=App PE=1 SV=3 | MOUSE | 5 |
| sp\|Q68FL4\|SAHH3_MOUSE | Putative adenosylhomocysteinase 3 OS=Mus musculus GN=Ahcyl2 PE=1 SV=1 | MOUSE | 5 |
| sp\|Q9JKF1\|IQGA1_MOUSE | Ras GTPase-activating-like protein IQGAP1 OS=Mus musculus GN=Iqgap1 PE=1 SV=2 | MOUSE | 5 |
| sp\|P01867\|IGG2B_MOUSE | Ig gamma-2B chain C region OS=Mus musculus GN=Igh-3 PE=1 SV=3 | MOUSE | 5 |
| sp\|O35685\|NUDC_MOUSE | Nuclear migration protein nudC OS=Mus musculus GN=Nudc PE=1 SV=1 | MOUSE | 5 |
| sp\|P24452\|CAPG_MOUSE | Macrophage-capping protein OS=Mus musculus GN=Capg PE=1 SV=2 | MOUSE | 5 |
| sp\|Q9WV54\|ASAH1_MOUSE | Acid ceramidase OS=Mus musculus GN=Asah1 PE=1 SV=1 | MOUSE | 5 |
| sp\|Q8BL97\|SRSF7_MOUSE | Serine/arginine-rich splicing factor 7 OS=Mus musculus GN=Srsf7 PE=1 SV=1 | MOUSE | 5 |
| sp\|P06797\|CATL1_MOUSE | Cathepsin L1 OS=Mus musculus GN=Ctsl PE=1 SV=2 | MOUSE | 5 |
| sp\|P14847\|CRP_MOUSE | C-reactive protein OS=Mus musculus GN=Crp PE=1 SV=2 | MOUSE | 5 |
| sp\|Q3UM45\|PP1R7_MOUSE | Protein phosphatase 1 regulatory subunit 7 OS=Mus musculus GN=Ppp1r7 PE=1 SV=2 | MOUSE | 5 |
| sp\|O88569\|ROA2_MOUSE | Heterogeneous nuclear ribonucleoproteins A2/B1 OS=Mus musculus GN=Hnrnpa2b1 PE=1 SV | MOUSE | 5 |
| sp\|Q9EST5\|AN32B_MOUSE | Acidic leucine-rich nuclear phosphoprotein 32 family member B OS=Mus musculus GN=Anp32 | MOUSE | 5 |
| sp\|P61982\|1433G_MOUSE | 14-3-3 protein gamma OS=Mus musculus GN=Ywhag PE=1 SV=2 | MOUSE | 5 |
| sp\|P06151\|LDHA_MOUSE | L-lactate dehydrogenase A chain OS=Mus musculus GN=Ldha PE=1 SV=3 | MOUSE | 4 |

| | | | |
|---|---|---|---|
| sp\|Q9Z0N1\|IF2G_MOUSE | Eukaryotic translation initiation factor 2 subunit 3, X-linked OS=Mus musculus GN=Eif2s3x PE= | MOUSE | 4 |
| sp\|Q9ESB3\|HRG_MOUSE | Histidine-rich glycoprotein OS=Mus musculus GN=Hrg PE=1 SV=2 | MOUSE | 4 |
| sp\|Q80SW1\|SAHH2_MOUSE | Putative adenosylhomocysteinase 2 OS=Mus musculus GN=Ahcyl1 PE=1 SV=1 | MOUSE | 4 |
| sp\|P60766\|CDC42_MOUSE | Cell division control protein 42 homolog OS=Mus musculus GN=Cdc42 PE=1 SV=2 | MOUSE | 4 |
| sp\|Q61646\|HPT_MOUSE | Haptoglobin OS=Mus musculus GN=Hp PE=1 SV=1 | MOUSE | 4 |
| sp\|Q9CQI3\|GMFB_MOUSE | Glia maturation factor beta OS=Mus musculus GN=Gmfb PE=1 SV=3 | MOUSE | 4 |
| sp\|P49945\|FRIL2_MOUSE | Ferritin light chain 2 OS=Mus musculus GN=Ftl2 PE=3 SV=2 | MOUSE | 4 |
| sp\|Q9WUM4\|COR1C_MOUSE | Coronin-1C OS=Mus musculus GN=Coro1c PE=1 SV=2 | MOUSE | 4 |
| sp\|P16675\|PPGB_MOUSE | Lysosomal protective protein OS=Mus musculus GN=Ctsa PE=1 SV=1 | MOUSE | 4 |
| sp\|Q9JHK5\|PLEK_MOUSE | Pleckstrin OS=Mus musculus GN=Plek PE=1 SV=1 | MOUSE | 4 |
| sp\|Q8BHG1\|NRDC_MOUSE | Nardilysin OS=Mus musculus GN=Nrd1 PE=1 SV=1 | MOUSE | 4 |
| sp\|O70251\|EF1B_MOUSE | Elongation factor 1-beta OS=Mus musculus GN=Eef1b PE=1 SV=5 | MOUSE | 4 |
| sp\|P17182\|ENOA_MOUSE | Alpha-enolase OS=Mus musculus GN=Eno1 PE=1 SV=3 | MOUSE | 4 |
| sp\|P08249\|MDHM_MOUSE | Malate dehydrogenase, mitochondrial OS=Mus musculus GN=Mdh2 PE=1 SV=3 | MOUSE | 4 |
| sp\|Q9D1G1\|RAB1B_MOUSE | Ras-related protein Rab-1B OS=Mus musculus GN=Rab1b PE=1 SV=1 | MOUSE | 4 |
| sp\|P62821\|RAB1A_MOUSE | Ras-related protein Rab-1A OS=Mus musculus GN=Rab1A PE=1 SV=3 | MOUSE | 4 |
| sp\|Q922Q8\|LRC59_MOUSE | Leucine-rich repeat-containing protein 59 OS=Mus musculus GN=Lrrc59 PE=1 SV=1 | MOUSE | 4 |
| sp\|P51150\|RAB7A_MOUSE | Ras-related protein Rab-7a OS=Mus musculus GN=Rab7a PE=1 SV=2 | MOUSE | 4 |
| sp\|P62827\|RAN_MOUSE | GTP-binding nuclear protein Ran OS=Mus musculus GN=Ran PE=1 SV=3 | MOUSE | 4 |
| sp\|P62962\|PROF1_MOUSE | Profilin-1 OS=Mus musculus GN=Pfn1 PE=1 SV=2 | MOUSE | 4 |
| sp\|P49182\|HEP2_MOUSE | Heparin cofactor 2 OS=Mus musculus GN=Serpind1 PE=1 SV=1 | MOUSE | 4 |
| sp\|O89017\|LGMN_MOUSE | Legumain OS=Mus musculus GN=Lgmn PE=1 SV=1 | MOUSE | 4 |
| sp\|Q9D8B3\|CHM4B_MOUSE | Charged multivesicular body protein 4b OS=Mus musculus GN=Chmp4b PE=1 SV=2 | MOUSE | 4 |
| sp\|Q8K1I3\|SPP24_MOUSE | Secreted phosphoprotein 24 OS=Mus musculus GN=Spp2 PE=1 SV=2 | MOUSE | 4 |
| sp\|Q9Z204\|HNRPC_MOUSE | Heterogeneous nuclear ribonucleoproteins C1/C2 OS=Mus musculus GN=Hnrnpc PE=1 SV=1 | MOUSE | 4 |
| sp\|P01869\|IGH1M_MOUSE | Ig gamma-1 chain C region, membrane-bound form OS=Mus musculus GN=Ighg1 PE=1 SV=2 | MOUSE | 4 |
| sp\|P01868\|IGHG1_MOUSE | Ig gamma-1 chain C region secreted form OS=Mus musculus GN=Ighg1 PE=1 SV=1 | MOUSE | 4 |
| sp\|P97384\|ANX11_MOUSE | Annexin A11 OS=Mus musculus GN=Anxa11 PE=1 SV=2 | MOUSE | 4 |
| sp\|O70194\|EIF3D_MOUSE | Eukaryotic translation initiation factor 3 subunit D OS=Mus musculus GN=Eif3d PE=1 SV=2 | MOUSE | 4 |
| sp\|Q01768\|NDKB_MOUSE | Nucleoside diphosphate kinase B OS=Mus musculus GN=Nme2 PE=1 SV=1 | MOUSE | 4 |
| sp\|P63001\|RAC1_MOUSE | Ras-related C3 botulinum toxin substrate 1 OS=Mus musculus GN=Rac1 PE=1 SV=1 | MOUSE | 4 |
| sp\|P60764\|RAC3_MOUSE | Ras-related C3 botulinum toxin substrate 3 OS=Mus musculus GN=Rac3 PE=1 SV=1 | MOUSE | 4 |
| sp\|Q9Z1Q5\|CLIC1_MOUSE | Chloride intracellular channel protein 1 OS=Mus musculus GN=Clic1 PE=1 SV=3 | MOUSE | 4 |
| sp\|P62897\|CYC_MOUSE | Cytochrome c, somatic OS=Mus musculus GN=Cycs PE=1 SV=2 | MOUSE | 4 |
| sp\|P33587\|PROC_MOUSE | Vitamin K-dependent protein C OS=Mus musculus GN=Proc PE=1 SV=2 | MOUSE | 4 |
| sp\|O08784\|TCOF_MOUSE | Treacle protein OS=Mus musculus GN=Tcof1 PE=1 SV=1 | MOUSE | 4 |
| sp\|Q60817\|NACA_MOUSE | Nascent polypeptide-associated complex subunit alpha OS=Mus musculus GN=Naca PE=1 S\ | MOUSE | 4 |
| sp\|P70670\|NACAM_MOUSE | Nascent polypeptide-associated complex subunit alpha, muscle-specific form OS=Mus muscul | MOUSE | 4 |
| sp\|B9EJ86\|OSBL8_MOUSE | Oxysterol-binding protein-related protein 8 OS=Mus musculus GN=Osbpl8 PE=1 SV=1 | MOUSE | 4 |
| sp\|P84104\|SRSF3_MOUSE | Serine/arginine-rich splicing factor 3 OS=Mus musculus GN=Srsf3 PE=1 SV=1 | MOUSE | 4 |
| sp\|P51859\|HDGF_MOUSE | Hepatoma-derived growth factor OS=Mus musculus GN=Hdgf PE=1 SV=2 | MOUSE | 4 |
| sp\|P02463\|CO4A1_MOUSE | Collagen alpha-1(IV) chain OS=Mus musculus GN=Col4a1 PE=1 SV=4 | MOUSE | 4 |
| sp\|P01865\|GCAM_MOUSE | Ig gamma-2A chain C region, membrane-bound form OS=Mus musculus GN=Igh-1a PE=1 SV= | MOUSE | 4 |
| sp\|P01863\|GCAA_MOUSE | Ig gamma-2A chain C region, A allele OS=Mus musculus GN=Ighg PE=1 SV=1 | MOUSE | 4 |
| sp\|P40142\|TKT_MOUSE | Transketolase OS=Mus musculus GN=Tkt PE=1 SV=1 | MOUSE | 4 |
| sp\|P55097\|CATK_MOUSE | Cathepsin K OS=Mus musculus GN=Ctsk PE=2 SV=2 | MOUSE | 4 |
| sp\|P34928\|APOC1_MOUSE | Apolipoprotein C-I OS=Mus musculus GN=Apoc1 PE=1 SV=1 | MOUSE | 4 |
| sp\|P62141\|PP1B_MOUSE | Serine/threonine-protein phosphatase PP1-beta catalytic subunit OS=Mus musculus GN=Ppp1 | MOUSE | 4 |
| sp\|P01837\|IGKC_MOUSE | Ig kappa chain C region OS=Mus musculus PE=1 SV=1 | MOUSE | 4 |
| sp\|P10639\|THIO_MOUSE | Thioredoxin OS=Mus musculus GN=Txn PE=1 SV=3 | MOUSE | 4 |
| sp\|P04939\|MUP3_MOUSE | Major urinary protein 3 OS=Mus musculus GN=Mup3 PE=1 SV=1 | MOUSE | 4 |
| sp\|Q9D8N0\|EF1G_MOUSE | Elongation factor 1-gamma OS=Mus musculus GN=Eef1g PE=1 SV=3 | MOUSE | 3 |
| sp\|P50580\|PA2G4_MOUSE | Proliferation-associated protein 2G4 OS=Mus musculus GN=Pa2g4 PE=1 SV=3 | MOUSE | 3 |
| sp\|Q05793\|PGBM_MOUSE | Basement membrane-specific heparan sulfate proteoglycan core protein OS=Mus musculus GI | MOUSE | 3 |
| sp\|O35405\|PLD3_MOUSE | Phospholipase D3 OS=Mus musculus GN=Pld3 PE=1 SV=1 | MOUSE | 3 |
| sp\|Q06890\|CLUS_MOUSE | Clusterin OS=Mus musculus GN=Clu PE=1 SV=1 | MOUSE | 3 |
| sp\|Q8CCK0\|H2AW_MOUSE | Core histone macro-H2A.2 OS=Mus musculus GN=H2afy2 PE=1 SV=3 | MOUSE | 3 |
| sp\|Q05144\|RAC2_MOUSE | Ras-related C3 botulinum toxin substrate 2 OS=Mus musculus GN=Rac2 PE=1 SV=1 | MOUSE | 3 |
| sp\|Q05117\|PPA5_MOUSE | Tartrate-resistant acid phosphatase type 5 OS=Mus musculus GN=Acp5 PE=1 SV=2 | MOUSE | 3 |
| sp\|A1L314\|MPEG1_MOUSE | Macrophage-expressed gene 1 protein OS=Mus musculus GN=Mpeg1 PE=1 SV=1 | MOUSE | 3 |
| sp\|P68037\|UB2L3_MOUSE | Ubiquitin-conjugating enzyme E2 L3 OS=Mus musculus GN=Ube2l3 PE=1 SV=1 | MOUSE | 3 |
| sp\|P57776\|EF1D_MOUSE | Elongation factor 1-delta OS=Mus musculus GN=Eef1d PE=1 SV=3 | MOUSE | 3 |
| sp\|P11031\|TCP4_MOUSE | Activated RNA polymerase II transcriptional coactivator p15 OS=Mus musculus GN=Sub1 PE= | MOUSE | 3 |
| sp\|P10493\|NID1_MOUSE | Nidogen-1 OS=Mus musculus GN=Nid1 PE=1 SV=2 | MOUSE | 3 |
| sp\|Q3UHX2\|HAP28_MOUSE | 28 kDa heat- and acid-stable phosphoprotein OS=Mus musculus GN=Pdap1 PE=1 SV=1 | MOUSE | 3 |
| sp\|Q9DCN2\|NB5R3_MOUSE | NADH-cytochrome b5 reductase 3 OS=Mus musculus GN=Cyb5r3 PE=1 SV=3 | MOUSE | 3 |

| | | | |
|---|---|---|---|
| sp\|P59325\|IF5_MOUSE | Eukaryotic translation initiation factor 5 OS=Mus musculus GN=Eif5 PE=1 SV=1 | MOUSE | 3 |
| sp\|Q99JI6\|RAP1B_MOUSE | Ras-related protein Rap-1b OS=Mus musculus GN=Rap1b PE=1 SV=2 | MOUSE | 3 |
| sp\|P62835\|RAP1A_MOUSE | Ras-related protein Rap-1A OS=Mus musculus GN=Rap1a PE=1 SV=1 | MOUSE | 3 |
| sp\|P28656\|NP1L1_MOUSE | Nucleosome assembly protein 1-like 1 OS=Mus musculus GN=Nap1l1 PE=1 SV=2 | MOUSE | 3 |
| sp\|P01899\|HA11_MOUSE | H-2 class I histocompatibility antigen, D-B alpha chain OS=Mus musculus GN=H2-D1 PE=1 S\ | MOUSE | 3 |
| sp\|P01897\|HA1L_MOUSE | H-2 class I histocompatibility antigen, L-D alpha chain OS=Mus musculus GN=H2-L PE=1 SV= | MOUSE | 3 |
| sp\|Q920B9\|SP16H_MOUSE | FACT complex subunit SPT16 OS=Mus musculus GN=Supt16h PE=1 SV=2 | MOUSE | 3 |
| sp\|P05202\|AATM_MOUSE | Aspartate aminotransferase, mitochondrial OS=Mus musculus GN=Got2 PE=1 SV=1 | MOUSE | 3 |
| sp\|Q60648\|SAP3_MOUSE | Ganglioside GM2 activator OS=Mus musculus GN=Gm2a PE=1 SV=2 | MOUSE | 3 |
| sp\|Q8BH64\|EHD2_MOUSE | EH domain-containing protein 2 OS=Mus musculus GN=Ehd2 PE=1 SV=1 | MOUSE | 3 |
| sp\|Q9CPW4\|ARPC5_MOUSE | Actin-related protein 2/3 complex subunit 5 OS=Mus musculus GN=Arpc5 PE=1 SV=3 | MOUSE | 3 |
| sp\|Q9QUI0\|RHOA_MOUSE | Transforming protein RhoA OS=Mus musculus GN=Rhoa PE=1 SV=1 | MOUSE | 3 |
| sp\|Q62159\|RHOC_MOUSE | Rho-related GTP-binding protein RhoC OS=Mus musculus GN=Rhoc PE=1 SV=2 | MOUSE | 3 |
| sp\|Q8VCM7\|FIBG_MOUSE | Fibrinogen gamma chain OS=Mus musculus GN=Fgg PE=1 SV=1 | MOUSE | 3 |
| sp\|P67984\|RL22_MOUSE | 60S ribosomal protein L22 OS=Mus musculus GN=Rpl22 PE=1 SV=2 | MOUSE | 3 |
| sp\|P23116\|EIF3A_MOUSE | Eukaryotic translation initiation factor 3 subunit A OS=Mus musculus GN=Eif3a PE=1 SV=5 | MOUSE | 3 |
| sp\|P09813\|APOA2_MOUSE | Apolipoprotein A-II OS=Mus musculus GN=Apoa2 PE=1 SV=2 | MOUSE | 3 |
| sp\|Q9WVA4\|TAGL2_MOUSE | Transgelin-2 OS=Mus musculus GN=Tagln2 PE=1 SV=4 | MOUSE | 3 |
| sp\|Q08857\|CD36_MOUSE | Platelet glycoprotein 4 OS=Mus musculus GN=Cd36 PE=1 SV=2 | MOUSE | 3 |
| sp\|Q8BH35\|CO8B_MOUSE | Complement component C8 beta chain OS=Mus musculus GN=C8b PE=1 SV=1 | MOUSE | 3 |
| sp\|Q60931\|VDAC3_MOUSE | Voltage-dependent anion-selective channel protein 3 OS=Mus musculus GN=Vdac3 PE=1 SV= | MOUSE | 3 |
| sp\|A6X935\|ITIH4_MOUSE | Inter alpha-trypsin inhibitor, heavy chain 4 OS=Mus musculus GN=Itih4 PE=1 SV=2 | MOUSE | 3 |
| sp\|P63242\|IF5A1_MOUSE | Eukaryotic translation initiation factor 5A-1 OS=Mus musculus GN=Eif5a PE=1 SV=2 | MOUSE | 3 |
| sp\|Q8BGY2\|IF5A2_MOUSE | Eukaryotic translation initiation factor 5A-2 OS=Mus musculus GN=Eif5a2 PE=1 SV=3 | MOUSE | 3 |
| sp\|Q9D1A2\|CNDP2_MOUSE | Cytosolic non-specific dipeptidase OS=Mus musculus GN=Cndp2 PE=1 SV=1 | MOUSE | 3 |
| sp\|P97298\|PEDF_MOUSE | Pigment epithelium-derived factor OS=Mus musculus GN=Serpinf1 PE=1 SV=2 | MOUSE | 3 |
| sp\|P06800\|PTPRC_MOUSE | Receptor-type tyrosine-protein phosphatase C OS=Mus musculus GN=Ptprc PE=1 SV=3 | MOUSE | 3 |
| sp\|P09528\|FRIH_MOUSE | Ferritin heavy chain OS=Mus musculus GN=Fth1 PE=1 SV=2 | MOUSE | 3 |
| sp\|P62702\|RS4X_MOUSE | 40S ribosomal protein S4, X isoform OS=Mus musculus GN=Rps4x PE=1 SV=2 | MOUSE | 3 |
| sp\|Q8BTM8\|FLNA_MOUSE | Filamin-A OS=Mus musculus GN=Flna PE=1 SV=5 | MOUSE | 3 |
| sp\|P62320\|SMD3_MOUSE | Small nuclear ribonucleoprotein Sm D3 OS=Mus musculus GN=Snrpd3 PE=1 SV=1 | MOUSE | 3 |
| sp\|P61027\|RAB10_MOUSE | Ras-related protein Rab-10 OS=Mus musculus GN=Rab10 PE=1 SV=1 | MOUSE | 3 |
| sp\|O88456\|CPNS1_MOUSE | Calpain small subunit 1 OS=Mus musculus GN=Capns1 PE=1 SV=1 | MOUSE | 3 |
| sp\|Q9CXW3\|CYBP_MOUSE | Calcyclin-binding protein OS=Mus musculus GN=Cacybp PE=1 SV=1 | MOUSE | 3 |
| sp\|O35841\|API5_MOUSE | Apoptosis inhibitor 5 OS=Mus musculus GN=Api5 PE=1 SV=2 | MOUSE | 3 |
| sp\|P08122\|CO4A2_MOUSE | Collagen alpha-2(IV) chain OS=Mus musculus GN=Col4a2 PE=1 SV=4 | MOUSE | 3 |
| sp\|P61028\|RAB8B_MOUSE | Ras-related protein Rab-8B OS=Mus musculus GN=Rab8b PE=1 SV=1 | MOUSE | 3 |
| sp\|P01864\|GCAB_MOUSE | Ig gamma-2A chain C region secreted form OS=Mus musculus PE=1 SV=1 | MOUSE | 3 |
| sp\|Q9QZS0\|CO4A3_MOUSE | Collagen alpha-3(IV) chain OS=Mus musculus GN=Col4a3 PE=1 SV=2 | MOUSE | 3 |
| sp\|P48962\|ADT1_MOUSE | ADP/ATP translocase 1 OS=Mus musculus GN=Slc25a4 PE=1 SV=4 | MOUSE | 3 |
| sp\|P15532\|NDKA_MOUSE | Nucleoside diphosphate kinase A OS=Mus musculus GN=Nme1 PE=1 SV=1 | MOUSE | 3 |
| sp\|P46467\|VPS4B_MOUSE | Vacuolar protein sorting-associated protein 4B OS=Mus musculus GN=Vps4b PE=1 SV=2 | MOUSE | 2 |
| sp\|P01896\|HA1Z_MOUSE | H-2 class I histocompatibility antigen, alpha chain (Fragment) OS=Mus musculus PE=2 SV=1 | MOUSE | 2 |
| sp\|P01898\|HA10_MOUSE | H-2 class I histocompatibility antigen, Q10 alpha chain OS=Mus musculus GN=H2-Q10 PE=1 S | MOUSE | 2 |
| sp\|P14426\|HA13_MOUSE | H-2 class I histocompatibility antigen, D-K alpha chain OS=Mus musculus GN=H2-D1 PE=1 S\ | MOUSE | 2 |
| sp\|Q61147\|CERU_MOUSE | Ceruloplasmin OS=Mus musculus GN=Cp PE=1 SV=2 | MOUSE | 2 |
| sp\|Q8R1B4\|EIF3C_MOUSE | Eukaryotic translation initiation factor 3 subunit C OS=Mus musculus GN=Eif3c PE=1 SV=1 | MOUSE | 2 |
| sp\|P23198\|CBX3_MOUSE | Chromobox protein homolog 3 OS=Mus musculus GN=Cbx3 PE=1 SV=2 | MOUSE | 2 |
| sp\|P60867\|RS20_MOUSE | 40S ribosomal protein S20 OS=Mus musculus GN=Rps20 PE=1 SV=1 | MOUSE | 2 |
| sp\|Q60668\|HNRPD_MOUSE | Heterogeneous nuclear ribonucleoprotein D0 OS=Mus musculus GN=Hnrnpd PE=1 SV=2 | MOUSE | 2 |
| sp\|Q7TNV0\|DEK_MOUSE | Protein DEK OS=Mus musculus GN=Dek PE=1 SV=1 | MOUSE | 2 |
| sp\|Q8VDN2\|AT1A1_MOUSE | Sodium/potassium-transporting ATPase subunit alpha-1 OS=Mus musculus GN=Atp1a1 PE=1 | MOUSE | 2 |
| sp\|Q9R0P5\|DEST_MOUSE | Destrin OS=Mus musculus GN=Dstn PE=1 SV=3 | MOUSE | 2 |
| sp\|P08752\|GNAI2_MOUSE | Guanine nucleotide-binding protein G(i) subunit alpha-2 OS=Mus musculus GN=Gnai2 PE=1 S | MOUSE | 2 |
| sp\|Q99KN9\|EPN4_MOUSE | Clathrin interactor 1 OS=Mus musculus GN=Clint1 PE=1 SV=2 | MOUSE | 2 |
| sp\|Q61656\|DDX5_MOUSE | Probable ATP-dependent RNA helicase DDX5 OS=Mus musculus GN=Ddx5 PE=1 SV=2 | MOUSE | 2 |
| sp\|P97821\|CATC_MOUSE | Dipeptidyl peptidase 1 OS=Mus musculus GN=Ctsc PE=1 SV=1 | MOUSE | 2 |
| sp\|P31996\|CD68_MOUSE | Macrosialin OS=Mus musculus GN=Cd68 PE=1 SV=1 | MOUSE | 2 |
| sp\|Q9Z1T1\|AP3B1_MOUSE | AP-3 complex subunit beta-1 OS=Mus musculus GN=Ap3b1 PE=1 SV=2 | MOUSE | 2 |
| sp\|P21460\|CYTC_MOUSE | Cystatin-C OS=Mus musculus GN=Cst3 PE=1 SV=2 | MOUSE | 2 |
| sp\|Q8VCG4\|CO8G_MOUSE | Complement component C8 gamma chain OS=Mus musculus GN=C8g PE=1 SV=1 | MOUSE | 2 |
| sp\|Q8BMJ3\|IF1AX_MOUSE | Eukaryotic translation initiation factor 1A, X-chromosomal OS=Mus musculus GN=Eif1ax PE=2 | MOUSE | 2 |
| sp\|Q60872\|IF1A_MOUSE | Eukaryotic translation initiation factor 1A OS=Mus musculus GN=Eif1a PE=2 SV=3 | MOUSE | 2 |
| sp\|Q61878\|PRG2_MOUSE | Bone marrow proteoglycan OS=Mus musculus GN=Prg2 PE=1 SV=1 | MOUSE | 2 |
| sp\|P25444\|RS2_MOUSE | 40S ribosomal protein S2 OS=Mus musculus GN=Rps2 PE=1 SV=3 | MOUSE | 2 |
| sp\|P16294\|FA9_MOUSE | Coagulation factor IX OS=Mus musculus GN=F9 PE=2 SV=3 | MOUSE | 2 |

| | | | |
|---|---|---|---|
| sp\|P14483\|HB2A_MOUSE | H-2 class II histocompatibility antigen, A beta chain OS=Mus musculus GN=H2-Ab1 PE=1 SV= | MOUSE | 2 |
| sp\|P97371\|PSME1_MOUSE | Proteasome activator complex subunit 1 OS=Mus musculus GN=Psme1 PE=1 SV=2 | MOUSE | 2 |
| sp\|P35980\|RL18_MOUSE | 60S ribosomal protein L18 OS=Mus musculus GN=Rpl18 PE=1 SV=3 | MOUSE | 2 |
| sp\|Q99L45\|IF2B_MOUSE | Eukaryotic translation initiation factor 2 subunit 2 OS=Mus musculus GN=Eif2s2 PE=1 SV=1 | MOUSE | 2 |
| sp\|P28665\|MUG1_MOUSE | Murinoglobulin-1 OS=Mus musculus GN=Mug1 PE=1 SV=3 | MOUSE | 2 |
| sp\|P28666\|MUG2_MOUSE | Murinoglobulin-2 OS=Mus musculus GN=Mug2 PE=1 SV=2 | MOUSE | 2 |
| sp\|Q9CZ13\|QCR1_MOUSE | Cytochrome b-c1 complex subunit 1, mitochondrial OS=Mus musculus GN=Uqcrc1 PE=1 SV=2 | MOUSE | 2 |
| sp\|O08795\|GLU2B_MOUSE | Glucosidase 2 subunit beta OS=Mus musculus GN=Prkcsh PE=1 SV=1 | MOUSE | 2 |
| sp\|O54734\|OST48_MOUSE | Dolichyl-diphosphooligosaccharide--protein glycosyltransferase 48 kDa subunit OS=Mus musc | MOUSE | 2 |
| sp\|P28063\|PSB8_MOUSE | Proteasome subunit beta type-8 OS=Mus musculus GN=Psmb8 PE=1 SV=2 | MOUSE | 2 |
| sp\|Q8K182\|CO8A_MOUSE | Complement component C8 alpha chain OS=Mus musculus GN=C8a PE=1 SV=1 | MOUSE | 2 |
| sp\|P17742\|PPIA_MOUSE | Peptidyl-prolyl cis-trans isomerase A OS=Mus musculus GN=Ppia PE=1 SV=2 | MOUSE | 2 |
| sp\|Q9R0Q7\|TEBP_MOUSE | Prostaglandin E synthase 3 OS=Mus musculus GN=Ptges3 PE=1 SV=1 | MOUSE | 2 |
| sp\|P52480\|KPYM_MOUSE | Pyruvate kinase PKM OS=Mus musculus GN=Pkm PE=1 SV=4 | MOUSE | 2 |
| sp\|P53026\|RL10A_MOUSE | 60S ribosomal protein L10a OS=Mus musculus GN=Rpl10a PE=1 SV=3 | MOUSE | 2 |
| sp\|O35295\|PURB_MOUSE | Transcriptional activator protein Pur-beta OS=Mus musculus GN=Purb PE=1 SV=3 | MOUSE | 2 |
| sp\|O08677\|KNG1_MOUSE | Kininogen-1 OS=Mus musculus GN=Kng1 PE=1 SV=1 | MOUSE | 2 |
| sp\|P97822\|AN32E_MOUSE | Acidic leucine-rich nuclear phosphoprotein 32 family member E OS=Mus musculus GN=Anp32 | MOUSE | 2 |
| sp\|Q9Z0M5\|LICH_MOUSE | Lysosomal acid lipase/cholesteryl ester hydrolase OS=Mus musculus GN=Lipa PE=1 SV=2 | MOUSE | 2 |
| sp\|Q9CQL1\|MGN2_MOUSE | Protein mago nashi homolog 2 OS=Mus musculus GN=Magohb PE=2 SV=1 | MOUSE | 2 |
| sp\|P61327\|MGN_MOUSE | Protein mago nashi homolog OS=Mus musculus GN=Magoh PE=2 SV=1 | MOUSE | 2 |
| sp\|Q8BH43\|WASF2_MOUSE | Wiskott-Aldrich syndrome protein family member 2 OS=Mus musculus GN=Wasf2 PE=1 SV=1 | MOUSE | 2 |
| sp\|Q9CXU9\|EIF1B_MOUSE | Eukaryotic translation initiation factor 1b OS=Mus musculus GN=Eif1b PE=1 SV=2 | MOUSE | 2 |
| sp\|P48024\|EIF1_MOUSE | Eukaryotic translation initiation factor 1 OS=Mus musculus GN=Eif1 PE=1 SV=2 | MOUSE | 2 |
| sp\|Q9EQ06\|DHB11_MOUSE | Estradiol 17-beta-dehydrogenase 11 OS=Mus musculus GN=Hsd17b11 PE=1 SV=1 | MOUSE | 2 |
| sp\|P28653\|PGS1_MOUSE | Biglycan OS=Mus musculus GN=Bgn PE=1 SV=1 | MOUSE | 2 |
| sp\|Q8K0E8\|FIBB_MOUSE | Fibrinogen beta chain OS=Mus musculus GN=Fgb PE=1 SV=1 | MOUSE | 2 |
| sp\|P01631\|KV2A7_MOUSE | Ig kappa chain V-II region 26-10 OS=Mus musculus PE=1 SV=1 | MOUSE | 2 |
| sp\|P30355\|AL5AP_MOUSE | Arachidonate 5-lipoxygenase-activating protein OS=Mus musculus GN=Alox5ap PE=1 SV=2 | MOUSE | 2 |
| sp\|P07091\|S10A4_MOUSE | Protein S100-A4 OS=Mus musculus GN=S100a4 PE=1 SV=1 | MOUSE | 2 |
| sp\|P24668\|MPRD_MOUSE | Cation-dependent mannose-6-phosphate receptor OS=Mus musculus GN=M6pr PE=1 SV=1 | MOUSE | 2 |
| sp\|Q9JMH6\|TRXR1_MOUSE | Thioredoxin reductase 1, cytoplasmic OS=Mus musculus GN=Txnrd1 PE=1 SV=3 | MOUSE | 2 |
| sp\|P57780\|ACTN4_MOUSE | Alpha-actinin-4 OS=Mus musculus GN=Actn4 PE=1 SV=1 | MOUSE | 2 |
| sp\|O88990\|ACTN3_MOUSE | Alpha-actinin-3 OS=Mus musculus GN=Actn3 PE=2 SV=1 | MOUSE | 2 |
| sp\|Q7TPR4\|ACTN1_MOUSE | Alpha-actinin-1 OS=Mus musculus GN=Actn1 PE=1 SV=1 | MOUSE | 2 |
| sp\|Q8JZQ9\|EIF3B_MOUSE | Eukaryotic translation initiation factor 3 subunit B OS=Mus musculus GN=Eif3b PE=1 SV=1 | MOUSE | 2 |
| sp\|O88531\|PPT1_MOUSE | Palmitoyl-protein thioesterase 1 OS=Mus musculus GN=Ppt1 PE=1 SV=2 | MOUSE | 2 |
| sp\|Q9Z0U1\|ZO2_MOUSE | Tight junction protein ZO-2 OS=Mus musculus GN=Tjp2 PE=1 SV=2 | MOUSE | 2 |
| sp\|P46061\|RAGP1_MOUSE | Ran GTPase-activating protein 1 OS=Mus musculus GN=Rangap1 PE=1 SV=2 | MOUSE | 2 |
| sp\|P14152\|MDHC_MOUSE | Malate dehydrogenase, cytoplasmic OS=Mus musculus GN=Mdh1 PE=1 SV=3 | MOUSE | 2 |
| sp\|P35278\|RAB5C_MOUSE | Ras-related protein Rab-5C OS=Mus musculus GN=Rab5c PE=1 SV=2 | MOUSE | 2 |
| sp\|Q9D1D4\|TMEDA_MOUSE | Transmembrane emp24 domain-containing protein 10 OS=Mus musculus GN=Tmed10 PE=1 S | MOUSE | 2 |
| sp\|P42208\|SEPT2_MOUSE | Septin-2 OS=Mus musculus GN=Sept2 PE=1 SV=2 | MOUSE | 2 |
| sp\|Q9D0J8\|PTMS_MOUSE | Parathymosin OS=Mus musculus GN=Ptms PE=1 SV=3 | MOUSE | 2 |
| sp\|Q9EQP2\|EHD4_MOUSE | EH domain-containing protein 4 OS=Mus musculus GN=Ehd4 PE=1 SV=1 | MOUSE | 2 |
| sp\|Q8R143\|PTTG_MOUSE | Pituitary tumor-transforming gene 1 protein-interacting protein OS=Mus musculus GN=Pttg1ip | MOUSE | 2 |
| sp\|O08709\|PRDX6_MOUSE | Peroxiredoxin-6 OS=Mus musculus GN=Prdx6 PE=1 SV=3 | MOUSE | 2 |
| sp\|P12246\|SAMP_MOUSE | Serum amyloid P-component OS=Mus musculus GN=Apcs PE=1 SV=2 | MOUSE | 2 |
| sp\|P34022\|RANG_MOUSE | Ran-specific GTPase-activating protein OS=Mus musculus GN=Ranbp1 PE=1 SV=2 | MOUSE | 2 |
| sp\|Q62093\|SRSF2_MOUSE | Serine/arginine-rich splicing factor 2 OS=Mus musculus GN=Srsf2 PE=1 SV=4 | MOUSE | 2 |
| sp\|Q61805\|LBP_MOUSE | Lipopolysaccharide-binding protein OS=Mus musculus GN=Lbp PE=1 SV=2 | MOUSE | 2 |
| sp\|P56565\|S10A1_MOUSE | Protein S100-A1 OS=Mus musculus GN=S100a1 PE=1 SV=2 | MOUSE | 2 |
| sp\|P00405\|COX2_MOUSE | Cytochrome c oxidase subunit 2 OS=Mus musculus GN=Mtco2 PE=1 SV=1 | MOUSE | 2 |
| sp\|Q9WVL7\|CXL15_MOUSE | C-X-C motif chemokine 15 OS=Mus musculus GN=Cxcl15 PE=1 SV=1 | MOUSE | 2 |
| sp\|Q9CWK8\|SNX2_MOUSE | Sorting nexin-2 OS=Mus musculus GN=Snx2 PE=1 SV=2 | MOUSE | 2 |
| sp\|Q9WV80\|SNX1_MOUSE | Sorting nexin-1 OS=Mus musculus GN=Snx1 PE=1 SV=1 | MOUSE | 2 |
| sp\|P21841\|PSPC_MOUSE | Pulmonary surfactant-associated protein C OS=Mus musculus GN=Sftpc PE=1 SV=1 | MOUSE | 2 |
| sp\|O08663\|MAP2_MOUSE | Methionine aminopeptidase 2 OS=Mus musculus GN=Metap2 PE=1 SV=1 | MOUSE | 2 |
| sp\|P61358\|RL27_MOUSE | 60S ribosomal protein L27 OS=Mus musculus GN=Rpl27 PE=1 SV=2 | MOUSE | 2 |
| sp\|P39447\|ZO1_MOUSE | Tight junction protein ZO-1 OS=Mus musculus GN=Tjp1 PE=1 SV=2 | MOUSE | 2 |
| sp\|O55022\|PGRC1_MOUSE | Membrane-associated progesterone receptor component 1 OS=Mus musculus GN=Pgrmc1 PE | MOUSE | 2 |
| sp\|P12265\|BGLR_MOUSE | Beta-glucuronidase OS=Mus musculus GN=Gusb PE=1 SV=2 | MOUSE | 2 |
| sp\|Q9CZM2\|RL15_MOUSE | 60S ribosomal protein L15 OS=Mus musculus GN=Rpl15 PE=2 SV=4 | MOUSE | 2 |
| sp\|Q99KH8\|STK24_MOUSE | Serine/threonine-protein kinase 24 OS=Mus musculus GN=Stk24 PE=1 SV=1 | MOUSE | 2 |
| sp\|Q99JT2\|STK26_MOUSE | Serine/threonine-protein kinase 26 OS=Mus musculus GN=Stk26 PE=1 SV=1 | MOUSE | 2 |
| sp\|P43276\|H15_MOUSE | Histone H1.5 OS=Mus musculus GN=Hist1h1b PE=1 SV=2 | MOUSE | 2 |

| | | | |
|---|---|---|---|
| sp\|Q06335\|APLP2_MOUSE | Amyloid-like protein 2 OS=Mus musculus GN=Aplp2 PE=1 SV=4 | MOUSE | 2 |
| sp\|Q6P4T2\|U520_MOUSE | U5 small nuclear ribonucleoprotein 200 kDa helicase OS=Mus musculus GN=Snrnp200 PE=1 | MOUSE | 2 |
| sp\|P10711\|TCEA1_MOUSE | Transcription elongation factor A protein 1 OS=Mus musculus GN=Tcea1 PE=1 SV=2 | MOUSE | 2 |
| sp\|Q9CQQ7\|AT5F1_MOUSE | ATP synthase F(0) complex subunit B1, mitochondrial OS=Mus musculus GN=Atp5f1 PE=1 SV | MOUSE | 2 |
| sp\|P49312\|ROA1_MOUSE | Heterogeneous nuclear ribonucleoprotein A1 OS=Mus musculus GN=Hnrnpa1 PE=1 SV=2 | MOUSE | 2 |
| sp\|Q9WTY4\|AQP5_MOUSE | Aquaporin-5 OS=Mus musculus GN=Aqp5 PE=1 SV=1 | MOUSE | 2 |
| sp\|O35744\|CHIL3_MOUSE | Chitinase-like protein 3 OS=Mus musculus GN=Chil3 PE=1 SV=2 | MOUSE | 2 |
| sp\|Q5FW60\|MUP20_MOUSE | Major urinary protein 20 OS=Mus musculus GN=Mup20 PE=1 SV=1 | MOUSE | 2 |
| sp\|P19001\|K1C19_MOUSE | Keratin, type I cytoskeletal 19 OS=Mus musculus GN=Krt19 PE=1 SV=1 | MOUSE | 2 |
| sp\|Q6IFX2\|K1C42_MOUSE | Keratin, type I cytoskeletal 42 OS=Mus musculus GN=Krt42 PE=1 SV=1 | MOUSE | 2 |
| sp\|P40240\|CD9_MOUSE | CD9 antigen OS=Mus musculus GN=Cd9 PE=1 SV=2 | MOUSE | 2 |
| sp\|Q8BG15\|CTSL2_MOUSE | CTD small phosphatase-like protein 2 OS=Mus musculus GN=Ctdspl2 PE=1 SV=1 | MOUSE | 2 |
